# Modelling Immune Dynamics in Locally Advanced MSI-H/dMMR Colorectal Cancer with Neoadjuvant Pembrolizumab Treatment: From Differential Equations to an Agent-Based Framework

**DOI:** 10.1101/2025.08.29.672046

**Authors:** Georgio Hawi, Peter Kim, Peter P. Lee

## Abstract

Colorectal cancer (CRC) is the third most common malignancy worldwide, and accounts for approximately 10% of all cancers and an estimated 850,000 deaths annually. Within CRC, MSI-H/dMMR tumours are highly immunogenic due to their high mutational burden and neoantigen load, yet can evade immunosurveillance via PD-1/PD-L1-mediated signalling. Pembrolizumab, an anti-PD-1 antibody approved for unresectable or metastatic MSI-H/dMMR CRC, is emerging as a promising neoadjuvant option in the locally advanced setting, inducing rapid, deep and durable immune responses. In this work, we construct a minimal model of neoadjuvant pembrolizumab therapy in locally advanced MSI-H/dMMR CRC (laMCRC) using ordinary differential equations (ODEs), providing a highly extensible model that captures the main immune dynamics involved. On the other hand, agent-based models (ABMs) naturally capture stochasticity, interactions at an individual level, and discrete events that lie beyond the scope of differential-equation formulations. As such, we also convert our ODE model, with parameters calibrated to experimental data, to an ABM, preserving its dynamics while providing a flexible platform for future mechanistic investigation and modelling.

## 1 Introduction

Colorectal cancer (CRC)—the world’s third most common malignancy, accounting for approximately 10% of all cancers [1] and more than 1.85 million cases with 850,000 deaths annually [2]—remains a major burden, with the American Cancer Society projecting 154,270 new CRC diagnoses and 52,900 deaths in the United States due to CRC in 2025 [3]. A key phenotype within CRC is that of microsatellite instability–high/mismatch repair–deficient (MSI-H/dMMR) disease, characterised by a highly immunogenic tumour microenvironment (TME) driven by markedly elevated somatic mutation rates, tumour mutational burden (TMB), and neoantigen load [4–6]. Although approximately 20% of stage II and 12% of stage III CRC tumours are MSI-H/dMMR [7, 8], patients with this hypermutant phenotype respond poorly to conventional 5-fluorouracil–based chemotherapy [9, 10] yet show striking responsiveness to immune checkpoint inhibitors (ICIs) [11].

Immune checkpoints, such as programmed cell death-1 (PD-1), a cell membrane receptor that is expressed on a variety of cell types, including activated T cells, activated NK cells and monocytes, normally downregulate immune responses after antigen activation [12]. PD-1 has been extensively studied in cancer, including MSI-H/dMMR CRC [13, 14]; engagement by its ligands PD-L1 and PD-L2 inhibits effector T-cell activity, suppresses pro-inflammatory cytokine secretion, and promotes expansion of regulatory T cells (Tregs) [15, 16]. Tumours further exploit this pathway by expressing PD-L1 themselves, thereby impairing cytotoxic T-lymphocyte (CTL) and natural killer (NK) cell activity [17]. However, blocking PD-1/PD-L1 complex formation reinvigorates effector T cells, enhances anti-tumour immunity, and improves clinical outcomes [18, 19]. Pembrolizumab, a fully human IgG4 anti–PD-1 antibody, demonstrated significant efficacy in metastatic MSI-H/dMMR CRC in the phase III KEYNOTE-177 trial (NCT02563002) [20], leading the FDA to approve pembrolizumab for first-line treatment of unresectable or metastatic MSI-H/dMMR CRC on June 29, 2020 [21].

Furthermore, in recent years, interest in neoadjuvant pembrolizumab for the treatment of high-risk stage II and stage III MSI-H/dMMR CRC has surged [22]. In the phase II NEOPRISM-CRC study (NCT05197322), 31 patients with high TMB and high-risk stage II–III MSI-H/dMMR CRC received three cycles of pembrolizumab followed by surgery 4–6 weeks later; pathologic complete response (pCR) was achieved in 17 patients, and no recurrences were observed at a median 6-month follow-up [23]. Another phase II trial, NCT04082572, evaluated the efficacy of neoadjuvant pembrolizumab in localised MSI-H/dMMR solid tumours [24]. In the locally advanced MSI-H/dMMR CRC (laMCRC) cohort, 27 patients received pembrolizumab 200 mg IV every 3 weeks either for up to 16 cycles or eight cycles followed by resection. Overall, 21 patients exhibited pCR, and among the 14 patients in the resection group, 11 achieved pCRs. At a median 9.5-month follow-up, only two recurrences/progressions were observed, and at a median 3-year follow-up, no late progression events occurred, demonstrating the efficacious and durable response of pembrolizumab in localised dMMR/MSI-H tumours [25].

Mathematical models provide a powerful framework for analysing the immunobiology underpinning diseases, and, when mechanistic, for investigating the dynamics of key biological components to improve theoretical understanding. They also enable the optimisation of treatment regimens whilst avoiding the substantial time and financial costs of human clinical trials. Numerous immunobiological models of CRC exist [26–29], and ICI therapy has been modelled extensively in other cancers [30– 34]. Nonetheless, there are a multitude of limitations and drawbacks to these models of CRC and ICI therapy, as detailed in [35]. In this work, we focus on our comprehensive model of neoadjuvant pembrolizumab therapy in laMCRC [35], which, to the authors’ knowledge, is the only deterministic immunobiological model of ICI therapy in CRC and addresses many of these drawbacks.

However, to effectively model complex biological systems, one must master the balance between model complexity and model accuracy. Traditional mechanistic models often incorporate many processes, leading to numerous state variables, inputs, and parameters [36]. As such, it is ideal to reduce models such that their key dynamics are preserved, while pruning parameters and inputs with minimal influence on the outputs. The full model in [35] is large, comprising complex delay integro-differential equations with many components and interactions. In this work, we construct a minimal model of pembrolizumab therapy in laMCRC using ordinary differential equations (ODEs), avoiding these complexities, to reveal the core immune dynamics and interactions, and accurately replicate the state-variable trajectories of the full model.

Agent-based models (ABMs) are well-suited to model complex biological systems, such as cancer, due to their ability to explicitly represent individual entities and their interactions, incorporate spatial structure, and capture phenotypic heterogeneity [37, 38]. Their inherent stochasticity enables the estimation of probabilities and predictive intervals for treatment response—features which are impossible to be predicted by differential equations due to their deterministic, mean-field nature [39]. As such, ABMs are being increasingly used to model cancer dynamics, with Kather et al. developing patient-informed ABMs have been used to study the response of combined immunotherapy and stroma-targeting therapies in CRC [40], as well as enable in-silico screening of combination immunotherapies [41]. For ICIs broadly, multiscale ABMs of PD-1/PD-L1 interactions have reproduced spatiotemporal tumour-CTL dynamics and linked response to ICIs to antigenicity and TMB [42].

Nonetheless, because ABMs typically encode many interactions and processes, their stochastic and computationally expensive simulations make statistical calibration of model parameters to experimental data difficult. In this work, we outline a practical workflow for converting ODE models to ABMs and apply this to our minimal ODE model of neoadjuvant pembrolizumab therapy in laMCRC, allowing for accurate and efficient parameter estimation of the ABM. We finally compare ABM trajectories of model components with those from the minimal ODE model and the full model from [35], verifying consistency and faithful reproduction of all dynamics, thereby laying a foundation for future ABMs of laMCRC and other cancers.

## 2 A Minimal Model of Locally Advanced MSI-H/dMMR CRC

The derivation of many equations in the minimal model closely follows [35], but is repeated here for the sake of completeness.

### 2.1 Model Variables

The variables and their units in the minimal model are shown in Table 1.

**Table 1.**
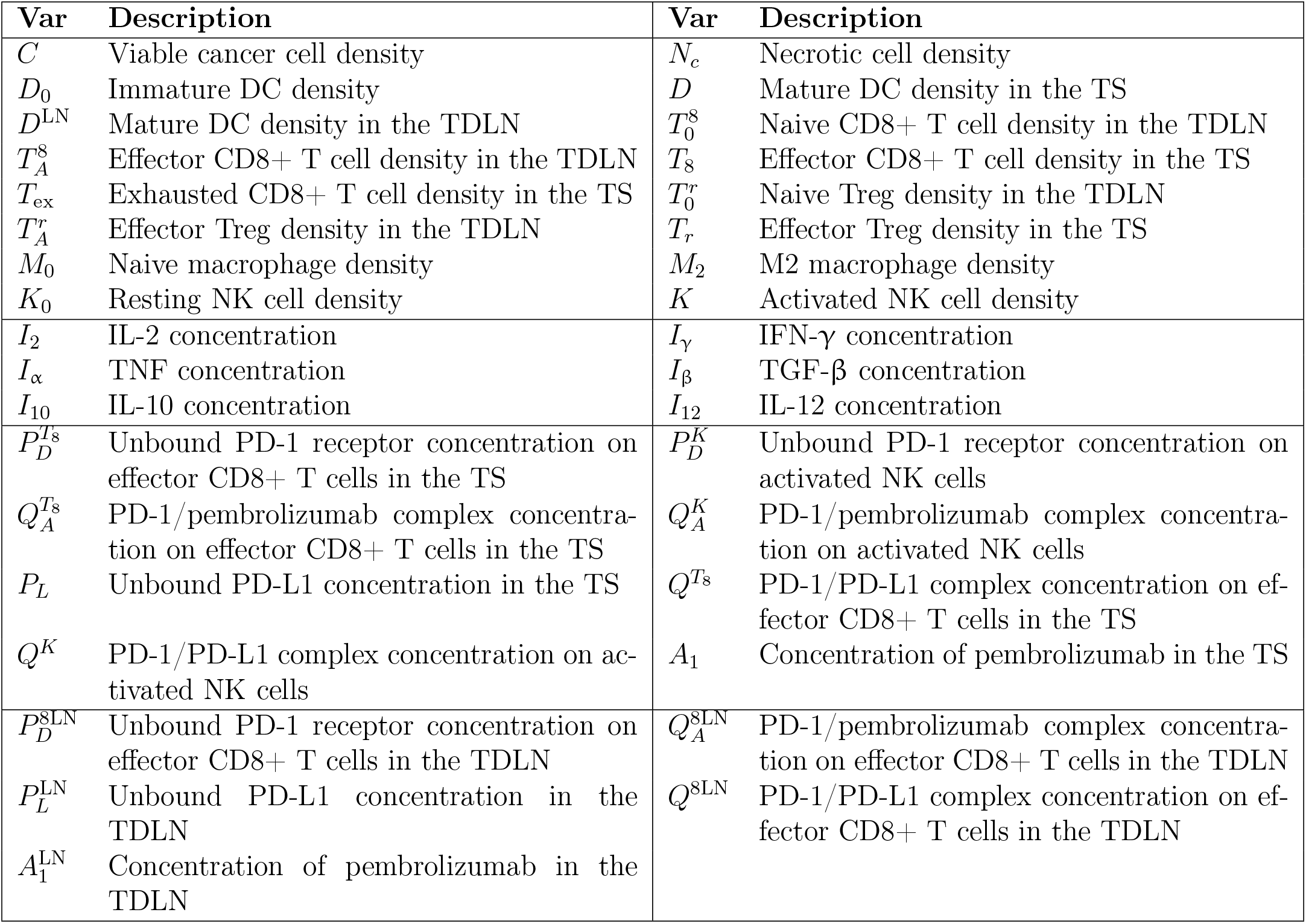
Variables used in the model. Quantities in the top box are in units of cell*/*cm^3^, quantities in the second box are in units of g*/*cm^3^, and all other quantities are in units of molec*/*cm^3^. All quantities pertain to the tumour site.

| Var | Description | Var | Description |
| --- | --- | --- | --- |
| $C$ | Viable cancer cell density | $N_c$ | Necrotic cell density |
| $D_0$ | Immature DC density | $D$ | Mature DC density in the TS |
| $D^{\text{LN}}$ | Mature DC density in the TDLN | $T_0^8$ | Naive CD8+ T cell density in the TDLN |
| $T_A^8$ | Effector CD8+ T cell density in the TDLN | $T_8$ | Effector CD8+ T cell density in the TS |
| $T_{\text{ex}}$ | Exhausted CD8+ T cell density in the TS | $T_0^r$ | Naive Treg density in the TDLN |
| $T_A^r$ | Effector Treg density in the TDLN | $T_r$ | Effector Treg density in the TS |
| $M_0$ | Naive macrophage density | $M_2$ | M2 macrophage density |
| $K_0$ | Resting NK cell density | $K$ | Activated NK cell density |
| $I_2$ | IL-2 concentration | $I_\gamma$ | IFN-γ concentration |
| $I_\alpha$ | TNF concentration | $I_\beta$ | TGF-β concentration |
| $I_{10}$ | IL-10 concentration | $I_{12}$ | IL-12 concentration |
| $P_D^{T_8}$ | Unbound PD-1 receptor concentration on effector CD8+ T cells in the TS | $P_D^K$ | Unbound PD-1 receptor concentration on activated NK cells |
| $Q_A^{T_8}$ | PD-1/pembrolizumab complex concentration on effector CD8+ T cells in the TS | $Q_A^K$ | PD-1/pembrolizumab complex concentration on activated NK cells |
| $P_L$ | Unbound PD-L1 concentration in the TS | $Q^{T_8}$ | PD-1/PD-L1 complex concentration on effector CD8+ T cells in the TS |
| $Q^K$ | PD-1/PD-L1 complex concentration on activated NK cells | $A_1$ | Concentration of pembrolizumab in the TS |
| $P_D^{8\text{LN}}$ | Unbound PD-1 receptor concentration on effector CD8+ T cells in the TDLN | $Q_A^{8\text{LN}}$ | PD-1/pembrolizumab complex concentration on effector CD8+ T cells in the TDLN |
| $P_L^{\text{LN}}$ | Unbound PD-L1 concentration in the TDLN | $Q^{8\text{LN}}$ | PD-1/PD-L1 complex concentration on effector CD8+ T cells in the TDLN |
| $A_1^{\text{LN}}$ | Concentration of pembrolizumab in the TDLN | | |

### 2.2 Model Diagrams

Diagrams encompassing the interactions of model components in the TS and TDLN are shown in Figure 1 and Figure 2, respectively.

**Figure 1.**
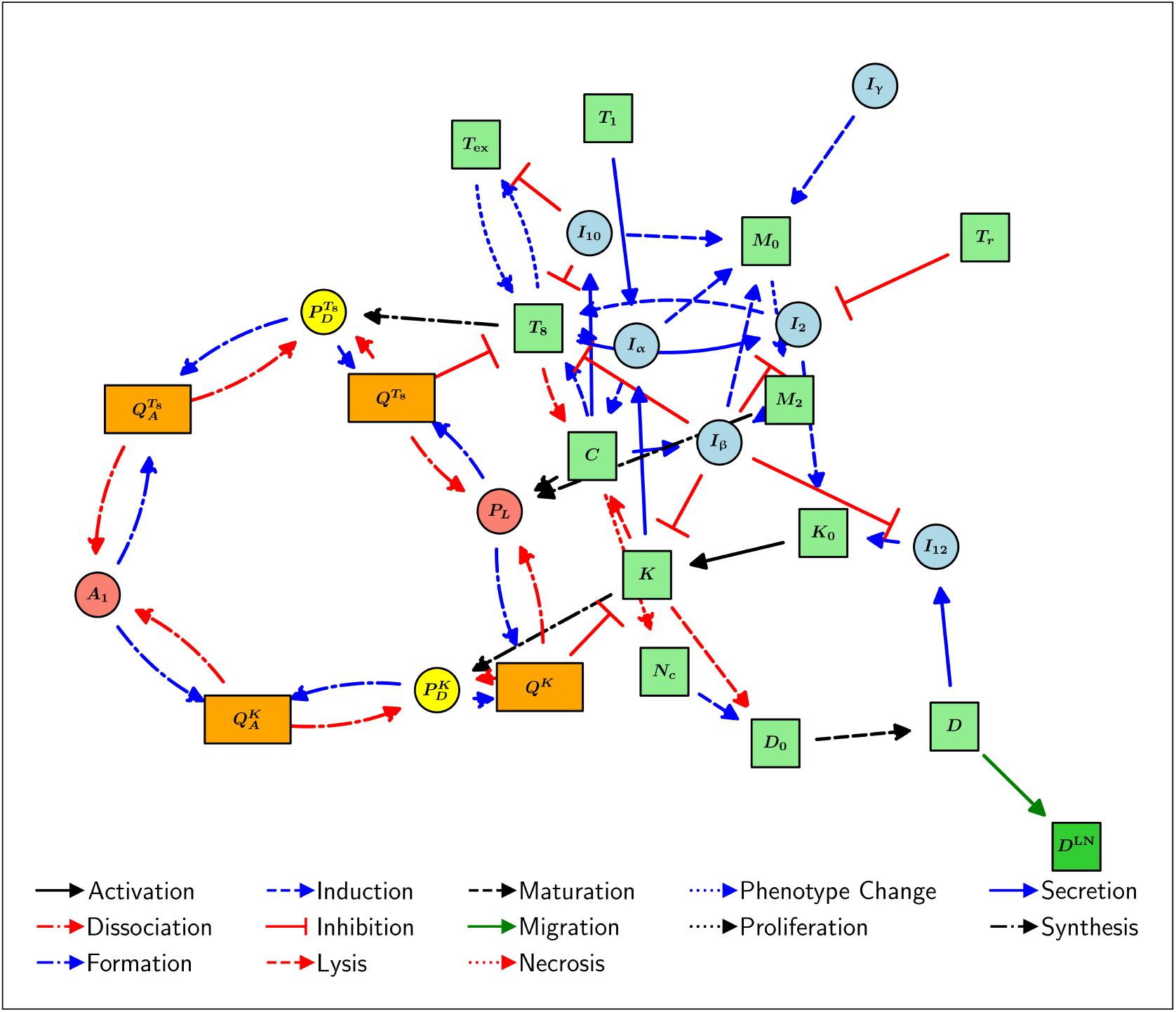
Schematic diagram of the interactions of model components in the TS.

**Figure 2.**
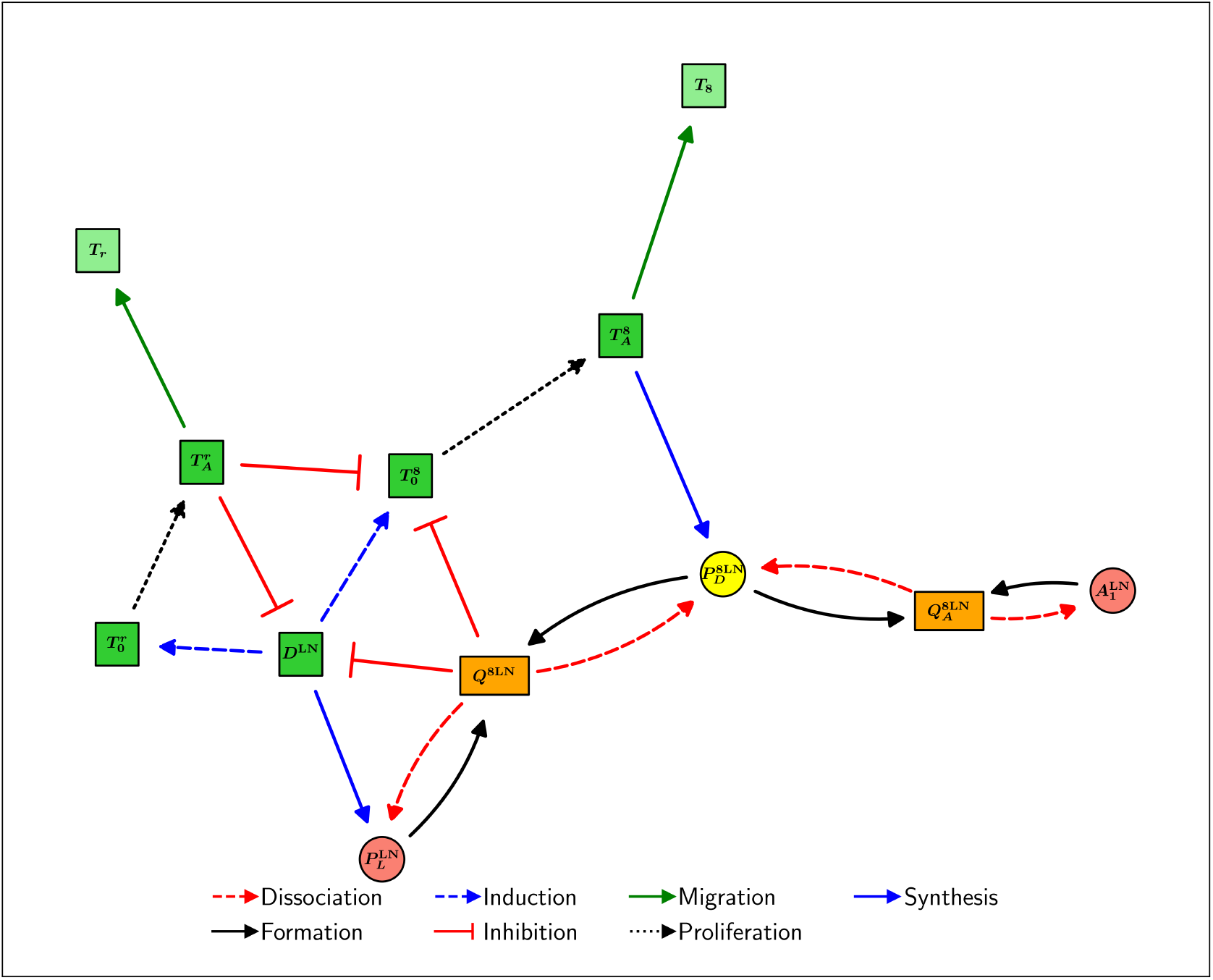
Schematic diagram of the interactions of model components in the TDLN.

### 2.3 Model Assumptions

For simplicity, we ignore spatial effects in the model, ignoring the effects of diffusion, advection, and chemotaxis by all species. We assume the system has one compartment, being the TS, located in the colon or rectum. This is a simplification since locally advanced CRC typically involves multiple tumour-draining lymph nodes [43]; however, for simplicity, we focus on the sentinel node and refer to it as the TDLN for the purposes of the model. We assume that cytokines in the TS are produced only by effector or activated cells and that DAMPs in the TS are only produced by necrotic cancer cells. We assume that all mature DCs considered in the TDLN are cancer-antigen-bearing and that all T cells considered in the TS are primed with cancer antigens. Furthermore, we assume that all activated T cells considered in the TDLN are activated with cancer antigens and that T cell proliferation/division follows a deterministic program. We ignore CD8+ memory T cells. We also assume that all Tregs are natural Tregs (nTregs), ignoring induced Tregs (iTregs). We also assume that the duration of pembrolizumab infusion is negligible compared to the timescale of the model. Therefore, we treat their infusions as an intravenous bolus so that drug absorption occurs immediately after infusion. Finally, we assume a constant solution history, where the history for each species is set to its respective initial condition.

We assume that all species, *X*_*i*_, degrade/die at a rate proportional to their concentration, with decay constant 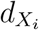. We assume that the rate of activation/polarisation of a species *X*_*i*_ by a species *X*_*j*_ follows the Michaelis-Menten kinetic law 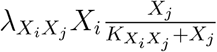, for rate constant 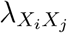, and half-saturation constant 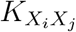. Similarly, we model the rate of inhibition of a species *X*_*i*_ by a species *X*_*j*_ using a term with form 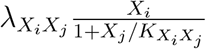 for rate constant 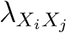, and inhibition constant 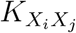. Production of *X*_*i*_ by *X*_*j*_ is modelled using mass-action kinetics unless otherwise specified, so that the rate at which *X*_*i*_ is formed is given by 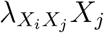 for some positive constant 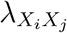. Finally, we assume that the rate of lysis of *X*_*i*_ by *X*_*j*_ follows mass-action kinetics in the case where *X*_*j*_ is a cell and follows Michaelis-Menten kinetics in the case where *X*_*j*_ is a cytokine.

### 2.4 Model Equations

#### 2.4.1 Equations for Cancer Cells (*C* and *N*_*c*_)

Viable cancer cells are killed by effector CD8+ T cells [44] and activated NK cells [45] through direct contact, whilst TNF indirectly eliminates cancer cells via activating cell death pathways [46–48]. In particular, TNF induces the necroptosis, programmed necrotic cell death, of cancer cells [47, 49]. We note that TGF-β and the PD-1/PD-L1 complex inhibit cancer cell lysis by CD8+ T cells [50–52], and that TGF-β and PD-1/PD-L1 have been shown to inhibit NK cell cytotoxicity [15, 53–57]. We assume that viable cancer cells grow logistically, as is done in many CRC models [26, 27, 29], due to space and resource competition in the TME. Combining these, we have

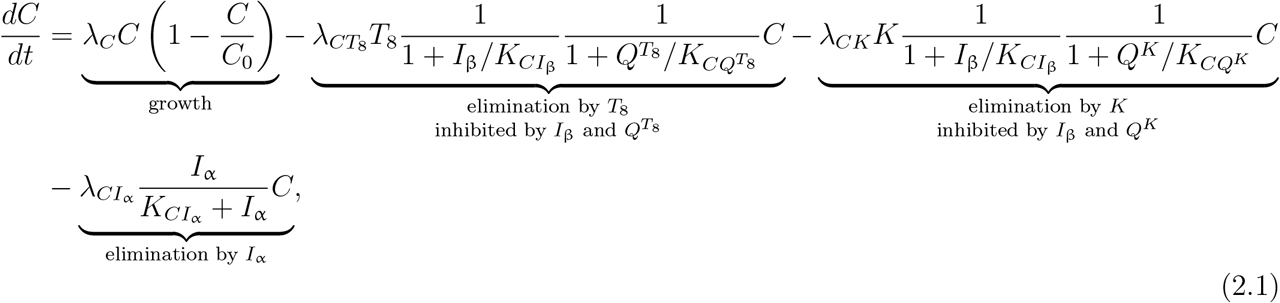

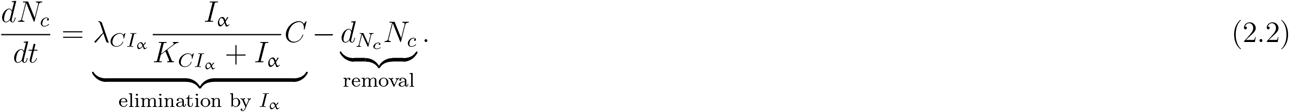

#### 2.4.2 Equations for Immature and Mature DCs in the TS (*D*_0_ and *D*)

Immature DCs are stimulated to mature via DAMPs, such as HMGB1 [58], whose concentration is assumed to be proportional to that of necrotic cancer cells. In addition, activated NK cells have been shown to efficiently kill immature DCs but not mature DCs; however, this is inhibited by TGF-β [59–61]. We also need to consider that some mature DCs migrate into the T cell zone of the TDLN and stimulate naive T cells, causing them to be activated [62, 63]. Assuming that immature DCs are supplied at a rate 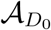, we have that

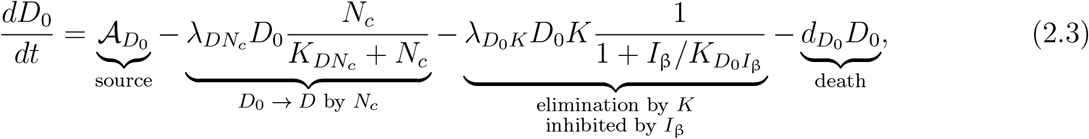

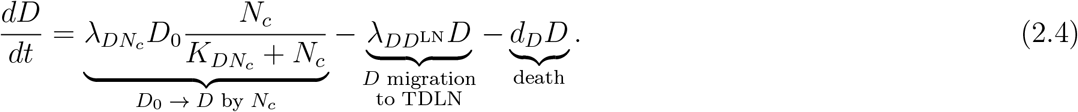

#### 2.4.3 Equation for Mature DCs in the TDLN (*D*^LN^)

We assume a fixed DC migration time of *τ*_*m*_ and also assume that only exp (−*d*_*D*_*τ*_*m*_) of the mature DCs that leave the TS survive migration. We also assume that the concentration of mature DCs in the TDLN does not change significantly during migration. Taking into account the volume change between the TS and the TDLN, we have that

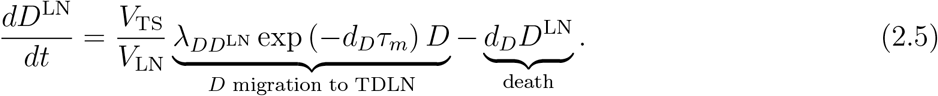

#### 2.4.4 Equation for Naive CD8+ T Cells in the TDLN 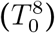

We assume that naive CD8+ T cells come into the TDLN at a constant rate and that they have not undergone cell division, nor will they until their activation, which we assume to occur instantaneously. For simplicity, we do not consider cytokines in the TDLN, absorbing their influence into 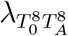. We do, however, explicitly take into account the influence of effector Tregs and the PD-1/PD-L1 complex in the TDLN, which have been shown to inhibit T cell activation via mechanisms including limiting naive T cells from binding to mature DCs [64–72]. Recalling that T cells that have become activated by mature DCs are no longer naive, and taking this all into account, leads to

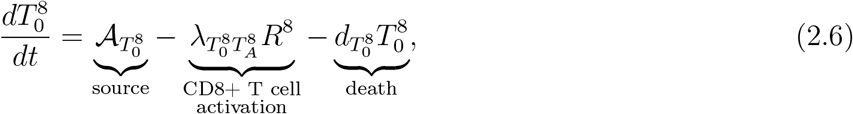

where *R*^8^ is defined as

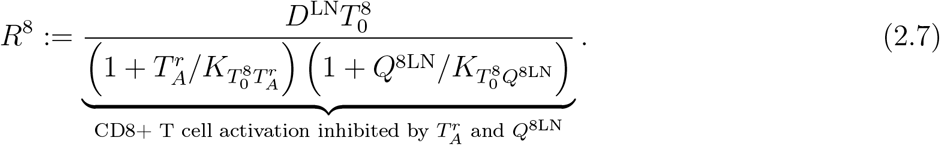

#### 2.4.5 Equation for Effector CD8+ T Cells in the TDLN 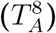

It is known that activated CD8+ T cells undergo clonal expansion in the TDLN and differentiate before they stop proliferating and migrate to the TS [73, 74].

We assume that activated CD8+ T cells proliferate up to 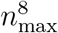 times, upon which they stop dividing. For simplicity, we assume that the death rate of CD8+ T cells that have not completed their division program is equal to 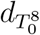, the death rate of naive CD8+ T cells, regardless of the number of cell divisions previously undergone. We also assume that only activated CD8+ T cells that have undergone 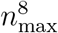 divisions become effector CD8+ T cells, which will leave the TDLN and migrate to the TS. Furthermore, we assume a constant cell cycle time of Δ_8_, except for the first cell division, which has a cycle time of 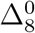. Thus, the duration of the activated CD8+ T cell division program to 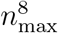 divisions is given by

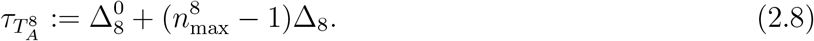

In particular, we must take into account that some T cells will die before the division program is complete, so we must introduce a shrinkage factor of 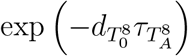. Additionally, we assume that the concentration of components in the TDLN does not change significantly during proliferation. Furthermore, we must also take into account that effector Tregs and the PD-1/PD-L1 complex inhibit CD8+ T cell proliferation throughout the program [65–67, 75, 76]. We must also consider that some of these effector CD8+ T cells will migrate to the TS to perform effector functions. We finally assume that the death rate of CD8+ T cells that have completed their division program is equal to the death rate of CD8+ T cells in the TS. Taking this all into account leads to

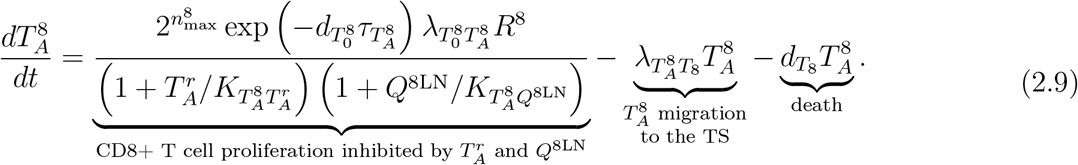

#### 2.4.6 Equations for Effector and Exhausted CD8+ T Cells in the TS (*T*_8_ and *T*_ex_)

We assume that it takes *τ*_*a*_ amount of time for effector CD8+ T cells in the TDLN to migrate to the TS. Like with DCs, we assume that the concentration of effector CD8+ T cells in the TDLN does not change significantly during migration. We must also account for CTL expansion due to IL-2 [77], noting that this proliferation is inhibited by effector Tregs [65–67]. Furthermore, the death of CD8+ T cells is resisted by IL-10 [78, 79].

However, chronic antigen exposure can cause effector CD8+ T cells to enter a state of exhaustion, where they lose their ability to kill cancer cells, and the rate of cytokine secretion significantly decreases [80–82]. We denote this exhausted CD8+ T cell population as *T*_ex_. It has also been shown that pembrolizumab can “reinvigorate” these cells back into the effector state [18, 83]. We model the re-invigoration and exhaustion using Michaelis-Menten terms in *A*_1_ and *C*, respectively. In particular, this has been shown to be more appropriate than simple mass-action kinetics as it accounts for extended antigen exposure [84].

As such, remembering to take the volume change between the TDLN and the TS into account, this implies that

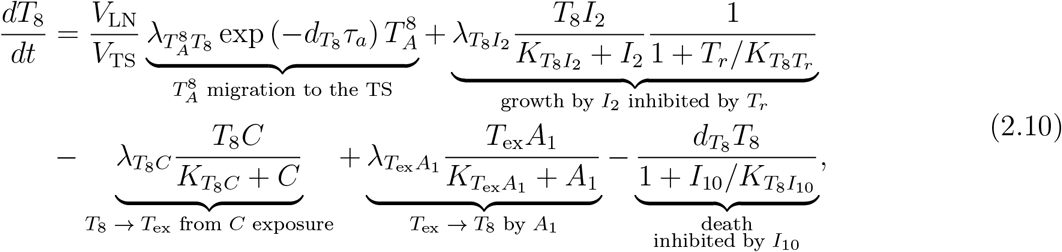

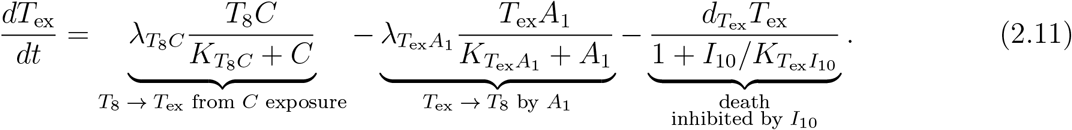

#### 2.4.7 Equation for Naive Tregs in the TDLN 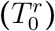

We also consider the concentration of naive Tregs in the TDLN, following the same procedure as for CD8+ T cells. We absorb the influence of cytokines on Treg activation via the kinetic rate constant 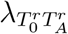. We also take into account that some mature DCs migrate into the TDLN and activate naïve Tregs, causing them to no longer be naive. Assuming that naive Tregs come into the TDLN at a rate 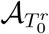, we can write a similar equation to (2.6):

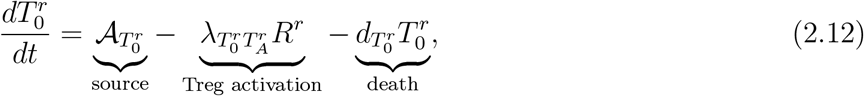

where *R*^*r*^ is defined as

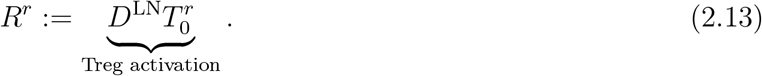

#### 2.4.8 Equation for Effector Tregs in the TDLN 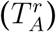

We assume that activated Tregs proliferate up to 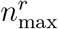 times, upon which they stop dividing and become effector Tregs. As before, we assume that the death rate of Tregs that have not completed their division program is equal to 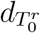, the death rate of naive Tregs. We assume a constant cell cycle time of Δ_*r*_, except for the first cell division, which has a cycle time of 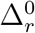. Thus, the duration of the activated Treg division program to 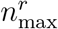 divisions is given by

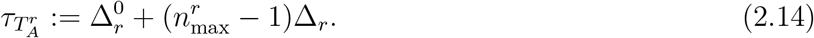

In particular, we must take into account that some T cells will die before the division program is complete, so we must introduce a shrinkage factor of 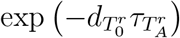. We, again, assume that the concentration of components in the TDLN does not change significantly during proliferation. We also assume that the death rate of effector Tregs in the TDLN is equal to the corresponding degradation rate in the TS. Taking this all into account, and incorporating effector Treg migration to the TS, leads to

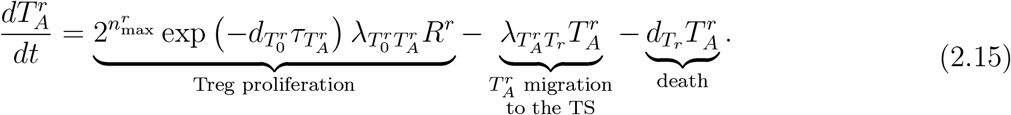

#### 2.4.9 Equation for Effector Tregs in the TS (*T*_*r*_)

Assuming that it also takes *τ*_*a*_ amount of time for Tregs to migrate to the TS, and that the concentration of effector Tregs in the TDLN does not change significantly during migration, we have that

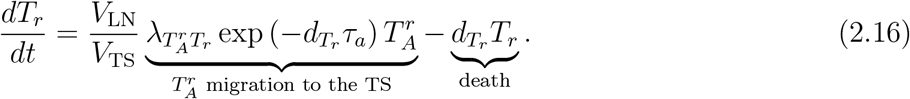

#### 2.4.10 Equations for Naive and M2 Macrophages (*M*_0_ and *M*_2_)

While we only consider naive and anti-inflammatory M2 macrophages, we must also take into account that some naive macrophages polarise to the pro-inflammatory M1 phenotype, whose density we denote as *M*_1_. In particular, IFN-γ and TNF polarise naive macrophages into M1 macrophages [85–88], whilst TGF-β and IL-10 polarise naive macrophages into the M2 phenotype [89–91]. Assuming a production rate 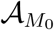 of naive macrophages, we thus have that

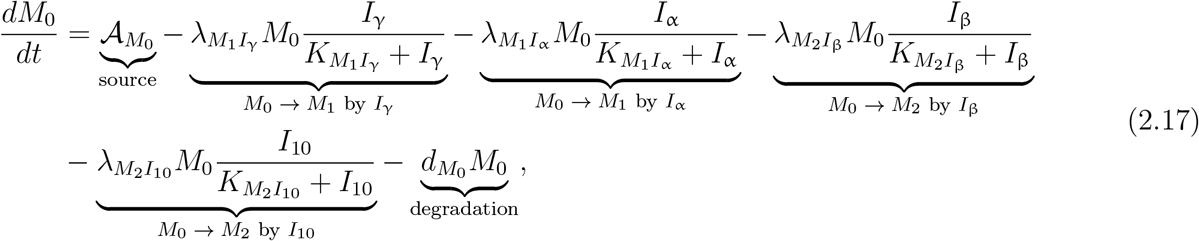

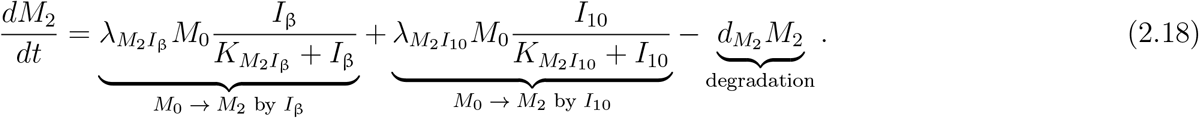

#### 2.4.11 Equations for Resting and Activated NK Cells (*K*_0_ and *K*)

Resting NK cells are activated by IL-2 and IL-12 [92, 93]. However, NK cell activation is inhibited by TGF-β [94]. Thus, assuming a supply rate 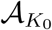 of resting NK cells, we have that

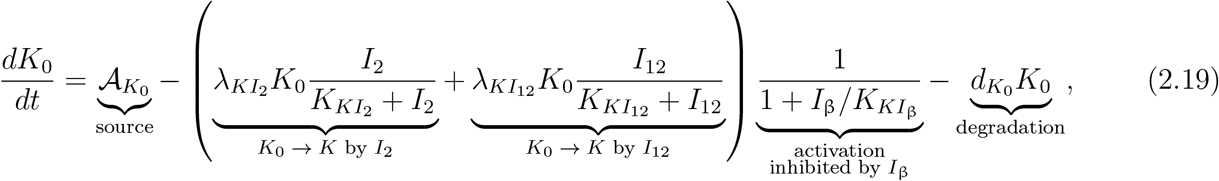

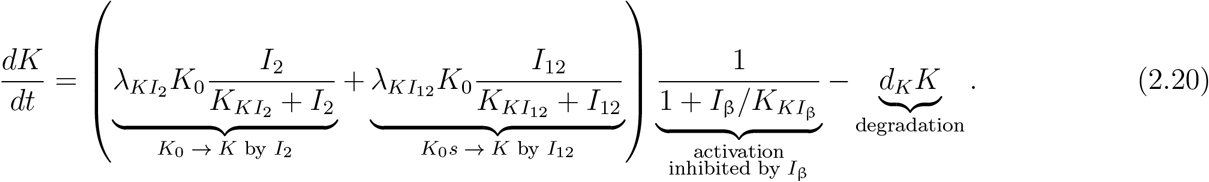

#### 2.4.12 Equation for IL-2 (*I*_2_)

IL-2 is produced by effector CD8+ T cells [95, 96], so that

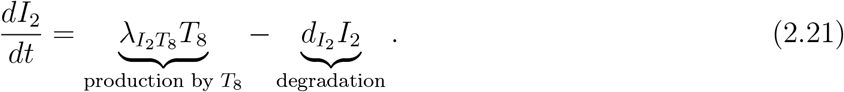

#### 2.4.13 Equation for IFN-γ (*I*_γ_)

IFN-γ is produced by activated NK cells [97] so that

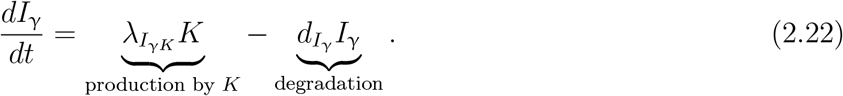

#### 2.4.14 Equation for TNF (*I*_α_)

TNF is produced by effector CD8+ T cells [98, 99] and activated NK cells [100, 101]. Hence,

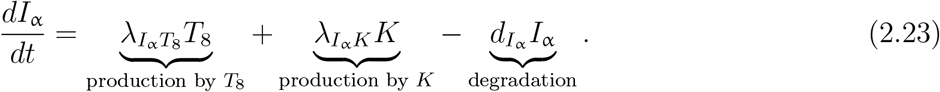

#### 2.4.15 Equation for TGF-β (*I*_β_)

TGF-β is produced by viable cancer cells [102] and M2 macrophages [90, 103]. Thus,

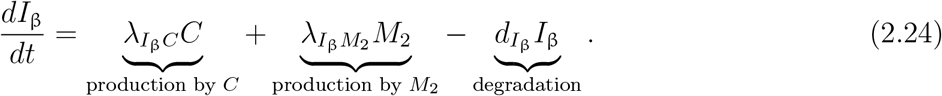

#### 2.4.16 Equation for IL-10 (*I*_10_)

IL-10 is produced by viable cancer cells [104, 105] so that

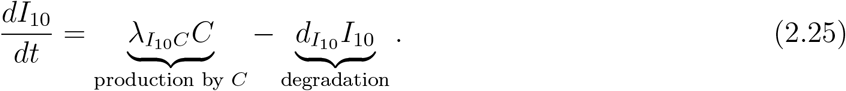

#### 2.4.17 Equation for IL-12 (*I*_12_)

IL-12 is produced by mature DCs [106] so that

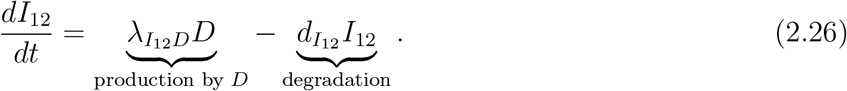

#### 2.4.18 Equations for Unbound PD-1 receptors on Cells in the TS (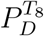 and 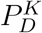)

It is known that PD-1 is expressed on the surface of effector CD8+ T cells [107–109] and activated NK cells [55, 57, 110]. We assume that the rate of PD-1 synthesis is proportional to the concentration of the cell expressing it. However, unbound PD-1 receptors on these PD-1-expressing cells can bind to either pembrolizumab or PD-L1, forming the PD-1/pembrolizumab and PD-1/PD-L1 complexes, respectively, resulting in the depletion of unbound PD-1 molecules [16, 111]. For simplicity, we assume that the formation and dissociation rates of the PD-1/PD-L1 and PD-1/pembrolizumab complexes are invariant of the type of cell expressing PD-1. Considering unbound PD-1 receptors on effector CD8+ T cells in the TS at first, and taking into account the degradation of PD-1 receptors, this motivates the equation for 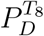 to be

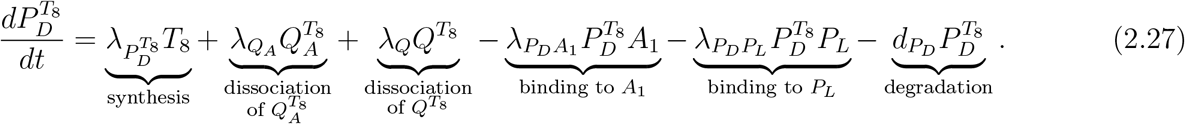

Similarly, we have that

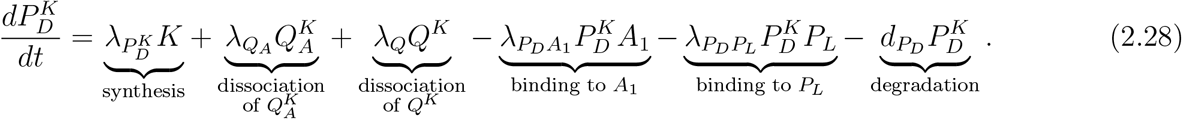

#### 2.4.19 Equations for the PD-1/pembrolizumab Complex on Cells in the TS (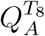 and 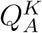)

Pembrolizumab binds to unbound PD-1 on the surfaces of PD-1-expressing cells in a 1:1 ratio [112], forming the PD-1/pembrolizumab complex in a reversible chemical process [113, 114]. We must also account for loss due to the endocytosis and internalisation of the PD-1/pembrolizumab complex from the surface of cells [115, 116]. We assume that the rates of PD-1/pembrolizumab complex internalisation and dissociation are invariant of the type of cell expressing PD-1, so that

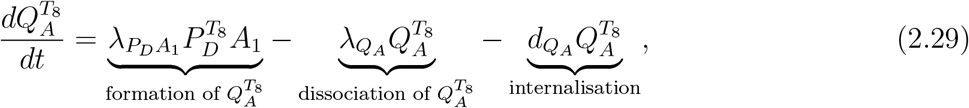

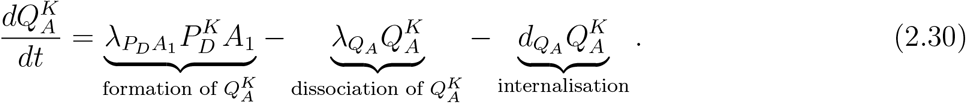

#### 2.4.20 Equation for Pembrolizumab in the TS (*A*_1_)

We assume that pembrolizumab is administered intravenously at times *t*_1_, *t*_2_, …, *t*_*n*_ with doses *ξ*_1_, *ξ*_2_, …, *ξ*_*n*_ respectively, assuming that the duration of infusion is negligible in comparison to the time period of interest. We also account for pembrolizumab depletion due to binding to unbound PD-1, replenishment due to PD-1/pembrolizumab complex dissociation, and elimination of pembrolizumab. It is important to note that the administered dose is not equal to the corresponding change in concentration in the TS. For simplicity, we assume linear pharmacokinetics so that, for some scaling factor *f*_pembro_, we have that

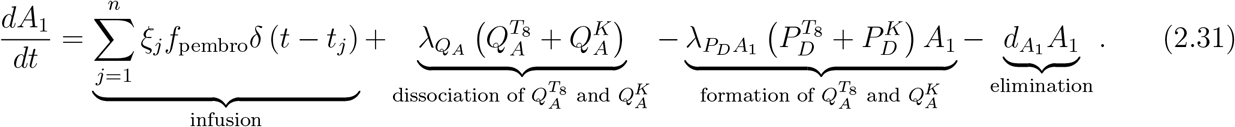

#### 2.4.21 Equation for Unbound PD-L1 in the TS (*P*_*L*_)

We also know that PD-L1 is expressed on the surface of viable cancer cells and M2 macrophages in the TS [117, 118]. We must take into account the synthesis of PD-L1, its depletion due to binding to unbound PD-1, replenishment due to PD-1/PD-L1 complex dissociation, and the degradation of PD-L1. Hence,

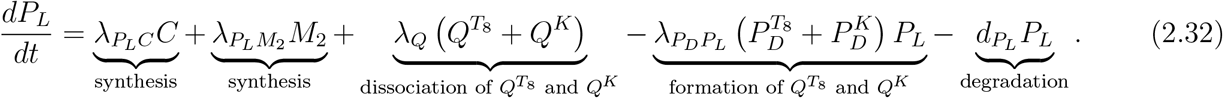

#### 2.4.22 Equations for the PD-1/PD-L1 Complex in the TS (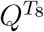 and *Q*^*K*^)

PD-L1 binds to unbound PD-1 receptors on the surfaces of PD-1-expressing cells in a 1:1 ratio [119], forming the PD-1/PD-L1 complex in a reversible chemical process. Considering 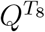 as an example, we can express its formation and dissociation via the reaction 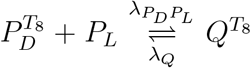. We assume that the degradation is negligible relative to the dissociation, so that

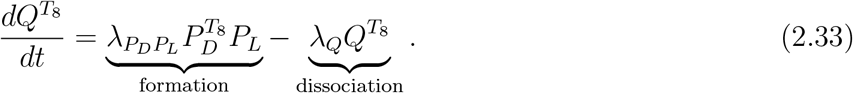

However, the dissociation rate constant of the PD-1/PD-L1 complex is 1.44 s^*−*1^, corresponding to a mean lifetime of less than 1 second [119]. As such, we employ a quasi-steady-state approximation (QSSA) for 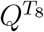, so that 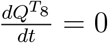, so that

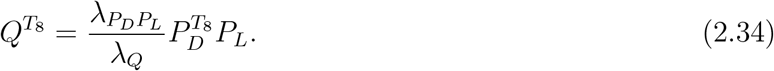

Similarly,

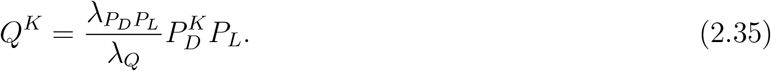

Furthermore, we can simplify (2.27), (2.28), and (2.32) by substituting in (2.34) and (2.35), so that

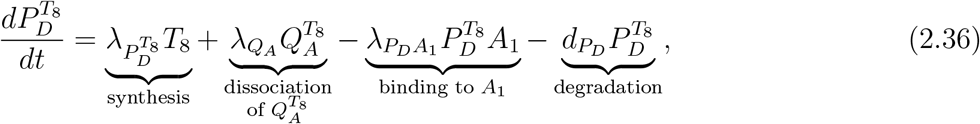

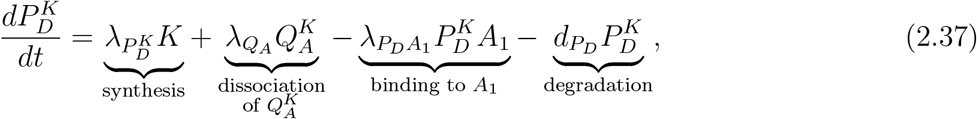

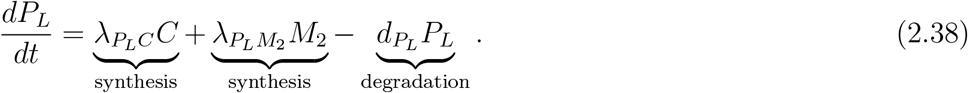

#### 2.4.23 Equation for Unbound PD-1 receptors on Cells in the TDLN 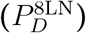

The equation for 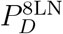 follows identically to that of (2.36). For simplicity, we assume that the formation and dissociation rates of the PD-1/pembrolizumab complex are identical in the TDLN and the TS so that

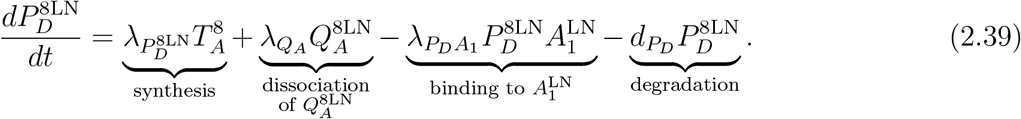

#### 2.4.24 Equation for the PD-1/pembrolizumab Complex on Cells in the TDLN 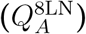

The equation for 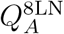 follows identically to that of (2.29). For simplicity, we assume that the rates of PD-1 receptor internalisation are identical in the TDLN and the TS, so that

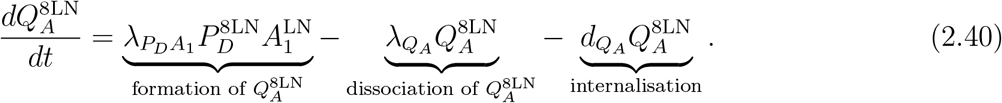

#### 2.4.25 Equation for Pembrolizumab in the TDLN 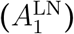

The equation for 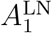 follows identically to that of (2.31) so that

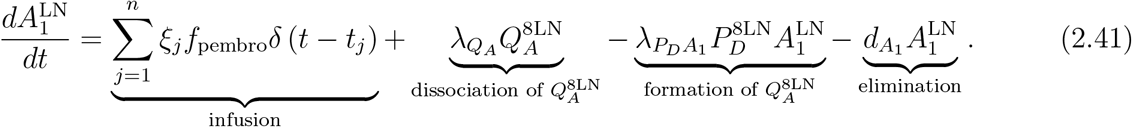

#### 2.4.26 Equation for Unbound PD-L1 in the TDLN 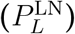

We know that PD-L1 is expressed on the surface of mature DCs in the TDLN [120]. The equation for 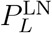 follows identically to (2.38) so that

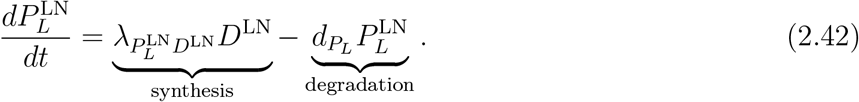

#### 2.4.27 Equations for the PD-1/PD-L1 Complex in the TDLN (*Q*^8LN^)

For simplicity, we assume that the formation and dissociation rates of the PD-1/PD-L1 complex are identical in the TDLN and the TS. The equation for *Q*^8LN^ follows identically from (2.34) so that

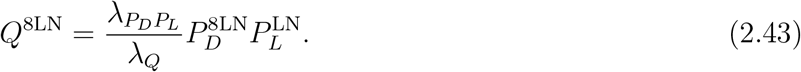

We note that throughout the model, the PD-1/PD-L1 complex appears only within an inhibition constant, making its absolute magnitude less important since it always appears as a ratio. One thing to note is that activated CD8+ T cells also express PD-1 receptors and PD-L1 ligands, and we assume that effector and activated cells express these in equal amounts. However, as discussed in [35], the ratio between effector and activated T cells can be assumed to remain roughly constant. Since the PD-1/PD-L1 complex concentration is linearly proportional to the product of PD-L1 concentration and unbound PD-1 receptor concentration, and PD-1/PD-L1-mediated inhibition of T cell proliferation in the TDLN appears only as a ratio, it is sufficient to consider only PD-1, PD-L1, and PD-1/PD-L1 concentrations on effector cells, as this will be appropriately scaled by the corresponding inhibition constants. Furthermore, this also justifies using the PD-1/PD-L1 complex concentration on effector T cells as a proxy for its concentration on activated T cells that have not yet undergone division, given that their ratio to effector cells remains roughly constant and that PD-1/PD-L1-mediated inhibition of T cell activation in the TDLN appears only as a ratio.

We also note that it is clear, from the structure of the model, that the non-negative orthant is positively forward invariant for all non-negative parameter choices, as the corresponding vector field is essentially non-negative; i.e., for each species *X*_*i*_, if *X*_*i*_ = 0 with all other species non-negative, then 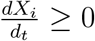 because every negative term in 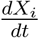 contains *X*_*i*_.

### 2.5 Model Reduction via QSSA

The model parameter values are estimated in Appendix B and are listed in Table B.5.

We observe that the degradation rates of cytokines are, in general, orders of magnitude larger than those of immune and cancer cells. In particular, IL-2, TNF, TGF-β, and IL-12 evolve on a very fast timescale, with degradation rates significantly higher than all other species in the model, causing them to equilibrate much more rapidly. Additionally, the combined death and migration rates of effector CD8+ T cells and Tregs, and the combined polarisation and death rates of naive macrophages, are much higher than the death rates of other model components, and compared with other model components, the PD-1/pembrolizumab complex undergoes dissociation or internalisation more rapidly. As such, we perform a QSSA and reduce the model by setting (2.9), (2.15), (2.17), (2.21) – (2.26), (2.29), (2.30), and (2.40) to 0 and solving for 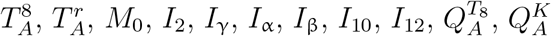, and 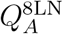 in terms of the other parameters and variables in the model. This minimally affects the system’s evolution after a very short period of transient behaviour [121], and we justify this by observing that, empirically, the deviation in system trajectories remains negligible for nearby parameter choices. Performing the QSSA leads to

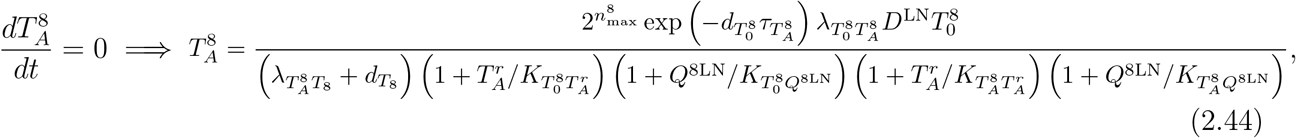

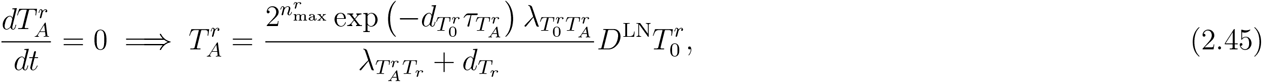

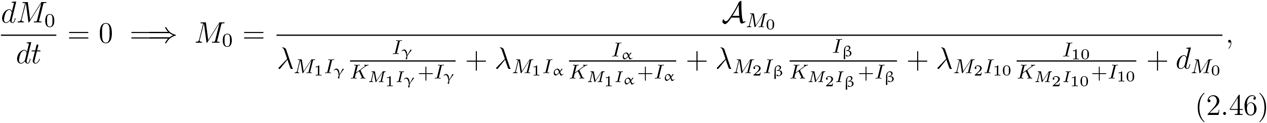

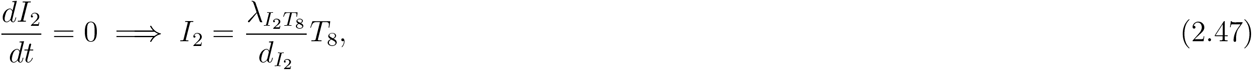

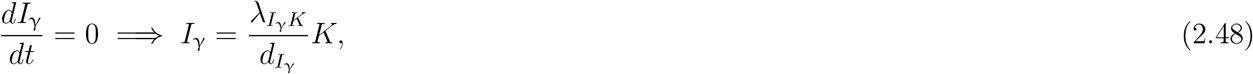

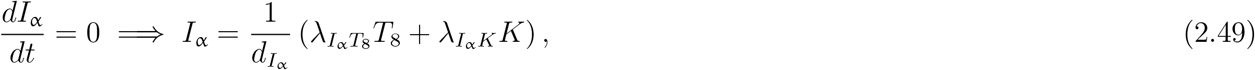

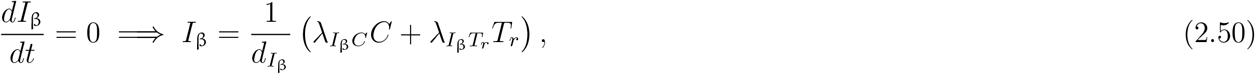

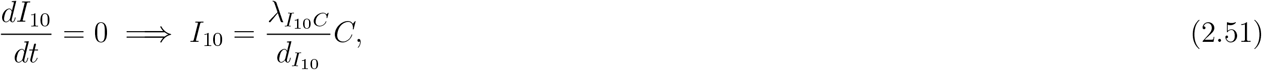

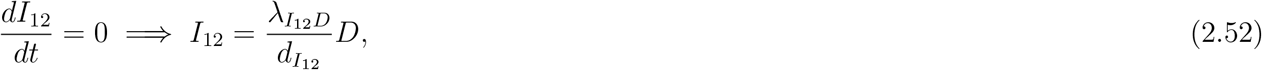

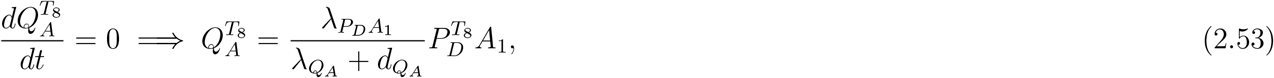

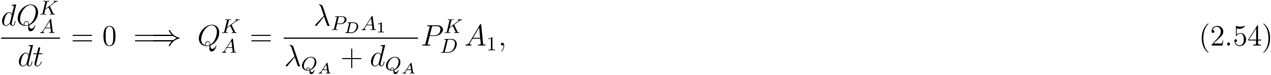

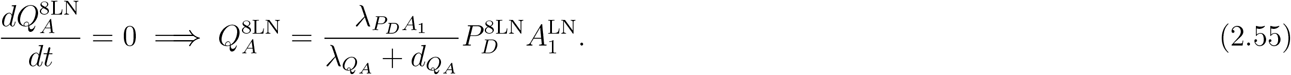

In particular, substituting (2.44), (2.45), and (2.53) – (2.55) into (2.10), (2.16), (2.36), (2.37), (2.60) (2.39), and (2.62), and simplifying, leads to

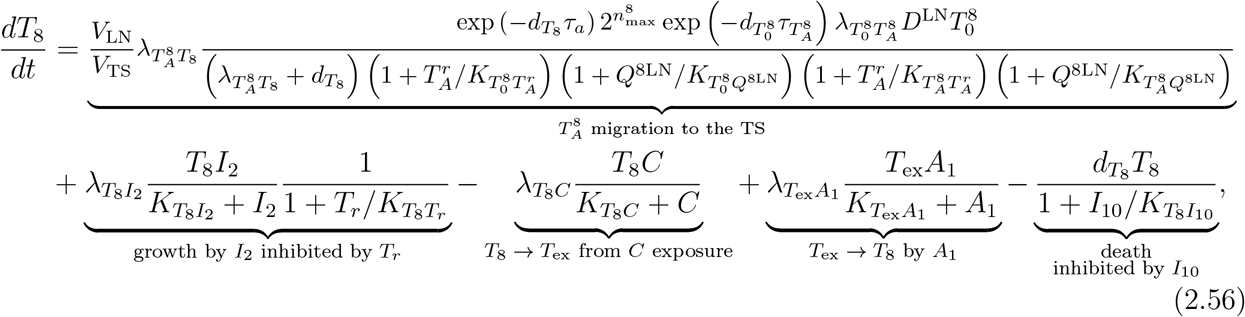

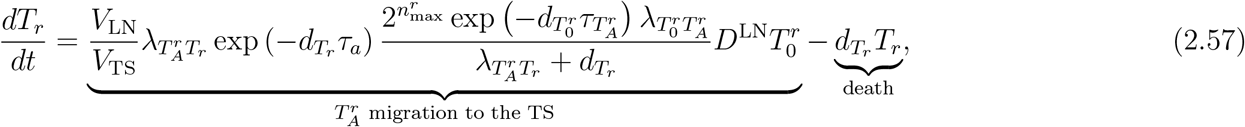

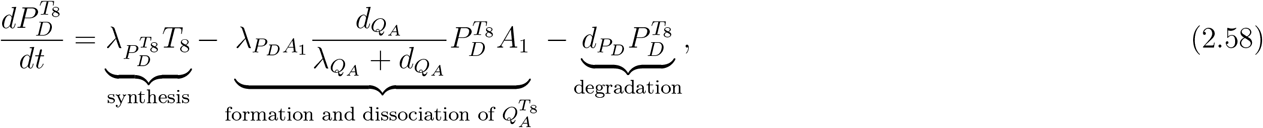

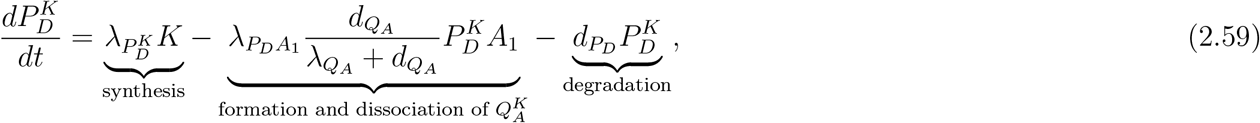

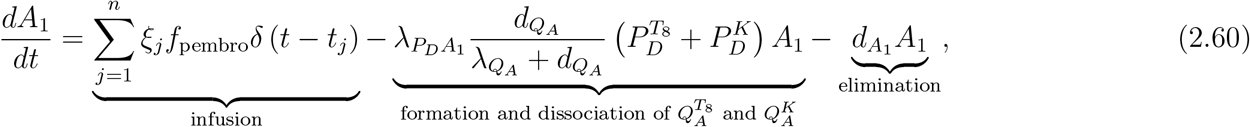

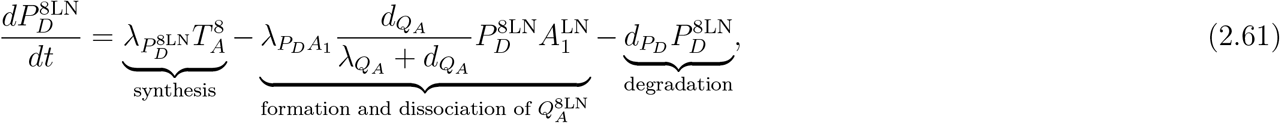

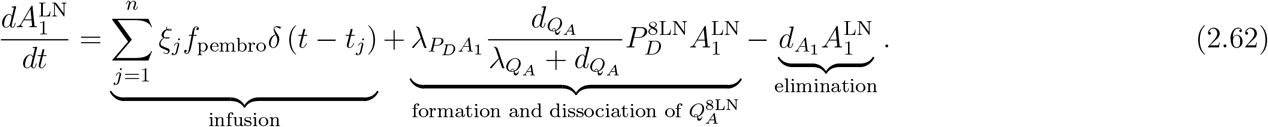

### 2.6 Conversion to an Agent-Based Model

To convert the ODE-based model to an ABM, we must realise that one-to-one agent representation of all biological entities is computationally intractable. As such, we first nondimensionalise the governing equations and model components, choosing parameter rescalings such that the original model’s structure is preserved. For each model species, *X*, we introduce a scaling factor, *g*_*X*_, and define the non-dimensional equivalent species, 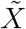, via 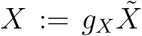. Then, substituting this into the model and simplifying leads to

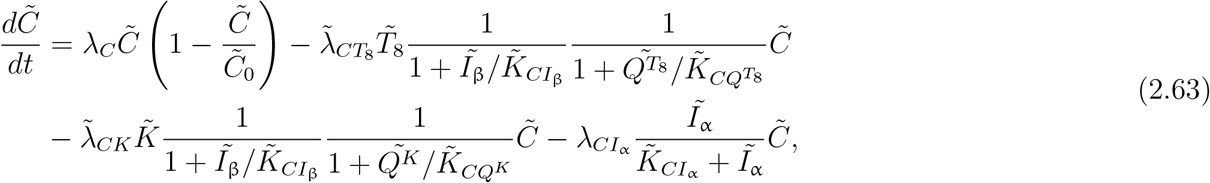

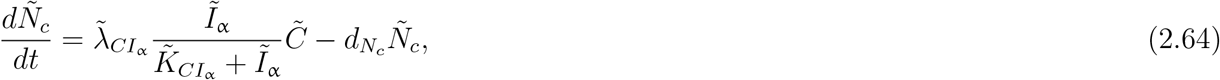

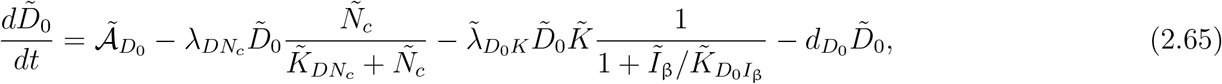

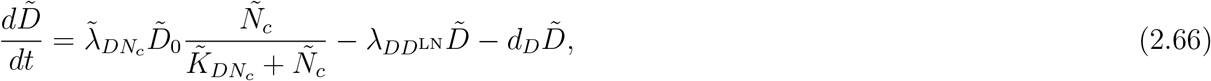

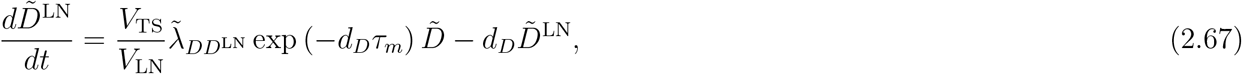

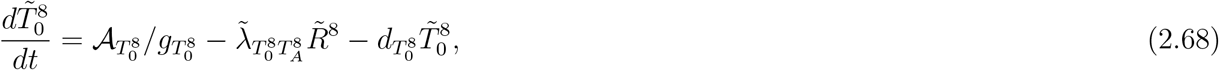

where 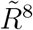 is defined as

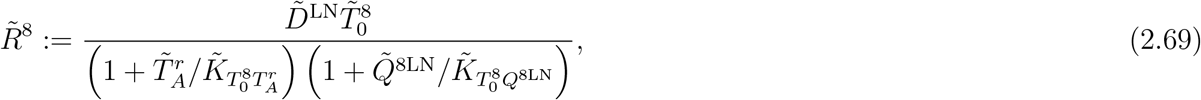

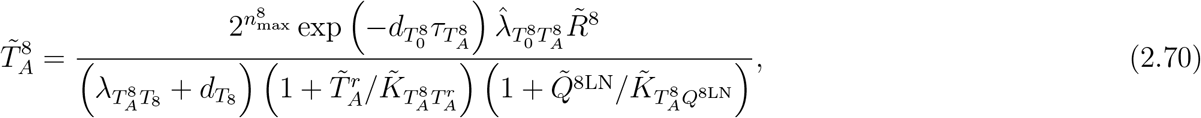

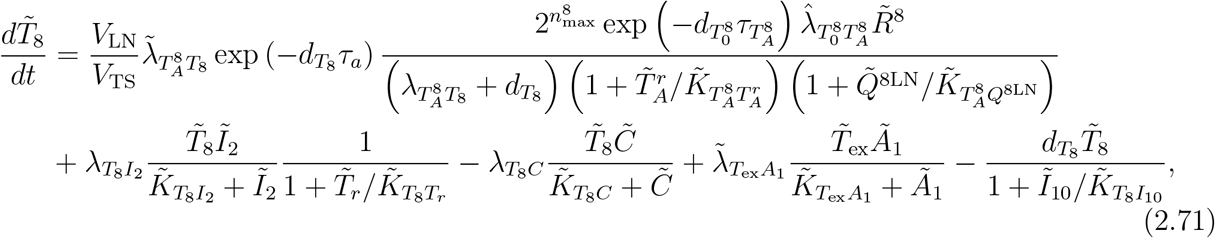

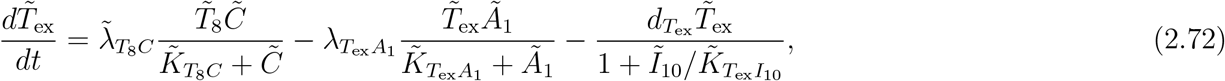

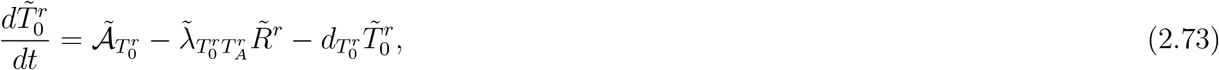

where 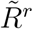 is defined as

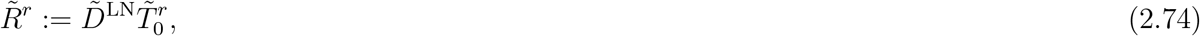

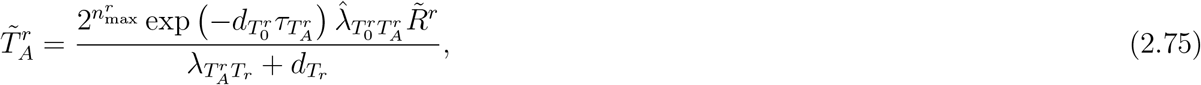

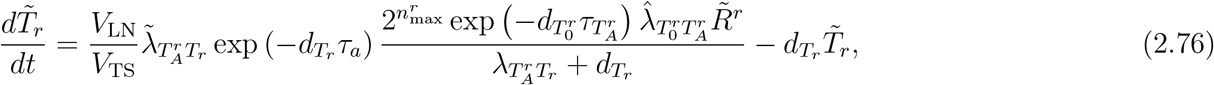

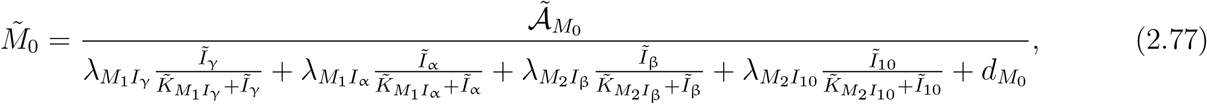

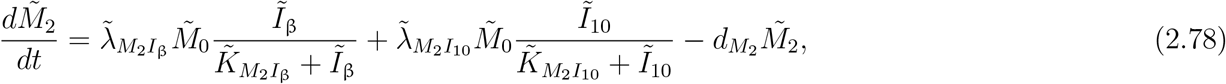

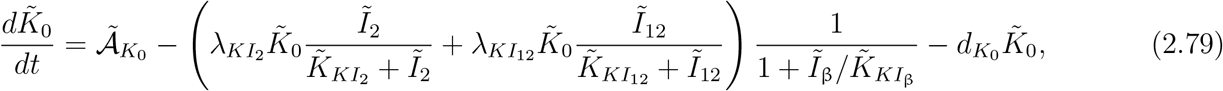

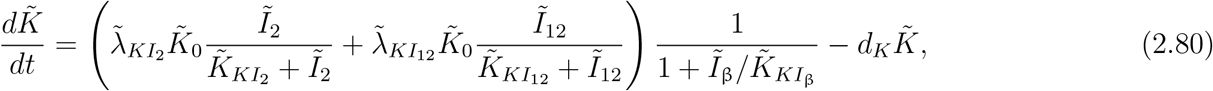

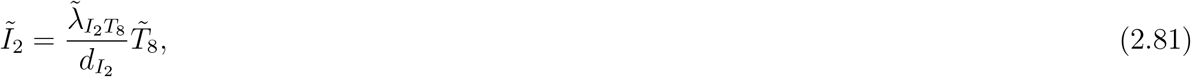

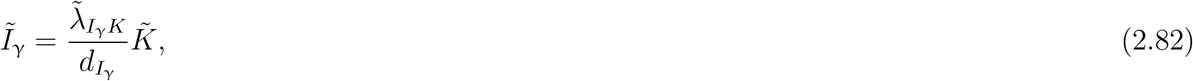

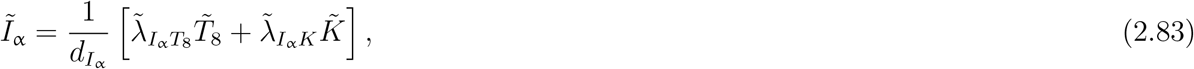

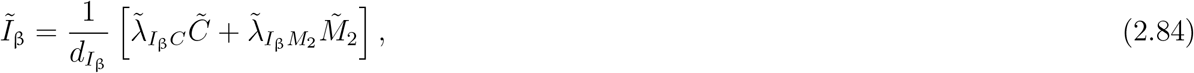

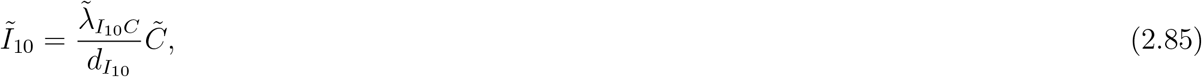

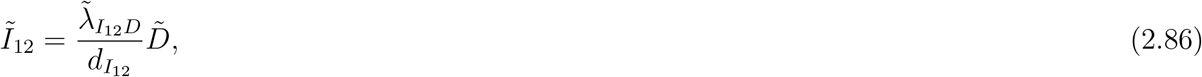

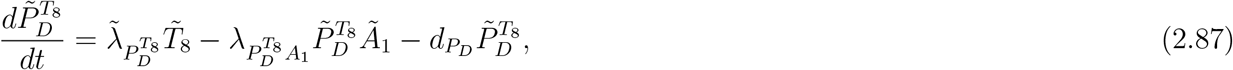

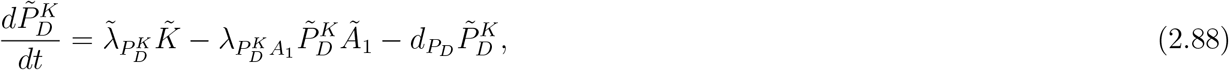

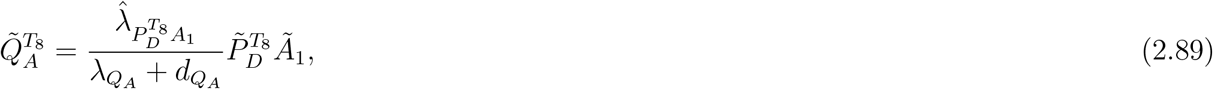

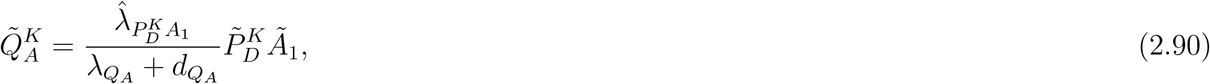

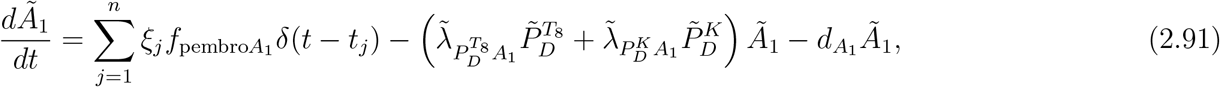

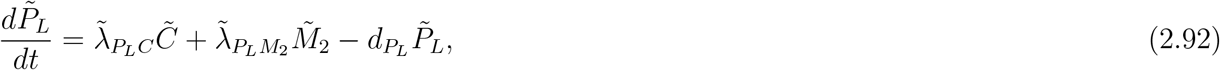

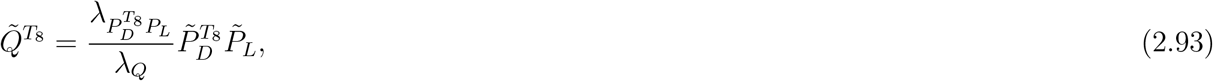

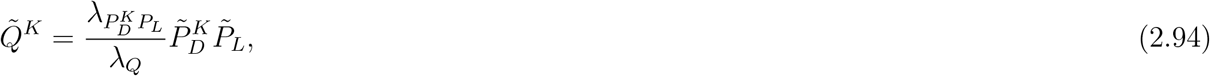

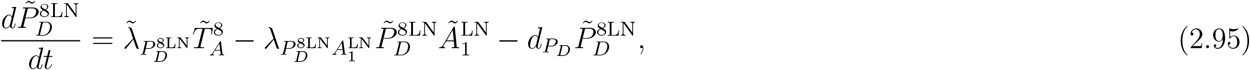

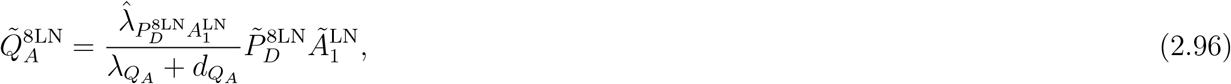

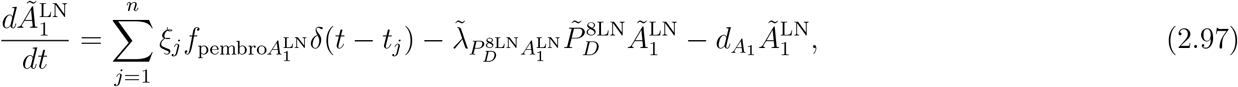

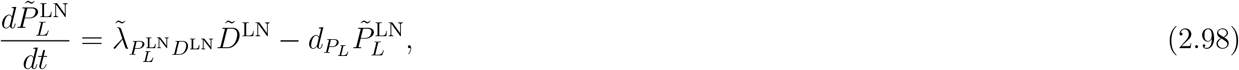

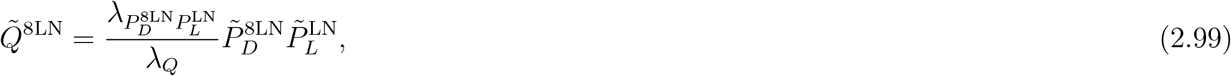

with scalings of

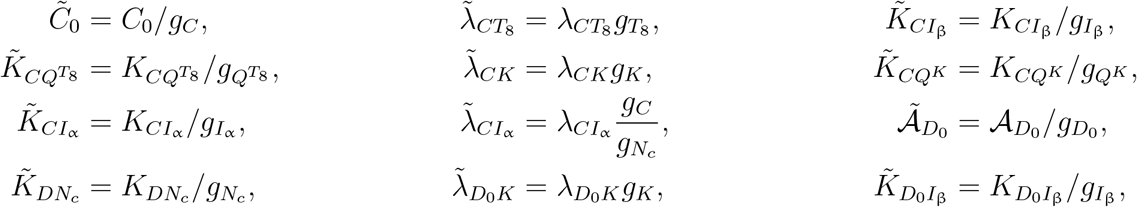

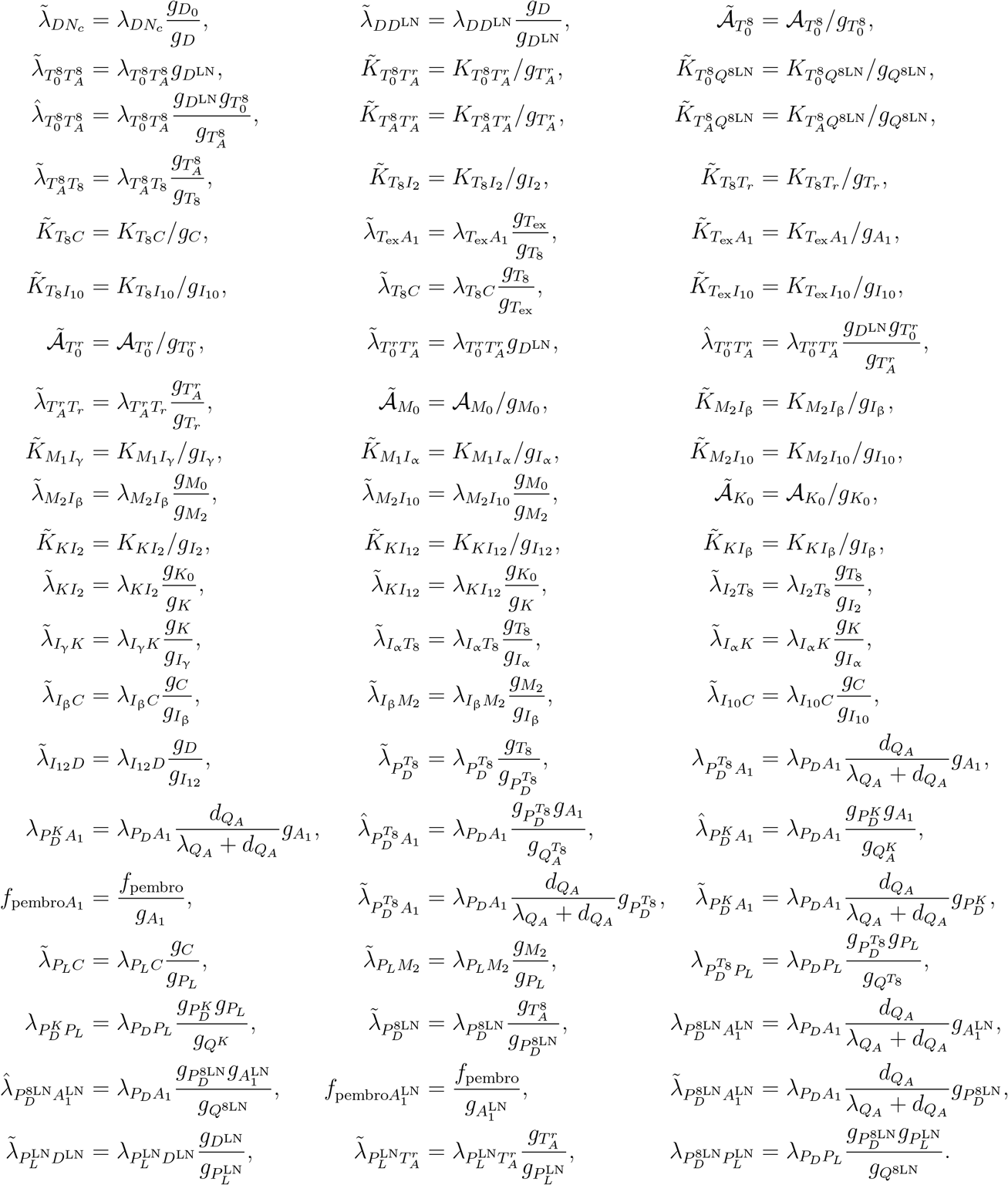

A derivation of this is shown in Appendix A, noting that we do not non-dimensionalise time, but refer to parameters as non-dimensionalised nonetheless.

For simplicity, we evolve the ABM on a uniform temporal grid, with spacing Δ*t*, employing a time-driven, synchronous update scheme for agent states and concentrations, which we now detail. Specifically, at each time step, a continuous-time event with rate *λ* is mapped to a per-step occurrence probability *p* = 1 − exp (−*λ*Δ*t*). We then perform a Bernoulli(*p*) trial, equivalently sampling *X* ∼ *U*[0, 1] and triggering the event if *X* ≤ *p*, to decide whether the event occurs for a given agent, after which the corresponding state-transition rule is applied. This construction is consistent with considering *T* ∼ Exp(*λ*), an exponential distribution with rate constant *λ*, since *p* = Pr[*T* ≤ Δ*t*] = 1 − exp (−*λ*Δ*t*), with *T* being interpreted as the waiting time for the corresponding continuous process to occur. In the mean-field limit, this yields a scheme that is the ABM analogue of the first-order forward Euler method for integrating the governing ODEs.

Processes in the ODE model can be organised into three categories: a) source/supply processes that introduce agents; b) loss processes that remove agents, such as degradation and lysis, and c) state transitions that change agent states (e.g., migration, activation, maturation, exhaustion, necrosis, complex formation/dissociation). For clarity, we provide examples of processes from each category, and explain how these are converted from the corresponding ODEs to the ABM.

We first consider the constant supply of immature DCs, with non-dimensionalised supply rate 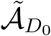.

This is equivalent to supplying 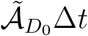 immature DCs at each time step. Taking into account the discrete nature of agents, this is equivalent to introducing 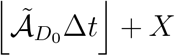 agents corresponding to 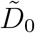, where 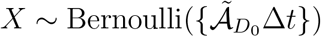. Here, ⌊°⌋ denotes the floor function, while {°} denotes the fractional part function, so that *x* = ⌊*x*⌋ + *x* for all *x* ∈ ℝ. We next translate the synthesis of PD-1 on the surface of NK cells, represented with non-dimensionalised rate constant 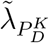. As such, for each activated NK cell agent, at each time step, we sample 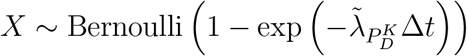. If *X* = 1, then we spawn an agent corresponding to 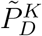.

Considering loss processes, we focus on the degradation of exhausted CD8+ T cells, which occurs with a non-dimensionalised rate 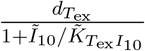. As such, at each time point, for each exhausted CD8+ T cell agent, we sample 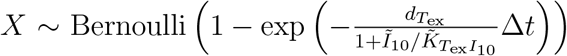, and if *X* = 1, then we remove the corresponding agent.

Finally, we consider processes involving transitions between agent states, which are amongst the most interesting and complex to implement. We consider the necrosis of viable cancer cells by TNF, which occurs with non-dimensionalised rate 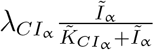. As such, at each time point, for each viable cancer cell agent, we sample 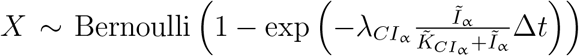. If *X* = 1, then we kill the corresponding agent, and spawn 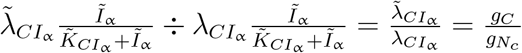 necrotic cancer cell agents. Recalling the discrete nature of agents, this amounts to spawning 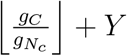 agents corresponding to 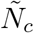, where 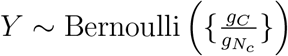.

The aforementioned conversion process provides a methodology for constructing ABMs from ODE systems with parameters calibrated to experimental data. Despite the non-spatial nature of ODEs, the agent-based framework easily accommodates the inclusion of spatial processes and interactions, such as diffusion and chemotaxis, by encoding them as agent-level properties and rules, should spatial structure be introduced.

## 3 Initial Conditions

### 3.1 Initial Conditions for Cells in the TS

The initial conditions for all cells in the TS are shown in Table 2 and are as in [35], except for naive macrophages. Justification for the choice of these values is in Appendix B.9.2.

**Table 2.**
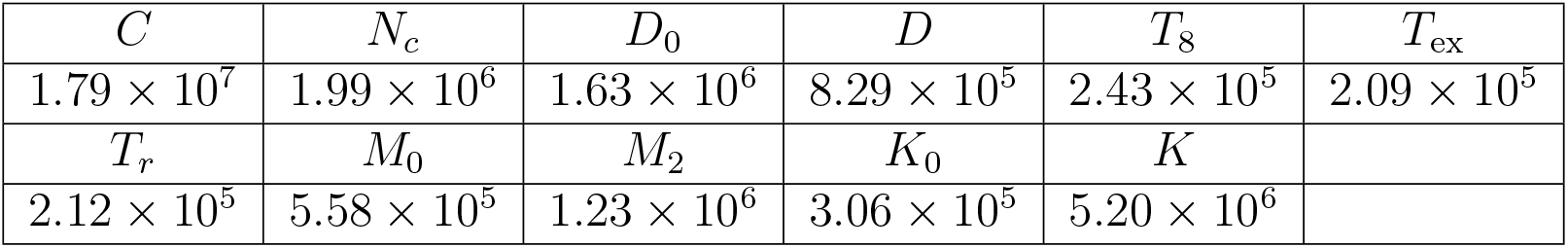
TS initial condition cell densities for the model. All values are in cell*/*cm^3^.

### 3.2 Initial Conditions for Cells in the TDLN

The initial conditions for all cells in the TDLN are shown in Table 3 and are as in [35], except for effector CD8+ T cells and effector Tregs. Justification for the choice of these values is in Appendix B.10.1 and Appendix B.10.2.

**Table 3.**
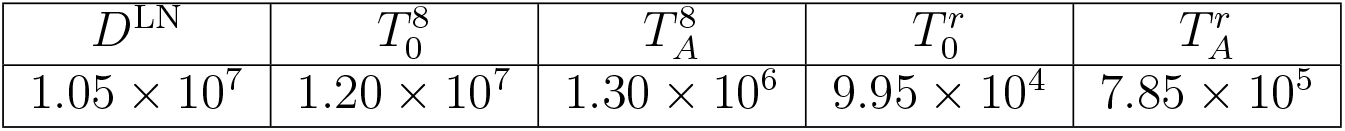
TDLN initial condition cell densities for the model. All values are in cell*/*cm^3^.

### 3.3 Initial Conditions for Cytokines in the TS

We choose the TS immune checkpoint-associated component initial conditions to be as in Table 4, with justification for the choice of these values in Appendix B.3.

**Table 4.**
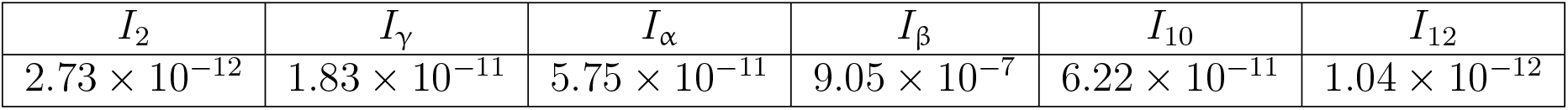
Cytokine initial conditions for the model. All values are in units of g*/*cm^3^.

| $I_2$ | $I_\gamma$ | $I_\alpha$ | $I_\beta$ | $I_{10}$ | $I_{12}$ |
| --- | --- | --- | --- | --- | --- |
| $2.73 \times 10^{-12}$ | $1.83 \times 10^{-11}$ | $5.75 \times 10^{-11}$ | $9.05 \times 10^{-7}$ | $6.22 \times 10^{-11}$ | $1.04 \times 10^{-12}$ |

### 3.4 Initial Conditions for Immune Checkpoint-Associated Components in the TS

We choose the TS immune checkpoint-associated component initial conditions to be as in Table 5, with justification for the choice of these values in Appendix B.12.

**Table 5.** TS immune checkpoint-associated component initial conditions for the model. All values are in units of molec*/*cm^3^.

| $P_D^{T_8}$ | $P_D^K$ | $P_L$ | $Q^{T_8}$ | $Q^K$ |
| --- | --- | --- | --- | --- |
| $6.70 \times 10^8$ | $2.87 \times 10^9$ | $3.55 \times 10^{12}$ | $5.06 \times 10^5$ | $2.17 \times 10^6$ |

### 3.5 Initial Conditions for Immune Checkpoint-Associated Components in the TDLN

We choose the TDLN immune checkpoint-associated component initial conditions to be as in Table 6, with justification for the choice of these values in Appendix B.13.

**Table 6.**
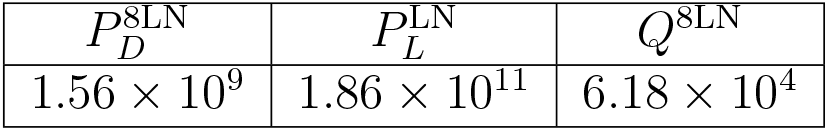
TDLN immune checkpoint-associated component initial conditions for the model. All values are in units of molec*/*cm^3^.

## 4 Results

In the absence of treatment, Figure 3 shows the time traces, up to 18 weeks, of the total cancer concentration, *V*, from the ODE model and ten ABM simulations, while Figure 4 shows the corresponding traces for all variables in the model.

**Figure 3.**
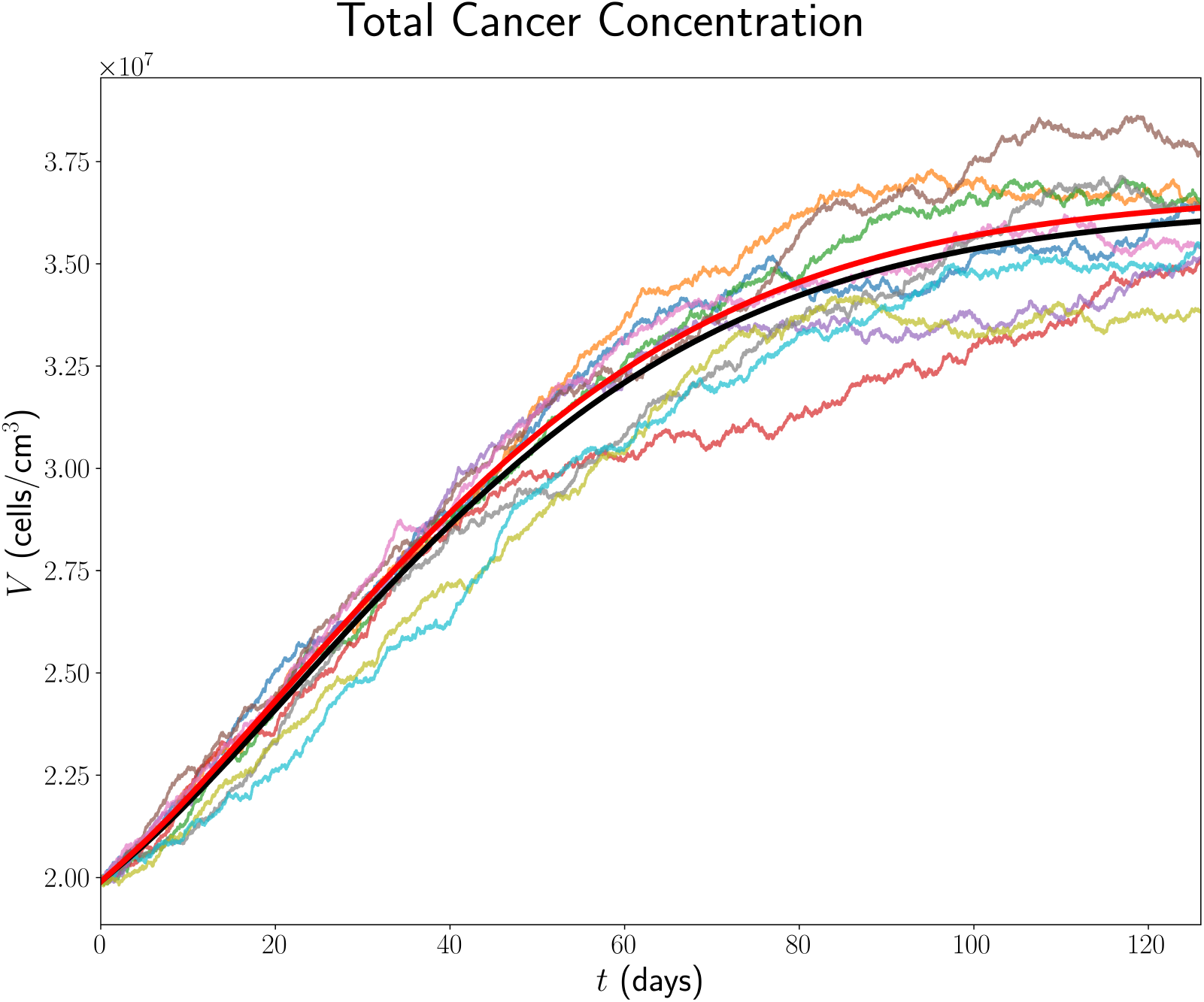
Time traces of *V* up to 18 weeks in the case of no treatment. The trajectory from the ODE model is shown in black, from the full model in [35] in red, from the full model in [35] in red, and the trajectories from ABM runs are coloured.

**Figure 4.**
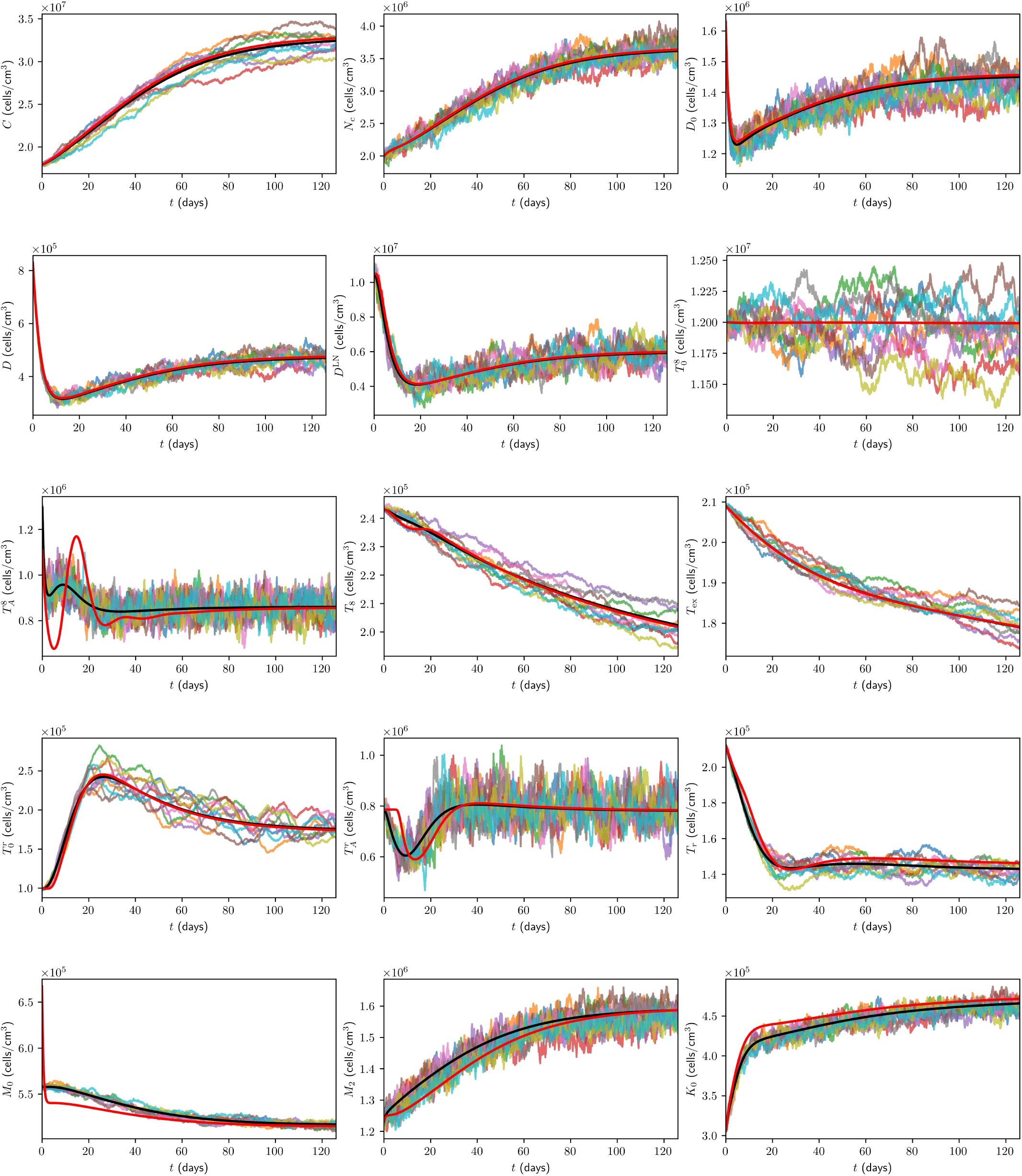
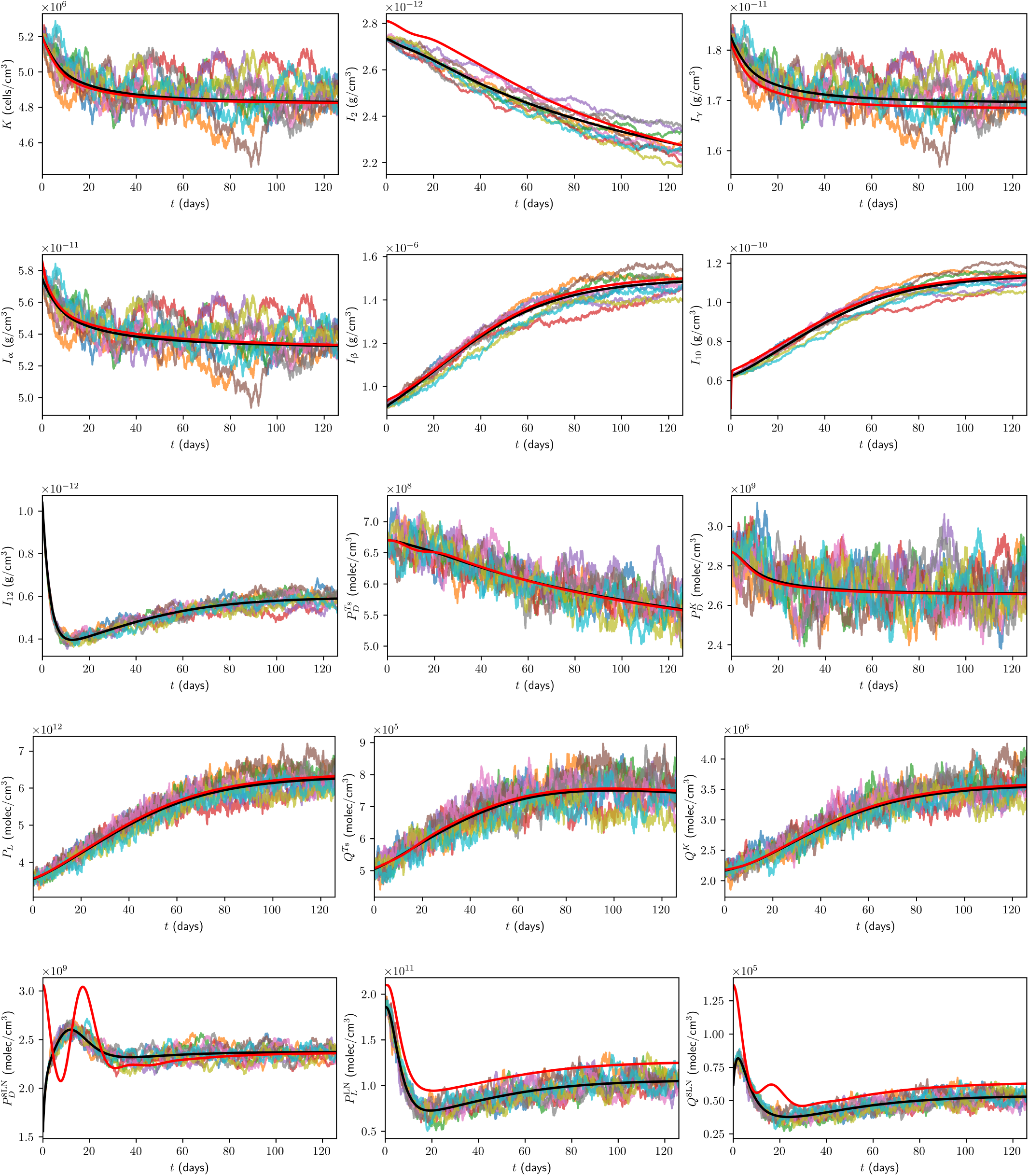
Time traces of variables in the model, in the case of no treatment, with the units of the variables as in Table 1. Trajectories from the ODE model are shown in black, from the full model in [35] in red, and the trajectories from ABM runs are coloured.

Furthermore, it is beneficial for us to compare the ODE and ABM models under neoadjuvant pembrolizumab therapy, which we assume commences at *t* = 0 days. As a benchmark for comparison, we consider treatment with 200 mg of pembrolizumab administered by intravenous infusion every 3 weeks, approved by the FDA for the first-line treatment of metastatic MSI-H/dMMR CRC [122]. Assuming a patient mass of *m* = 80 kg, these correspond to *ξ*_*j*_ = 200 mg, *t*_*j*_ = 21(*j* − 1), *n* = 6, *ξ*_pembro_ = 2.5 mg*/*kg, and *η*_pembro_ = 3 weeks. Time traces of *V* and all model variables, up to 18 weeks, from the ODE model and ten ABM simulations are shown in Figure 5 and Figure 6 in the case of this treatment, respectively.

**Figure 5.**
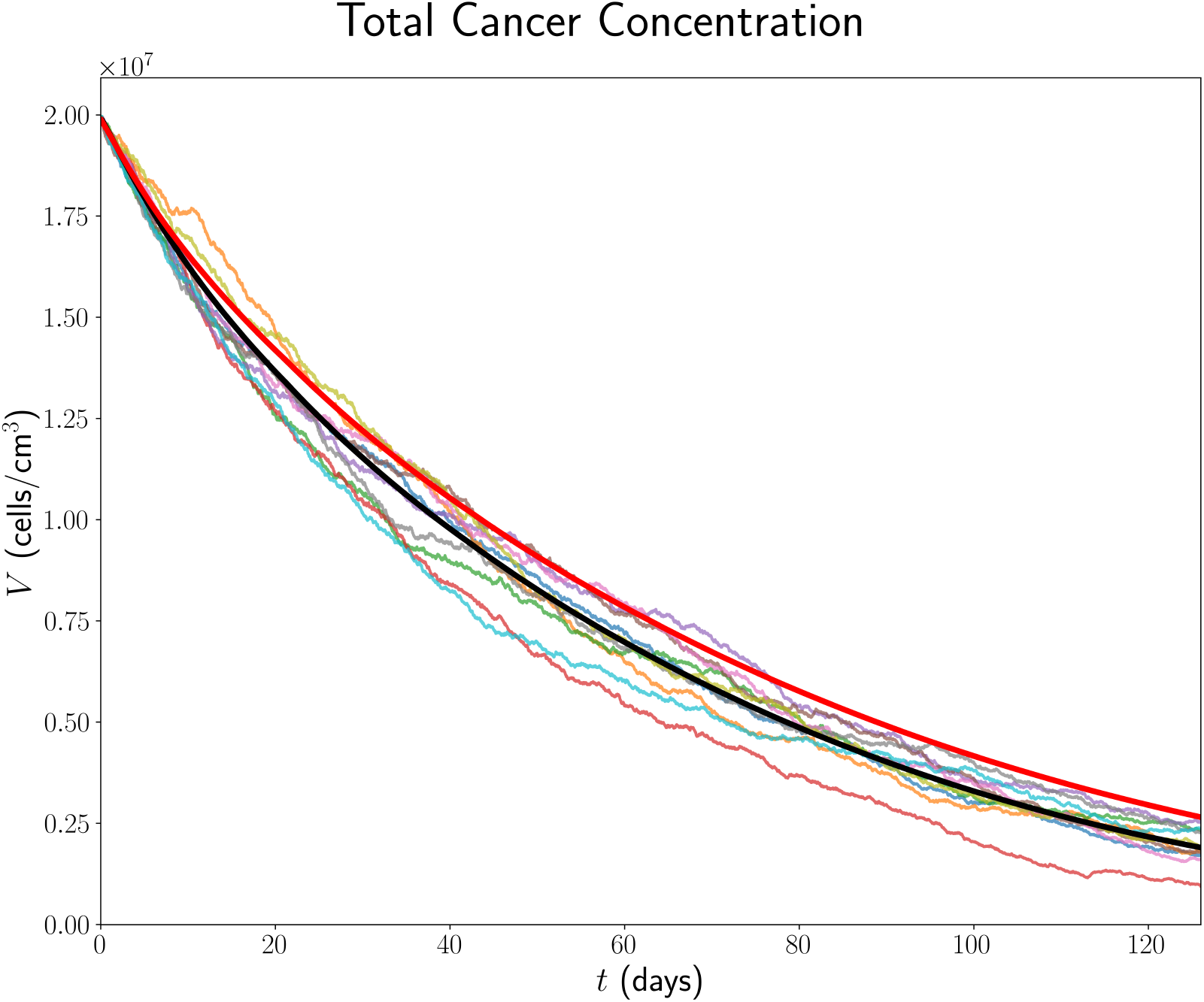
Time traces of *V* up to 18 weeks in the case of treatment with triweekly 200 mg pembrolizumab. The trajectory from the ODE model is shown in black, from the full model in [35] in red, and the trajectories from ABM runs are coloured.

**Figure 6.**
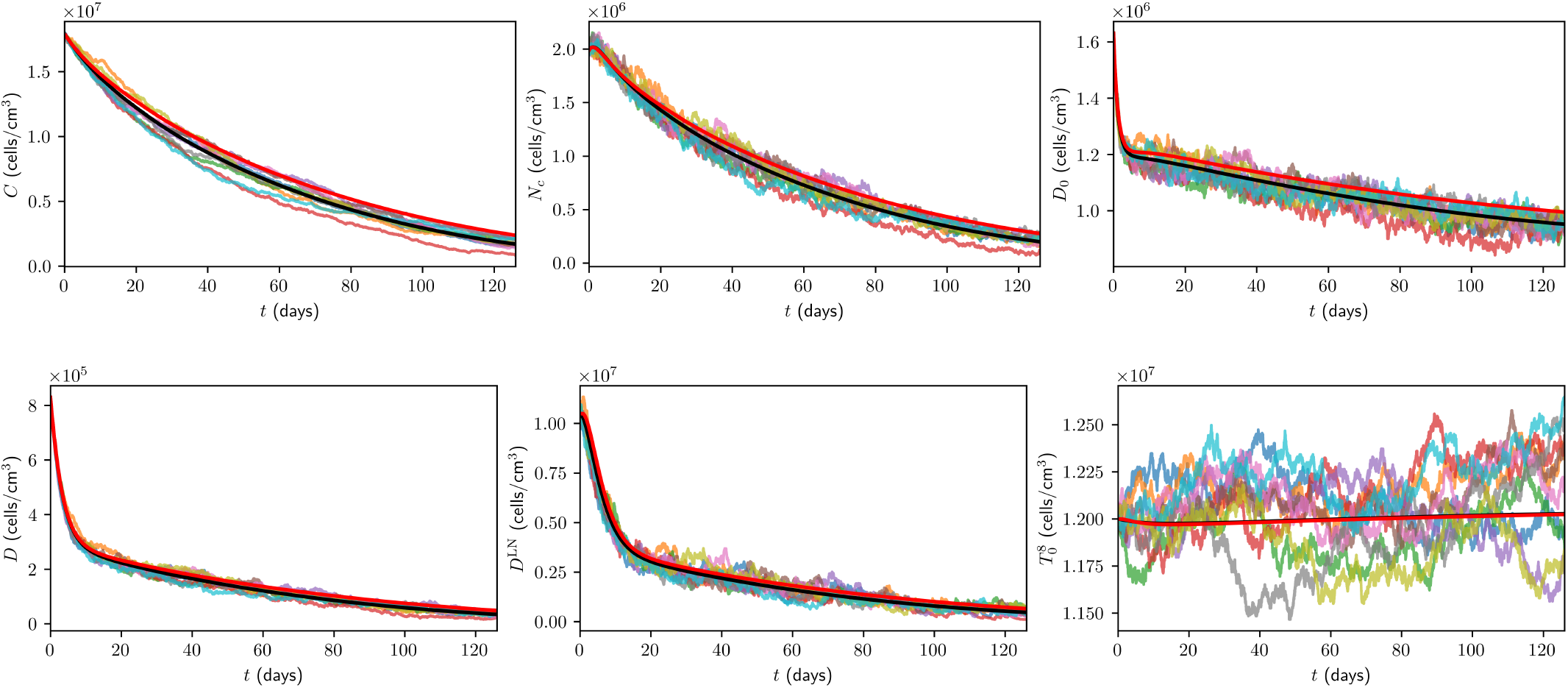
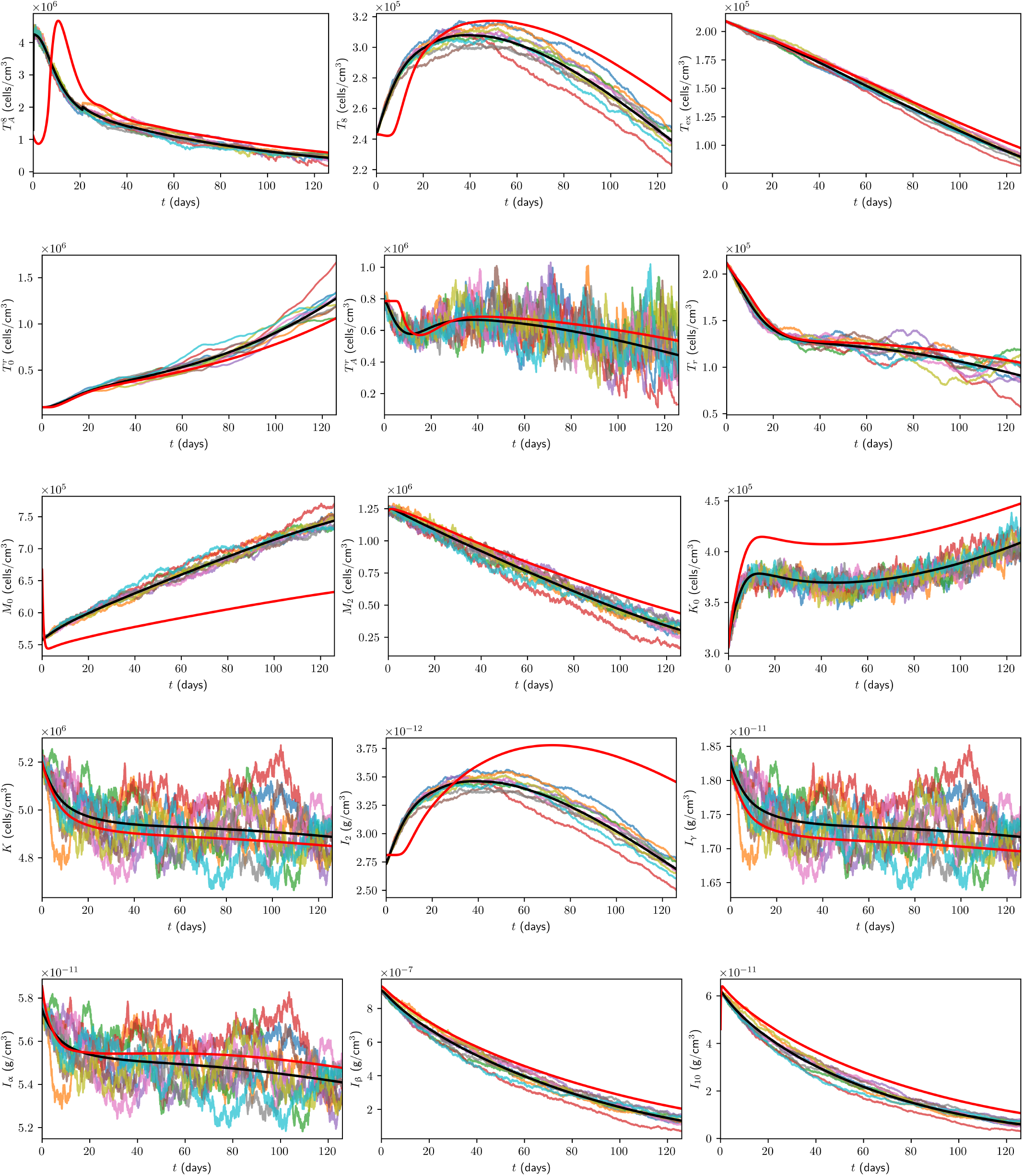
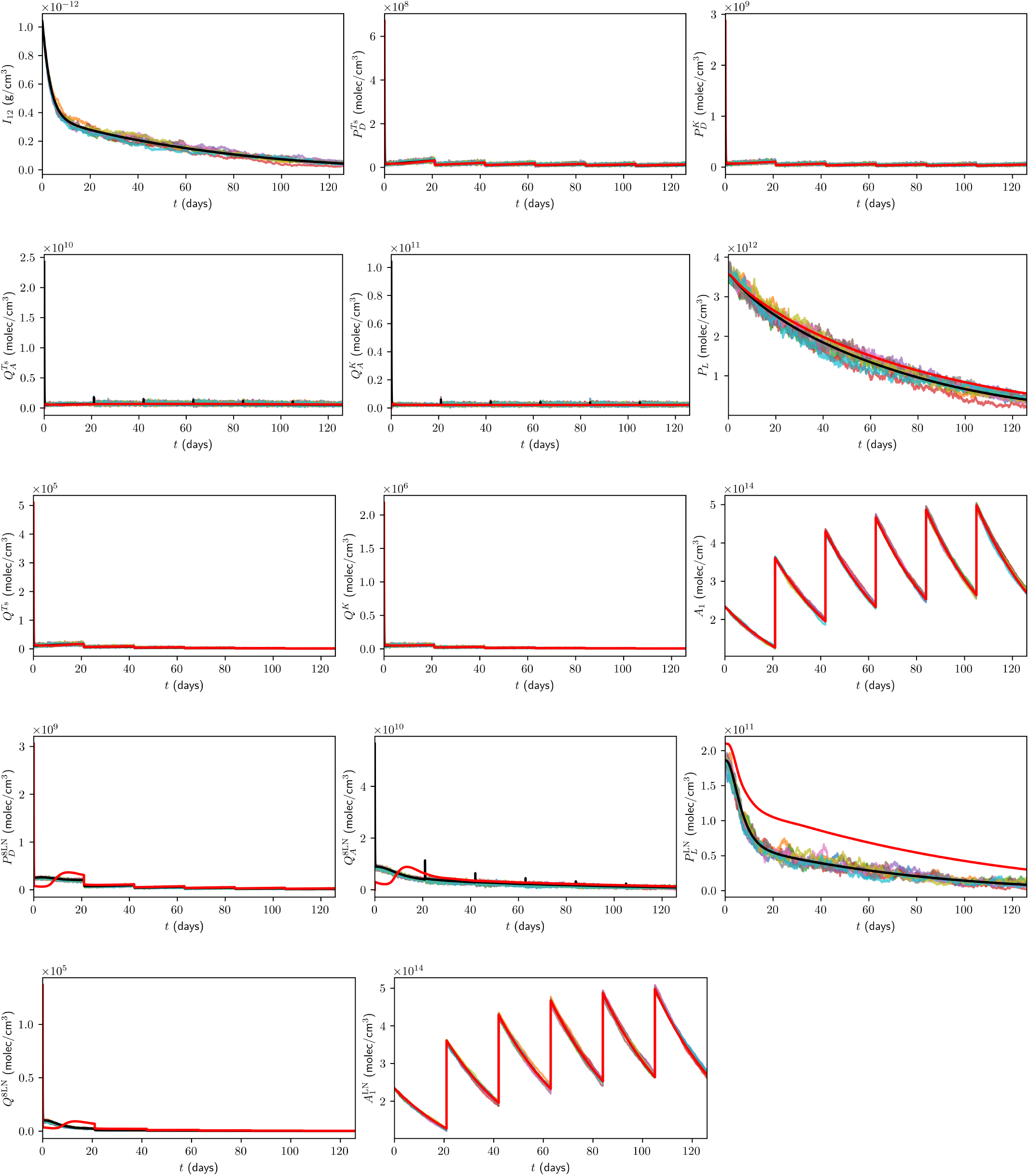
Time traces of variables in the model, in the case of treatment with triweekly 200 mg pembrolizumab, with the units of the variables as in Table 1. Trajectories from the ODE model are shown in black, from the full model in [35] in red, and the trajectories from ABM runs are coloured.

All ODE simulations of the minimal model were performed in MATLAB using the ode23 solver with the initial conditions stated in Section 3, whereas all DDE simulations of the full model were performed in MATLAB using the dde23 solver with the initial conditions stated in [35].

All ABM simulations were performed in Python 3.11 using the Mesa framework [123] with a constant time step of Δ*t* = 0.01 day, and with group sizes as follows: 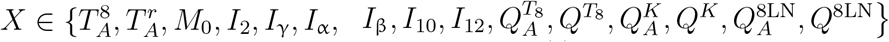, we set *g*_*X*_ = 1; for all *Y* ∈ {*D*_0_, *D, T*_8_, *T*_ex_, *T*_*r*_, *M*_2_, *K*_0_, *K*}, we set 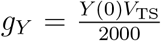 and for all 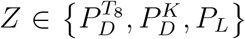, we set 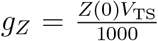. Additionally 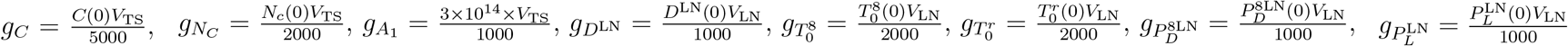, and 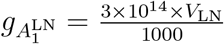.

## Discussion

Referencing Figure 3, Figure 4, Figure 5, and Figure 6, it is evident that the minimal model accurately reproduces the trajectories of the full model under both treatment and no-treatment conditions. In particular, the core dynamics are preserved, and we refer the reader to [35] for a detailed discussion on their biological implications and corresponding mechanistic insights. However, it is important to note that we observe some minor deviations, particularly after a couple of weeks, for variables related to the concentration of effector CD8+ T cells, naive macrophages, and immune checkpoint-associated components in the TDLN. These deviations are expected due to the nature and application of QSSA, which results in different initial conditions for these variables compared to the full and minimal models. Additionally, there are minor differences in the steady states and initial conditions of PD-L1 and PD-1/PD-L1 complex concentrations in the TDLN, which contribute to these deviations. This is more pronounced under treatment: PD-L1 in the TDLN originates solely from mature DCs, and pembrolizumab-driven tumour reduction decreases DAMP secretion from necrotic cancer cells, thereby reducing DC maturation and the influx of mature DCs to the TDLN. However, PD-L1 in the TDLN only influences the system through PD-1/PD-L1 complex formation, which appears through inhibition constants, rendering its absolute concentration, like that of several cytokines, less critical; accordingly, these deviations have minimal impact on the trajectories of other state variables, as observed.

In constructing a minimal model of pembrolizumab therapy in laMCRC, we removed variables corresponding to the concentration of DAMPs, CD4+ T cells and associated variables, and M1 macrophages, alongside their associated parameters. Additionally, with the application of QSSAs, further eliminated explicit dependencies on effector CD8+ T cells and Tregs in the TDLN, naive macrophages, IL-2, IFN-γ, TNF, TGF-β, IL-10, IL-12, and the PD-1/pembrolizumab complex on effector NK cells and effector CD8+ T cells in the TS and TDLN. This, in tandem with only retaining the most influential processes, results in an approximate 50% reduction in system dimensionality and a nearly 50% reduction in the number of parameters compared to the full model—representing a significant simplification while still accurately replicating model trajectories.

Consequently, replacing the integrals in the full model with point estimates is justified, suggesting that delay integro-differential equations may be unnecessarily complex for modelling the immunobiology of MSI-H/dMMR CRC, and that ODEs may be sufficient. We also hypothesise that the minimal model cannot be easily simplified or reduced without significantly affecting model trajectories, so that each component of the minimal model is functionally important, contributing meaningfully to the system’s behaviour. However, verification of this requires methods such as sensitivity analysis [124–127] and identifiability analysis [128], which are beyond the scope of this work.

We observe that the ABM faithfully replicates the trajectories of the minimal ODE model under both treatment and no-treatment conditions, considering Figure 3, Figure 4, Figure 5, and Figure 6, with the ABM exhibiting stochastic fluctuations around the ODE trajectories—as expected, since the ODE system represents the mean-field limit of the ABM. In particular, there is pronounced variation and stochasticity arising in model populations that undergo significant proliferation, especially cancer cells, NK cells, and T cells, since randomness can propagate, as proliferation events and bursty clonal expansion amplify and propagate randomness over time. This, in turn, induces fluctuations in the concentrations of secreted cytokines and in the expression of immune checkpoints, yielding correlated oscillations in IL-2, IFN-γ, TNF, TGF-β, IL-10, and tumoural PD-L1, which often directly influence the concentration of these cells, creating positive feedback loops, and reinforcing stochasticity. For example, IL-2 mediates effector CD8+ T cell expansion in the TS, whereas TGF-β inhibits effector CD8+ T cell and activated NK cell cytotoxicity against cancer cells. This results in amplified variability in effector and cancer cell concentrations in the ABM, manifesting as oscillations around the deterministic mean-field trajectories of the ODE system—nonetheless, maintaining close agreement with those governed by the deterministic model.

It should be noted that the models have several limitations, many of which exist for simplicity, but addressing these issues offers exciting avenues for future research.

- We ignored spatial effects in the model; however, their resolution can provide information about the distribution and clustering of different immune cell types in the TME and their clinical implications [129, 130]. Nevertheless, the ABM is naturally amenable to adding spatial mechanisms— including chemotaxis along cytokine gradients, explicit tumour and TDLN geometries, and the diffusion and advection of cells, cytokines, and proteins.
- We did not consider T cell avidity, the overall strength of a TCR-pMHC interaction, which governs whether a cancer cell will be successfully killed [131]. In particular, high-avidity T cells are necessary for lysing cancer cells and durable tumour eradication, while low-avidity T cells are ineffective and may inhibit high-avidity T cells [132, 133].
- We also did not consider the influence of cytokines in the TDLN for T cell activation and proliferation, which are important in influencing effector T cell differentiation [134, 135].
- The model does not explicitly account for additional anatomical compartments such as the spleen, nor does it directly model lymph node metastasis, which may impact the accuracy of systemic immune dynamics and tumour-specific responses; however, these features can be readily accommodated in ABMs.

In this work, we have constructed a minimal ODE model of neoadjuvant pembrolizumab therapy in laMCRC, and verified that it accurately reproduces the state-variable trajectories of larger, more comprehensive models, thereby preserving the most significant biological interactions and resultant tumour–immune dynamics. This, in turn, enhances interpretability and parameter identifiability and, combined with the model’s self-contained nature and high extensibility, provides a robust foundation for future analysis and experimentation. Building on this foundation, we presented a practical and efficient workflow for translating differential-equation-based models into ABMs, enabling efficient and accurate calibration of model parameters and stochastic, individual-level representation while preserving consistency with the underlying deterministic system. Applying this framework to the minimal ODE model, we showed that the calibrated ABM faithfully reproduces its trajectories, thereby providing a principled basis for incorporating spatial effects, additional anatomical compartments, and patient-specific heterogeneity in future studies. Collectively, these advances establish a rigorous, flexible modelling paradigm that bridges parsimonious, calibrated deterministic dynamical systems with calibrated, stochastic agent-based modelling—applicable to laMCRC and other biological systems— thereby accelerating treatment optimisation and personalisation and improving therapeutic outcomes.

## 6 CRediT Authorship Contribution Statement

**Georgio Hawi**: conceptualisation, data curation, formal analysis, funding acquisition, investigation, methodology, project administration, resources, software, validation, visualisation, writing — original draft, writing — review & editing.

**Peter S. Kim**: conceptualisation, formal analysis, funding acquisition, investigation, methodology, project administration, resources, supervision, validation, visualisation, writing — original draft, writing — review & editing.

**Peter P. Lee**: conceptualisation, formal analysis, investigation, methodology, project administration, resources, supervision, validation, visualisation, writing — original draft, writing — review & editing.

## 8 Data Availability

All data and procedures are available within the manuscript and its Supporting Information file. Accompanying code is available at https://doi.org/10.5281/zenodo.16930420.

## 9 Acknowledgements

This work was supported by an Australian Government Research Training Program Scholarship. PSK gratefully acknowledges support from the Australian Research Council Discovery Project (DP230100485).

