## Supporting Information for "Modelling Immune Dynamics in Locally Advanced MSI-H/dMMR Colorectal Cancer with Neoadjuvant Pembrolizumab Treatment: From Differential Equations to an Agent-Based Framework"

#### A Non-dimensionalised ODE Model Derivation

Substituting $X = g_X\tilde{X}$, for each species $X$, into the model leads to

$$\begin{aligned}
\frac{d(g_C\tilde{C})}{dt} &= \lambda_C g_C\tilde{C}\left(1-\frac{g_C\tilde{C}}{C_0}\right) - \lambda_{CT_8} g_{T_8}\tilde{T}_8 \frac{1}{1+g_{I_\beta}\tilde{I}_\beta/K_{CI_\beta}} \frac{1}{1+g_{Q^{T_8}}\tilde{Q}^{T_8}/K_{CQ^{T_8}}} g_C\tilde{C} \\
&\quad - \lambda_{CK} g_K\tilde{K}\frac{1}{1+g_{I_\beta}\tilde{I}_\beta/K_{CI_\beta}}\frac{1}{1+g_{Q^K}\tilde{Q}^K/K_{CQ^K}} g_C\tilde{C} - \lambda_{CI_\alpha}\frac{g_{I_\alpha}\tilde{I}_\alpha}{K_{CI_\alpha}+g_{I_\alpha}\tilde{I}_\alpha} g_C\tilde{C}, \\
\frac{d(g_{N_c}\tilde{N}_c)}{dt} &= \lambda_{CI_\alpha}\frac{g_{I_\alpha}\tilde{I}_\alpha}{K_{CI_\alpha}+g_{I_\alpha}\tilde{I}_\alpha} g_C\tilde{C} - d_{N_c} g_{N_c}\tilde{N}_c, \\
\frac{d(g_{D_0}\tilde{D}_0)}{dt} &= \mathcal{A}_{D_0} - \lambda_{DN_c} g_{D_0}\tilde{D}_0 \frac{g_{N_c}\tilde{N}_c}{K_{DN_c}+g_{N_c}\tilde{N}_c} - \lambda_{D_0K} g_{D_0}\tilde{D}_0 g_K\tilde{K}\frac{1}{1+g_{I_\beta}\tilde{I}_\beta/K_{D_0I_\beta}} - d_{D_0} g_{D_0}\tilde{D}_0, \\
\frac{d(g_D\tilde{D})}{dt} &= \lambda_{DN_c} g_{D_0}\tilde{D}_0\frac{g_{N_c}\tilde{N}_c}{K_{DN_c}+g_{N_c}\tilde{N}_c} - \lambda_{DD^{\mathrm{LN}}} g_D\tilde{D} - d_D g_D\tilde{D}, \\
\frac{d(g_{D^{\mathrm{LN}}}\tilde{D}^{\mathrm{LN}})}{dt} &= \frac{V_{\mathrm{TS}}}{V_{\mathrm{LN}}}\lambda_{DD^{\mathrm{LN}}} e^{-d_D\tau_m} g_D\tilde{D} - d_D g_{D^{\mathrm{LN}}}\tilde{D}^{\mathrm{LN}}, \\
\frac{d(g_{T_0^8}\tilde{T}_0^8)}{dt} &= \mathcal{A}_{T_0^8} - \lambda_{T_0^8T_A^8}\tilde{R}^8 - d_{T_0^8} g_{T_0^8}\tilde{T}_0^8,
\end{aligned}$$

where $\tilde{R}^8$ is defined as

$$
\begin{aligned}
\tilde{R}^8 &:= \frac{g_{D^{\mathrm{LN}}}\tilde{D}^{\mathrm{LN}} g_{T_0^8}\tilde{T}_0^8}{\left(1+g_{T_A^r}\tilde{T}_A^r/K_{T_0^8T_A^r}\right)\left(1+g_{Q^{8\mathrm{LN}}}\tilde{Q}^{8\mathrm{LN}}/K_{T_0^8Q^{8\mathrm{LN}}}\right)},\\
g_{T_A^8}\tilde{T}_A^8 &= \frac{2^{n_{\max}^8}e^{-d_{T_0^8}\tau_{T_A^8}}\lambda_{T_0^8T_A^8}\tilde{R}^8}{\left(\lambda_{T_A^8T_8}+d_{T_8}\right)\left(1+g_{T_A^r}\tilde{T}_A^r/K_{T_A^8T_A^r}\right)\left(1+g_{Q^{8\mathrm{LN}}}\tilde{Q}^{8\mathrm{LN}}/K_{T_A^8Q^{8\mathrm{LN}}}\right)},\\
\frac{d(g_{T_8}\tilde{T}_8)}{dt} &= \frac{V_{\mathrm{LN}}}{V_{\mathrm{TS}}}\lambda_{T_A^8T_8}e^{-d_{T_8}\tau_a}g_{T_A^8}\tilde{T}_A^8+\lambda_{T_8I_2}\frac{g_{T_8}\tilde{T}_8g_{I_2}\tilde{I}_2}{K_{T_8I_2}+g_{I_2}\tilde{I}_2}\frac{1}{1+g_{T_r}\tilde{T}_r/K_{T_8T_r}}\\
&\quad-\lambda_{T_8C}\frac{g_{T_8}\tilde{T}_8g_C\tilde{C}}{K_{T_8C}+g_C\tilde{C}}+\lambda_{T_{\mathrm{ex}}A_1}\frac{g_{T_{\mathrm{ex}}}\tilde{T}_{\mathrm{ex}}g_{A_1}\tilde{A}_1}{K_{T_{\mathrm{ex}}A_1}+g_{A_1}\tilde{A}_1}-\frac{d_{T_8}g_{T_8}\tilde{T}_8}{1+g_{I_{10}}\tilde{I}_{10}/K_{T_8I_{10}}},\\
\frac{d(g_{T_{\mathrm{ex}}}\tilde{T}_{\mathrm{ex}})}{dt} &= \lambda_{T_8C}\frac{g_{T_8}\tilde{T}_8g_C\tilde{C}}{K_{T_8C}+g_C\tilde{C}}-\lambda_{T_{\mathrm{ex}}A_1}\frac{g_{T_{\mathrm{ex}}}\tilde{T}_{\mathrm{ex}}g_{A_1}\tilde{A}_1}{K_{T_{\mathrm{ex}}A_1}+g_{A_1}\tilde{A}_1}-\frac{d_{T_{\mathrm{ex}}}g_{T_{\mathrm{ex}}}\tilde{T}_{\mathrm{ex}}}{1+g_{I_{10}}\tilde{I}_{10}/K_{T_{\mathrm{ex}}I_{10}}},\\
\frac{d(g_{T_0^r}\tilde{T}_0^r)}{dt} &= \mathcal{A}_{T_0^r}-\lambda_{T_0^rT_A^r}\tilde{R}^r-d_{T_0^r}g_{T_0^r}\tilde{T}_0^r,
\end{aligned}
$$

where $\tilde{R}^r$ is defined as

$$
\begin{aligned}
\tilde{R}^r &:= g_{D^{\mathrm{LN}}}\tilde{D}^{\mathrm{LN}}g_{T_0^r}\tilde{T}_0^r,\\
g_{T_A^r}\tilde{T}_A^r &= \frac{2^{n_{\max}^r}e^{-d_{T_0^r}\tau_{T_A^r}}\lambda_{T_0^rT_A^r}\tilde{R}^r}{\lambda_{T_A^rT_r}+d_{T_r}},\\
\frac{d(g_{T_r}\tilde{T}_r)}{dt} &= \frac{V_{\mathrm{LN}}}{V_{\mathrm{TS}}}\lambda_{T_A^rT_r}g_{T_A^r}e^{-d_{T_r}\tau_a}\tilde{T}_A^r-d_{T_r}g_{T_r}\tilde{T}_r,\\
g_{M_0}\tilde{M}_0 &= \frac{\mathcal{A}_{M_0}}{\lambda_{M_1I_\gamma}\frac{g_{I_\gamma}I_\gamma}{K_{M_1I_\gamma}+g_{I_\gamma}I_\gamma}+\lambda_{M_1I_\alpha}\frac{g_{I_\alpha}I_\alpha}{K_{M_1I_\alpha}+g_{I_\alpha}I_\alpha}+\lambda_{M_2I_\beta}\frac{I_\beta}{g_{I_\beta}K_{M_2I_\beta}+I_\beta}+\lambda_{M_2I_{10}}\frac{I_{10}}{K_{M_2I_{10}}+g_{I_{10}}I_{10}}+d_{M_0}},\\
\frac{d(g_{M_2}\tilde{M}_2)}{dt} &= \lambda_{M_2I_\beta}g_{M_0}\tilde{M}_0\frac{I_\beta}{g_{I_\beta}K_{M_2I_\beta}+I_\beta}+\lambda_{M_2I_{10}}g_{M_0}\tilde{M}_0\frac{I_{10}}{K_{M_2I_{10}}+g_{I_{10}}I_{10}}-d_{M_2}g_{M_2}\tilde{M}_2,\\
\frac{d(g_{K_0}\tilde{K}_0)}{dt} &= \mathcal{A}_{K_0}-\left(\lambda_{KI_2}g_{K_0}\tilde{K}_0\frac{g_{I_2}\tilde{I}_2}{K_{KI_2}+g_{I_2}\tilde{I}_2}+\lambda_{KI_{12}}g_{K_0}\tilde{K}_0\frac{g_{I_{12}}\tilde{I}_{12}}{K_{KI_{12}}+g_{I_{12}}\tilde{I}_{12}}\right)\frac{1}{1+g_{I_\beta}\tilde{I}_\beta/K_{KI_\beta}}\\
&\quad-d_{K_0}g_{K_0}\tilde{K}_0,\\
\frac{d(g_K\tilde{K})}{dt} &= \left(\lambda_{KI_2}g_{K_0}\tilde{K}_0\frac{g_{I_2}\tilde{I}_2}{K_{KI_2}+g_{I_2}\tilde{I}_2}+\lambda_{KI_{12}}g_{K_0}\tilde{K}_0\frac{g_{I_{12}}\tilde{I}_{12}}{K_{KI_{12}}+g_{I_{12}}\tilde{I}_{12}}\right)\frac{1}{1+g_{I_\beta}\tilde{I}_\beta/K_{KI_\beta}}-d_Kg_K\tilde{K},\\
g_{I_2}\tilde{I}_2 &= \frac{\lambda_{I_2T_8}}{d_{I_2}}g_{T_8}\tilde{T}_8,\\
g_{I_\gamma}\tilde{I}_\gamma &= \frac{\lambda_{I_\gamma K}}{d_{I_\gamma}}g_K\tilde{K},\\
g_{I_\alpha}\tilde{I}_\alpha &= \frac{1}{d_{I_\alpha}}\left[\lambda_{I_\alpha T_8}g_{T_8}\tilde{T}_8+\lambda_{I_\alpha K}g_K\tilde{K}\right],\\
g_{I_\beta}\tilde{I}_\beta &= \frac{1}{d_{I_\beta}}\left[\lambda_{I_\beta C}g_C\tilde{C}+\lambda_{I_\beta M_2}g_{M_2}\tilde{M}_2\right],
\end{aligned}
$$

$$
\begin{aligned}
g_{I_{10}}\tilde{I}_{10} &= \frac{\lambda_{I_{10}C}}{d_{I_{10}}} g_C \tilde{C},\\
g_{I_{12}}\tilde{I}_{12} &= \frac{\lambda_{I_{12}D}}{d_{I_{12}}} g_D \tilde{D},\\
\frac{d(g_{P_D^{T_8}}\tilde{P}_D^{T_8})}{dt} &= \lambda_{P_D^{T_8}} g_{T_8}\tilde{T}_8 - \lambda_{P_D A_1}\frac{d_{Q_A}}{\lambda_{Q_A}+d_{Q_A}} g_{P_D^{T_8}}\tilde{P}_D^{T_8} g_{A_1}\tilde{A}_1 - d_{P_D} g_{P_D^{T_8}}\tilde{P}_D^{T_8},\\
\frac{d(g_{P_D^{K}}\tilde{P}_D^{K})}{dt} &= \lambda_{P_D^{K}} g_{K}\tilde{K} - \lambda_{P_D A_1}\frac{d_{Q_A}}{\lambda_{Q_A}+d_{Q_A}} g_{P_D^{T_8}} g_{P_D^{K}}\tilde{P}_D^{K} g_{A_1}\tilde{A}_1 - d_{P_D} g_{P_D^{K}}\tilde{P}_D^{K},\\
g_{Q_A^{T_8}}\tilde{Q}_A^{T_8} &= \frac{\lambda_{P_D A_1}}{\lambda_{Q_A}+d_{Q_A}} g_{P_D^{T_8}}\tilde{P}_D^{T_8} g_{A_1}\tilde{A}_1,\\
g_{Q_A^{K}}\tilde{Q}_A^{T_8} &= \frac{\lambda_{P_D A_1}}{\lambda_{Q_A}+d_{Q_A}} g_{P_D^{K}}\tilde{P}_D^{K} g_{A_1}\tilde{A}_1,\\
\frac{d(g_{A_1}\tilde{A}_1)}{dt} &= \sum_{j=1}^{n}\xi_j f_{\text{pembro}}\delta(t-t_j) - \lambda_{P_D A_1}\frac{d_{Q_A}}{\lambda_{Q_A}+d_{Q_A}}(g_{P_D^{T_8}}\tilde{P}_D^{T_8} + g_{P_D^{K}}\tilde{P}_D^{K}) g_{A_1}\tilde{A}_1 - d_{A_1} g_{A_1}\tilde{A}_1,\\
\frac{d(g_{P_L}\tilde{P}_L)}{dt} &= \lambda_{P_L C} g_C\tilde{C} + \lambda_{P_L M_2} g_{M_2}\tilde{M}_2 - d_{P_L} g_{P_L}\tilde{P}_L,\\
g_{Q^{T_8}}\tilde{Q}^{T_8} &= \frac{\lambda_{P_D P_L}}{\lambda_Q} g_{P_D^{T_8}}\tilde{P}_D^{T_8} g_{P_L}\tilde{P}_L,\\
g_{Q^{K}}\tilde{Q}^{K} &= \frac{\lambda_{P_D P_L}}{\lambda_Q} g_{P_D^{K}}\tilde{P}_D^{K} g_{P_L}\tilde{P}_L,\\
\frac{d(g_{P_D^{8\text{LN}}}\tilde{P}_D^{8\text{LN}})}{dt} &= \lambda_{P_D^{8\text{LN}}} g_{T_A^8}\tilde{T}_A^8 - \lambda_{P_D A_1}\frac{d_{Q_A}}{\lambda_{Q_A}+d_{Q_A}} g_{P_D^{8\text{LN}}}\tilde{P}_D^{8\text{LN}} g_{A_1^{\text{LN}}}\tilde{A}_1^{\text{LN}} - d_{P_D} g_{P_D^{8\text{LN}}}\tilde{P}_D^{8\text{LN}},\\
g_{Q_A^{8\text{LN}}}\tilde{Q}_A^{8\text{LN}} &= \frac{\lambda_{P_D A_1}}{\lambda_{Q_A}+d_{Q_A}} g_{P_D^{8\text{LN}}}\tilde{P}_D^{8\text{LN}} g_{A_1}\tilde{A}_1,\\
\frac{d(g_{A_1^{\text{LN}}}\tilde{A}_1^{\text{LN}})}{dt} &= \sum_{j=1}^{n}\xi_j f_{\text{pembro}}\delta(t-t_j) - \lambda_{P_D A_1}\frac{d_{Q_A}}{\lambda_{Q_A}+d_{Q_A}} g_{P_D^{8\text{LN}}}\tilde{P}_D^{8\text{LN}} g_{A_1^{\text{LN}}}\tilde{A}_1^{\text{LN}} - d_{A_1} g_{A_1^{\text{LN}}}\tilde{A}_1^{\text{LN}},\\
\frac{d(g_{P_L^{\text{LN}}}\tilde{P}_L^{\text{LN}})}{dt} &= \lambda_{P_L^{\text{LN}} D^{\text{LN}}} g_{D^{\text{LN}}}\tilde{D}^{\text{LN}} - d_{P_L} g_{P_L^{\text{LN}}}\tilde{P}_L^{\text{LN}},\\
g_{Q^{8\text{LN}}}\tilde{Q}^{8\text{LN}} &= \frac{\lambda_{P_D P_L}}{\lambda_Q} g_{P_D^{8\text{LN}}}\tilde{P}_D^{8\text{LN}} g_{P_L^{\text{LN}}}\tilde{P}_L^{\text{LN}}.
\end{aligned}
$$

Simplifying and rearranging these equations leads to

$$
\begin{aligned}
\frac{d\tilde{C}}{dt} &= \lambda_C\tilde{C}\left(1-\frac{\tilde{C}}{C_0/g_C}\right) - \lambda_{CT_8} g_{T_8}\tilde{T}_8\frac{1}{1+\tilde{I}_\beta/(K_{CI_\beta}/g_{I_\beta})}\frac{1}{1+\tilde{Q}^{T_8}/(K_{CQ^{T_8}}/g_{Q^{T_8}})}\tilde{C}\\
&\quad - \lambda_{CK} g_K\tilde{K}\frac{1}{1+\tilde{I}_\beta/(K_{CI_\beta}/g_{I_\beta})}\frac{1}{1+\tilde{Q}^{K}/(K_{CQ^{K}}/g_{Q^{K}})}\tilde{C} - \lambda_{CI_\alpha}\frac{\tilde{I}_\alpha}{K_{CI_\alpha}/g_{I_\alpha}+\tilde{I}_\alpha}\tilde{C},\\
\frac{d\tilde{N}_c}{dt} &= \lambda_{CI_\alpha}\frac{g_C}{g_{N_c}}\frac{\tilde{I}_\alpha}{K_{CI_\alpha}/g_{I_\alpha}+\tilde{I}_\alpha}\tilde{C} - d_{N_c}\tilde{N}_c,\\
\frac{d\tilde{D}_0}{dt} &= \mathcal{A}_{D_0}/g_{D_0} - \lambda_{DN_c}\tilde{D}_0\frac{\tilde{N}_c}{K_{DN_c}/g_{N_c}+\tilde{N}_c} - \lambda_{D_0 K} g_K\tilde{D}_0\tilde{K}\frac{1}{1+\tilde{I}_\beta/(K_{D_0 I_\beta}/g_{I_\beta})} - d_{D_0}\tilde{D}_0,
\end{aligned}
$$

$$\frac{d\tilde{D}}{dt} = \lambda_{DN_c}\frac{g_{D_0}}{g_D}\tilde{D}_0\frac{\tilde{N}_c}{K_{DN_c}/g_{N_c}+\tilde{N}_c} - \lambda_{DD^{\mathrm{LN}}}\tilde{D} - d_D\tilde{D},$$

$$\frac{d\tilde{D}^{\mathrm{LN}}}{dt} = \frac{V_{\mathrm{TS}}}{V_{\mathrm{LN}}}\lambda_{DD^{\mathrm{LN}}}\frac{g_D}{g_{D^{\mathrm{LN}}}}e^{-d_D\tau_m}\tilde{D} - d_D\tilde{D}^{\mathrm{LN}},$$

$$\frac{d\tilde{T}_0^8}{dt} = \tilde{A}_{T_0^8} - \frac{\lambda_{T_0^8T_A^8}}{g_{T_0^8}}\tilde{R}^8 - d_{T_0^8}\tilde{T}_0^8,$$

where $\tilde{R}^8$ is defined as

$$\tilde{R}^8 := \frac{g_{D^{\mathrm{LN}}}g_{T_0^8}\tilde{D}^{\mathrm{LN}}\tilde{T}_0^8}{\left(1+\tilde{T}_A^r/(K_{T_0^8T_A^r}/g_{T_A^r})\right)\left(1+\tilde{Q}^{8\mathrm{LN}}/(K_{T_0^8Q^{8\mathrm{LN}}}/g_{Q^{8\mathrm{LN}}})\right)},$$

$$\tilde{T}_A^8 = \frac{2^{n_{\max}^8}e^{-d_{T_0^8}\tau_{T_A^8}}\frac{\lambda_{T_0^8T_A^8}}{g_{T_A^8}}\tilde{R}^8}{\left(\lambda_{T_A^8T_8}+d_{T_8}\right)\left(1+\tilde{T}_A^r/(K_{T_A^8T_A^r}/g_{T_A^r})\right)\left(1+\tilde{Q}^{8\mathrm{LN}}/(K_{T_A^8Q^{8\mathrm{LN}}}/g_{Q^{8\mathrm{LN}}})\right)},$$

$$\begin{aligned}\frac{d\tilde{T}_8}{dt} &= \frac{V_{\mathrm{LN}}}{V_{\mathrm{TS}}}\lambda_{T_A^8T_8}\frac{g_{T_A^8}}{g_{T_8}}e^{-d_{T_8}\tau_a}\tilde{T}_A^8 + \lambda_{T_8I_2}\frac{\tilde{T}_8\tilde{I}_2}{K_{T_8I_2}/g_{I_2}+\tilde{I}_2}\frac{1}{1+\tilde{T}_r/(K_{T_8T_r}/g_{T_r})}\\ &\quad - \lambda_{T_8C}\frac{\tilde{T}_8\tilde{C}}{K_{T_8C}/g_C+\tilde{C}} + \lambda_{T_{\mathrm{ex}}A_1}\frac{g_{T_{\mathrm{ex}}}}{g_{T_8}}\frac{\tilde{T}_{\mathrm{ex}}\tilde{A}_1}{K_{T_{\mathrm{ex}}A_1}/g_{A_1}+\tilde{A}_1} - \frac{d_{T_8}\tilde{T}_8}{1+\tilde{I}_{10}/\left(K_{T_8I_{10}}/g_{I_{10}}\right)},\end{aligned}$$

$$\frac{d\tilde{T}_{\mathrm{ex}}}{dt} = \lambda_{T_8C}\frac{g_{T_8}}{g_{T_{\mathrm{ex}}}}\frac{\tilde{T}_8\tilde{C}}{K_{T_8C}/g_C+\tilde{C}} - \lambda_{T_{\mathrm{ex}}A_1}\frac{\tilde{T}_{\mathrm{ex}}\tilde{A}_1}{K_{T_{\mathrm{ex}}A_1}/g_{A_1}+\tilde{A}_1} - \frac{d_{T_{\mathrm{ex}}}\tilde{T}_{\mathrm{ex}}}{1+\tilde{I}_{10}/\left(K_{T_{\mathrm{ex}}I_{10}}/g_{I_{10}}\right)},$$

$$\frac{d\tilde{T}_0^r}{dt} = \mathcal{A}_{T_0^r}/g_{T_0^r} - \lambda_{T_0^rT_A^r}/g_{T_0^r}\tilde{R}^r - d_{T_0^r}\tilde{T}_0^r,$$

where $\tilde{R}^r$ is defined as

$$\tilde{R}^r := g_{D^{\mathrm{LN}}}g_{T_0^r}\tilde{D}^{\mathrm{LN}}\tilde{T}_0^r,$$

$$\tilde{T}_A^r = \frac{2^{n_{\max}^r}e^{-d_{T_0^r}\tau_{T_A^r}}\frac{\lambda_{T_0^rT_A^r}}{g_{T_A^r}}\tilde{R}^r}{\lambda_{T_A^rT_r}+d_{T_r}},$$

$$\frac{d\tilde{T}_r}{dt} = \frac{V_{\mathrm{LN}}}{V_{\mathrm{TS}}}\lambda_{T_A^rT_r}\frac{g_{T_A^r}}{g_{T_r}}e^{-d_{T_r}\tau_a}\tilde{T}_A^r - d_{T_r}\tilde{T}_r,$$

$$\tilde{M}_0 = \frac{\mathcal{A}_{M_0}/g_{M_0}}{\lambda_{M_1I_\gamma}\frac{I_\gamma}{\left(K_{M_1I_\gamma}/g_{I_\gamma}\right)+I_\gamma} + \lambda_{M_1I_\alpha}\frac{I_\alpha}{\left(K_{M_1I_\alpha}/g_{I_\alpha}\right)+I_\alpha} + \lambda_{M_2I_\beta}\frac{I_\beta}{\left(K_{M_2I_\beta}/g_{I_\beta}\right)+I_\beta} + \lambda_{M_2I_{10}}\frac{I_{10}}{\left(K_{M_2I_{10}}/g_{I_{10}}\right)+I_{10}} + d_{M_0}},$$

$$\frac{d\tilde{M}_2}{dt} = \frac{\lambda_{M_2I_\beta}g_{M_0}}{g_{M_2}}\tilde{M}_0\frac{I_\beta}{\left(K_{M_2I_\beta}/g_{I_\beta}\right)+I_\beta} + \frac{\lambda_{M_2I_{10}}g_{M_0}}{g_{M_2}}\tilde{M}_0\frac{I_{10}}{\left(K_{M_2I_{10}}/g_{I_{10}}\right)+I_{10}} - d_{M_2}\tilde{M}_2,$$

$$\frac{d\tilde{K}_0}{dt} = \mathcal{A}_{K_0}/g_{K_0} - \left(\lambda_{KI_2}\tilde{K}_0\frac{\tilde{I}_2}{K_{KI_2}/g_{I_2}+\tilde{I}_2} + \lambda_{KI_{12}}\tilde{K}_0\frac{\tilde{I}_{12}}{K_{KI_{12}}/g_{I_{12}}+\tilde{I}_{12}}\right)\frac{1}{1+\tilde{I}_\beta/(K_{KI_\beta}/g_{I_\beta})} - d_{K_0}\tilde{K}_0,$$

$$\frac{d\tilde{K}}{dt} = \left(\lambda_{KI_2}\frac{g_{K_0}}{g_K}\tilde{K}_0\frac{\tilde{I}_2}{K_{KI_2}/g_{I_2}+\tilde{I}_2} + \lambda_{KI_{12}}\frac{g_{K_0}}{g_K}\tilde{K}_0\frac{\tilde{I}_{12}}{K_{KI_{12}}/g_{I_{12}}+\tilde{I}_{12}}\right)\frac{1}{1+\tilde{I}_\beta/(K_{KI_\beta}/g_{I_\beta})} - d_K\tilde{K},$$

$$
\begin{aligned}
\tilde{I}_2 &= \frac{\lambda_{I_2T_8} g_{T_8}/g_{I_2}}{d_{I_2}} \tilde{T}_8, \\
\tilde{I}_\gamma &= \frac{\lambda_{I_\gamma K} g_K/g_{I_\gamma}}{d_{I_\gamma}} \tilde{K}, \\
\tilde{I}_\alpha &= \frac{1}{d_{I_\alpha}} \left[ \lambda_{I_\alpha} \frac{g_{T_8}}{g_{I_\alpha}} \tilde{T}_8 + \lambda_{I_\alpha K} \frac{g_K}{g_{I_\alpha}} \tilde{K} \right], \\
\tilde{I}_\beta &= \frac{1}{d_{I_\beta}} \left[ \lambda_{I_\beta C} \frac{g_C}{g_{I_\beta}} \tilde{C} + \lambda_{I_\beta M_2} \frac{g_{M_2}}{g_{I_\beta}} \tilde{M}_2 \right], \\
\tilde{I}_{10} &= \frac{\lambda_{I_{10}C} g_C/g_{I_{10}}}{d_{I_{10}}} \tilde{C}, \\
\tilde{I}_{12} &= \frac{\lambda_{I_{12}D} g_D/g_{I_{12}}}{d_{I_{12}}} \tilde{D}, \\
\frac{d\tilde{P}_D^{T_8}}{dt} &= \lambda_{P_D^{T_8}} \frac{g_{T_8}}{g_{P_D^{T_8}}} \tilde{T}_8 - \lambda_{P_D A_1} \frac{d_{Q_A}}{\lambda_{Q_A} + d_{Q_A}} g_{P_D^{T_8}} g_{A_1} \tilde{P}_D^{T_8} \tilde{A}_1 - d_{P_D} \tilde{P}_D^{T_8}, \\
\frac{d\tilde{P}_D^{K}}{dt} &= \lambda_{P_D^{K}} \frac{g_{K}}{g_{P_D^{K}}} \tilde{K} - \lambda_{P_D A_1} \frac{d_{Q_A}}{\lambda_{Q_A} + d_{Q_A}} g_{P_D^{T_8}} g_{A_1} \tilde{P}_D^{K} \tilde{A}_1 - d_{P_D} \tilde{P}_D^{K}, \\
\tilde{Q}_A^{T_8} &= \frac{\lambda_{P_D A_1} g_{P_D^{T_8}} g_{A_1}/g_{Q_A^{T_8}}}{(\lambda_{Q_A} + d_{Q_A})} \tilde{P}_D^{T_8} \tilde{A}_1, \\
\tilde{Q}_A^{K} &= \frac{\lambda_{P_D A_1} g_{P_D^{K}} g_{A_1}/g_{Q_A^{K}}}{(\lambda_{Q_A} + d_{Q_A})} \tilde{P}_D^{K} \tilde{A}_1, \\
\frac{d\tilde{A}_1}{dt} &= \sum_{j=1}^{n} \xi_j \frac{f_{\text{pembro}}}{g_{A_1}} \delta(t - t_j) - \lambda_{P_D A_1} \frac{d_{Q_A}}{\lambda_{Q_A} + d_{Q_A}} (g_{P_D^{T_8}} \tilde{P}_D^{T_8} + g_{P_D^{K}} \tilde{P}_D^{K}) \tilde{A}_1 - d_{A_1} \tilde{A}_1, \\
\frac{d\tilde{P}_L}{dt} &= \lambda_{P_L C} \frac{g_C}{g_{P_L}} \tilde{C} + \lambda_{P_L M_2} \frac{g_{M_2}}{g_{P_L}} \tilde{M}_2 - d_{P_L} \tilde{P}_L, \\
\tilde{Q}^{T_8} &= \frac{\lambda_{P_D P_L} g_{P_D^{T_8}} g_{P_L}/g_{Q^{T_8}}}{\lambda_Q} \tilde{P}_D^{T_8} \tilde{P}_L, \\
\tilde{Q}^{K} &= \frac{\lambda_{P_D P_L} g_{P_D^{K}} g_{P_L}/g_{Q^{K}}}{\lambda_Q} \tilde{P}_D^{K} \tilde{P}_L, \\
\frac{d\tilde{P}_D^{8\text{LN}}}{dt} &= \lambda_{P_D^{8\text{LN}}} \frac{g_{T_A^8}}{g_{P_D^{8\text{LN}}}} \tilde{T}_A^8 - \lambda_{P_D A_1} \frac{d_{Q_A}}{\lambda_{Q_A} + d_{Q_A}} g_{P_D^{8\text{LN}}} g_{A_1^{\text{LN}}} \tilde{P}_D^{8\text{LN}} \tilde{P}_D^{8\text{LN}} \tilde{A}_1^{\text{LN}} - d_{P_D} \tilde{P}_D^{8\text{LN}}, \\
\tilde{Q}_A^{8\text{LN}} &= \frac{\lambda_{P_D A_1} g_{P_D^{8\text{LN}}} g_{A_1}/g_{Q_A^{8\text{LN}}}}{(\lambda_{Q_A} + d_{Q_A})} \tilde{P}_D^{8\text{LN}} \tilde{A}_1^{\text{LN}}, \\
\frac{d\tilde{A}_1^{\text{LN}}}{dt} &= \sum_{j=1}^{n} \xi_j \frac{f_{\text{pembro}}}{g_{A_1^{\text{LN}}}} \delta(t - t_j) - \lambda_{P_D A_1} \frac{d_{Q_A}}{\lambda_{Q_A} + d_{Q_A}} g_{P_D^{8\text{LN}}} g_{P_D^{8\text{LN}}} \tilde{P}_D^{8\text{LN}} \tilde{A}_1^{\text{LN}} - d_{A_1} \tilde{A}_1^{\text{LN}}, \\
\frac{d\tilde{P}_L^{\text{LN}}}{dt} &= \lambda_{P_L^{\text{LN}} D^{\text{LN}}} \frac{g_{D^{\text{LN}}}}{g_{P_L^{\text{LN}}}} \tilde{D}^{\text{LN}} - d_{P_L} \tilde{P}_L^{\text{LN}}, \\
\tilde{Q}^{8\text{LN}} &= \frac{\lambda_{P_D P_L} g_{P_D^{8\text{LN}}} g_{P_L^{\text{LN}}}/g_{Q^{8\text{LN}}}}{\lambda_Q} \tilde{P}_D^{8\text{LN}} \tilde{P}_L^{\text{LN}}.
\end{aligned}
$$

From this, the equivalence of the model to the equations in Section 2.4 is as stated in Section 2.6.

### B Parameter Estimation

We estimate all parameters, where possible, under the assumption that no pembrolizumab has/will be administered. The exception to this is the parameters directly related to pembrolizumab treatment, for which the assumptions are explicitly stated during estimation. Many of the assumptions and techniques in this section are adopted from [1].

#### B.1 Steady States and Initial Conditions for Cells in the TS

The steady states and initial conditions for all cells in the TS are as in [1], except for naive macrophages. The resultant steady states and initial conditions for the model, derived using ImmucellAI [2] and CIBERSORTx [3] or QSSA, are shown in Table B.1 and Table B.2, respectively. Justification for the choice of naive macrophage initial conditions is in Appendix B.9.2

| $C$ | $N_c$ | $D_0$ | $D$ | $T_8$ | $T_{\text{ex}}$ |
|---|---|---|---|---|---|
| $3.31 \times 10^7$ | $3.68 \times 10^6$ | $1.46 \times 10^6$ | $4.78 \times 10^5$ | $1.78 \times 10^5$ | $1.40 \times 10^5$ |
| $T_r$ | $M_0$ | $M_2$ | $K_0$ | $K$ | |
| $1.45 \times 10^5$ | $5.16 \times 10^5$ | $1.60 \times 10^6$ | $4.82 \times 10^5$ | $4.82 \times 10^6$ | |

Table B.1: TS steady-state cell densities for the model, combining estimates derived from ImmuCellAI and CIBERSORTx. All values are in cell/cm$^3$.

| $C$ | $N_c$ | $D_0$ | $D$ | $T_8$ | $T_{\text{ex}}$ |
|---|---|---|---|---|---|
| $1.79 \times 10^7$ | $1.99 \times 10^6$ | $1.63 \times 10^6$ | $8.29 \times 10^5$ | $2.43 \times 10^5$ | $2.09 \times 10^5$ |
| $T_r$ | $M_0$ | $M_2$ | $K_0$ | $K$ | |
| $2.12 \times 10^5$ | $5.58 \times 10^5$ | $1.23 \times 10^6$ | $3.06 \times 10^5$ | $5.20 \times 10^6$ | |

Table B.2: TS initial condition cell densities for the model, combining estimates derived from ImmuCellAI and CIBERSORTx, or QSSA. All values are in cell/cm$^3$.

We note that, technically, ImmuCellAI is an enrichment-based method that does not provide absolute immune cell proportions but rather estimates abundances across various immune cell subtypes not reported by CIBERSORTx. However, normalising these abundances provides a good approximation of the true immune cell proportions, thereby allowing ImmuCellAI to be justifiably employed to estimate immune cell steady states and initial conditions.

#### B.2 Steady States and Initial Conditions for Cells in the TDLN

The steady states and initial conditions for all cells in the TDLN are as in [1], except for effector CD8+ T cells and effector Tregs. The resultant steady states and initial conditions for the model, derived using ImmucellAI or QSSA, are shown in Table B.3 and Table B.4, respectively. Justification for the choice of effector CD8+ T cell and effector Treg initial conditions is in Appendix B.10.1 and Appendix B.10.2.

| $D^{\text{LN}}$ | $T_0^8$ | $T_A^8$ | $T_0^r$ | $T_A^r$ |
|---|---|---|---|---|
| $6.04 \times 10^6$ | $1.20 \times 10^7$ | $8.60 \times 10^5$ | $1.72 \times 10^5$ | $7.81 \times 10^5$ |

Table B.3: TDLN steady-state cell densities for the model, using estimates derived from ImmuCellAI. All values are in cell/cm$^3$.

| $D^{\mathrm{LN}}$ | $T_0^8$ | $T_A^8$ | $T_0^r$ | $T_A^r$ |
|---|---|---|---|---|
| $1.05 \times 10^7$ | $1.20 \times 10^7$ | $1.30 \times 10^6$ | $9.95 \times 10^4$ | $7.85 \times 10^5$ |

Table B.4: TDLN initial condition cell densities for the model, using estimates derived from ImmuCellAI or QSSA. All values are in cell/cm$^3$.

#### B.3 Steady States and Initial Conditions for Cytokines

To estimate cytokine steady states and initial conditions, we look at the respective experimental tissue concentration data, noting that 1 cm$^3$ = 1 mL for all cytokine measurements. We note that cytokines only appear in the model within an inhibition or half-saturation constant, making their absolute magnitude less important since they always appear as a ratio.

##### B.3.1 Estimates for IL-2 ($I_2$)

The tissue concentration of IL-2 in CRC is very low and was found to be below the lower limit of quantification in various experiments [4, 5]. In tumour supernatants of invasive ductal cancer, the median IL-2 concentration was found to be 2.1 pg/mL with the interquartile range being 2.0 pg/mL – 4.9 pg/mL [6]. We assume similar concentrations of IL-2 in the tissue of CRC patients.

Taking into account the well-documented anti-tumour properties of IL-2 [7, 8], we assume that $I_2$ has a steady-state value of $2.00 \times 10^{-12}$ g/cm$^3$. The initial condition for $I_2$ is justified in Appendix B.8.1.

##### B.3.2 Estimates for IFN-γ ($I_\gamma$)

It was found in [4] that the median tissue concentration of IFN-γ in CRC patients was 15.2 pg/mL, with the upper quartile concentration being approximately 16.9 pg/mL. It was found in [9] that the serum concentration of IFN-γ in stage IV CRC patients (median ≈ 20.75 pg/mL) is significantly higher than that of stage I-III patients (median ≈ 1 pg/mL). We thus set the steady state of $I_\gamma$ to $1.69 \times 10^{-11}$ g/cm$^3$. The initial condition for $I_\alpha$ is justified in Appendix B.8.2.

##### B.3.3 Estimates for TNF ($I_\alpha$)

It was found in [9] that in advanced CRC patients, i.e those with stage III or stage IV disease, the mean TNF tissue concentration was ≈ 53 pg/mL, with the concentration one standard deviation below the mean being approximately 16 pg/mL. Furthermore, the serum TNF concentration in stage IV CRC patients (median 20.3 pg/mL) is significantly higher than in stage III CRC patients (median 16.0 pg/mL) [10]. We thus set the steady state of $I_\alpha$ to be $5.30 \times 10^{-11}$ g/cm$^3$. The initial condition for $I_\alpha$ is justified in Appendix B.8.3.

##### B.3.4 Estimates for TGF-β ($I_\beta$)

It was found in [11] that in CRC patients, the mean TGF-β tissue concentration was 1311.5 pg/mg, with the concentration one standard error below the mean being 1153.9 pg/mg. Assuming a tissue density of 1.03 g/mL, these correspond to tissue concentrations of $1.19 \times 10^6$ pg/mL and $1.51 \times 10^6$ pg/mL, respectively. Taking into account the immunosuppressive nature of TGF-β, we set the steady state of $I_\beta$ to be $1.51 \times 10^{-6}$ g/cm$^3$. The initial condition for $I_\beta$ is justified in Appendix B.8.4.

##### B.3.5 Estimates for IL-10 ($I_{10}$)

It was found in [9] that in advanced CRC patients, i.e those with stage III or stage IV disease, the mean IL-10 tissue concentration was 115 pg/mL, with the concentration one standard deviation below the mean being approximately 46 pg/mL. Furthermore, the serum IL-10 concentration in stage IV CRC patients (mean 36.02 pg/mL) is significantly higher than in stage III CRC patients (mean 17.07 pg/mL) [12]. We thus set the steady state of $I_{10}$ to be $1.15 \times 10^{-10}$ g/cm$^3$. The initial condition for $I_{10}$ is justified in Appendix B.8.5.

##### B.3.6 Estimates for IL-12 ($I_{12}$)

The tissue concentration of IL-12 in CRC is very low, and was found to be below the lower limit of quantification in various experiments [4, 5]. In CRC cells in 3D spheroid cultures, the mean IL-12 concentration from cell supernatants was found to be $\approx 0.6$ pg/mL, with the concentration one standard deviation above the mean being $\approx 0.625$ pg/mL [13]. Furthermore, the serum IL-12 concentration in metastatic CRC patients is lower than in patients without distant metastasis [14]. We thus set the steady state of $I_{12}$ to be $6.00 \times 10^{-13}$ g/cm$^3$. The initial condition for $I_{12}$ is justified in Appendix B.8.6.

#### B.4 TDLN Parameters

##### B.4.1 Estimate for $V_{\text{TS}}$

The mean tumour volume in CRC patients with a T stage of T4a or an N stage of N2 was found to be 27.56 cm$^3$ and 27.57 cm$^3$, respectively [15]. As such, we set $V_{\text{TS}} = 2.76 \times 10^1$ cm$^3$.

##### B.4.2 Estimate for $V_{\text{LN}}$

The mean diameter of lymph nodes in CRC patients where cancer has metastasised was found to be 5.6 mm in [16]. Assuming a spherical lymph node, this corresponds to $V_{\text{LN}} = \frac{4}{3} \times 2.8^3 \times \pi$ mm$^3$ $= 9.20 \times 10^{-2}$ cm$^3$.

##### B.4.3 Estimate for $\tau_m$

In [17], it took 18 hours for DCs, which acquired antigen from a site of subcutaneous injection, to arrive at the lymph node. We assume that this migration time is the same for DCs acquiring cancer antigens from the TS so that $\tau_m = 18$ hr $= 0.75$ day.

##### B.4.4 Estimate for $\tau_a$

To estimate $\tau_a$, we note that T cells in the TDLN travel at speeds of $11 - 14$ $\mu$m/min, in comparison to DCs which migrate at speeds of $3 - 6$ $\mu$m/min [18]. We thus have that $\tau_a = \frac{4.5}{12.5}\tau_m \approx 0.27$ day.

##### B.4.5 Estimates for CD8+ T cells

Using data from [19], we estimate that naive CD8+ T cells take 2 days to activate, and so set $\tau_{\text{act}}^8 = 2$ day. It was found in [20] that activated CD8+ T cells required 39 hours on average to complete their first cell division, so we set $\Delta_8^0 = 39$ hr $= 1.63$ day. Furthermore, the average division time for subsequent cell cycles is 8.6 hours [20]; however, it can vary between $5 - 28$ hours. Thus, we set $\Delta_8 = 8.6$ hr $= 0.36$ day. It was shown in [21] that fully activated CD8+ T cells divide a minimum of $7 - 10$ times; however, they can divide more if persistent antigen exposure is present. Indeed, in

Lymphocytic Choriomeningitis Virus (LCV), CD8+ T cells can divide more than 15 times [22]. We perform a compromise and set $n^8_{\max} = 10$. We thus have that $\tau_{T^8_A} = 4.87$ day.

##### B.4.6 Estimates for Tregs

It takes between 12 and 24 hours for the first CD4+ T cell division to occur, with subsequent divisions occurring at a rate of approximately 10 hours per cell division [23]. We assume that the cell division rates of Tregs and CD4+ T helper cells are the same, so that $\Delta^0_r = 0.77$ day and $\Delta_r = 0.42$ day. We thus have that $\tau_{T^r_A} = 2.87$ day. It was found in [24] that in mice, 6 days after tumour implantation, 45% of Tregs in the TDLN had undergone at least 1 division, and 14% had undergone more than six divisions. We thus set $n^r_{\max} = 6$.

#### B.5 Half-Saturation Constants

We recall that for some species $X$, $K_X$ is denoted the half-saturation constant of $X$ in a term of the form

$$\frac{X}{K_X + X}.$$

For simplicity, we assume that if $\overline{X}$ denotes the steady-state value of $X$, then

$$\frac{\overline{X}}{K_X + \overline{X}} = \frac{1}{2} \implies K_X = \overline{X}. \tag{B.1}$$

This implies that

$$\begin{aligned}
K_{DN_c} &= \overline{N_c} = 3.68 \times 10^6 \text{ cell/cm}^3,\\
K_{T_8 I_2} = K_{K I_2} &= \overline{I_2} = 2.00 \times 10^{-12} \text{ g/cm}^3,\\
K_{T_8 C} &= \overline{C} = 3.31 \times 10^7 \ \left(\text{cell/cm}^3\right),\\
K_{M_1 I_\gamma} &= \overline{I_\gamma} = 1.69 \times 10^{-11} \text{ g/cm}^3,\\
K_{C I_\alpha} = K_{T_8 I_\alpha} = K_{M_1 I_\alpha} &= \overline{I_\alpha} = 5.30 \times 10^{-11} \text{ g/cm}^3,\\
K_{M_2 I_\beta} &= \overline{I_\beta} = 1.51 \times 10^{-6} \text{ g/cm}^3,\\
K_{M_2 I_{10}} &= \overline{I_{10}} = 1.15 \times 10^{-10} \text{ g/cm}^3,\\
K_{K I_{12}} &= \overline{I_{12}} = 6.00 \times 10^{-13} \text{ g/cm}^3.
\end{aligned}$$

To estimate $K_{T_{\text{ex}} A_1}$, we note that the value of the geometric mean $C_{\text{avg}}$ of pembrolizumab in serum at steady state varied minimally regardless of whether pembrolizumab was administered at 200 mg every 3 weeks, or 400 mg every 6 weeks [25]. This was equal to approximately 50.8 $\mu$g/mL, and we assumed this to be the same in tissue, so we take $C_{\text{avg}} = 5.08 \times 10^{-5}$ g/cm$^3$ $= 2.05 \times 10^{14}$ molec/cm$^3$, noting that the molecular mass of pembrolizumab is approximately 149,000 g/mol [26]. Thus, we assume that $K_{T_{\text{ex}} A_1} = 2.05 \times 10^{14}$ molec/cm$^3$.

#### B.6 Inhibition Constants

We recall that for some species $X$, $K_X$ is denoted as the inhibition constant of $X$ in a term of the form

$$\frac{1}{1 + X/K_X}.$$

For simplicity, we assume that if $\overline{X}$ denotes the steady-state value of $X$, then

$$\frac{1}{1+\overline{X}/K_X} = \frac{1}{2} \implies K_X = \overline{X}. \tag{B.2}$$

This implies that

$$\begin{aligned}
K_{T_8T_r} &= \overline{T_r} = 1.30 \times 10^5 \text{ cell/cm}^3,\\
K_{CI_\beta} = K_{D_0I_\beta} = K_{KI_\beta} &= \overline{I_\beta} = 1.51 \times 10^{-6} \text{ g/cm}^3,\\
K_{T_8I_{10}} = K_{T_{ex}I_{10}} &= \overline{I_{10}} = 1.15 \times 10^{-10} \text{ g/cm}^3,\\
K_{CQ^{T_8}} &= \overline{Q^{T_8}} = 6.67 \times 10^5 \text{ molec/cm}^3,\\
K_{CQ^K} &= \overline{Q^K} = 3.61 \times 10^6 \text{ molec/cm}^3,\\
K_{T_0^8T_A^r} = K_{T_A^8T_A^r} &= \overline{T_A^r} = 7.81 \times 10^5 \text{ cell/cm}^3,\\
K_{T_0^8Q^{8\text{LN}}} = K_{T_A^8Q^{8\text{LN}}} &= \overline{Q^{8\text{LN}}} = 5.40 \times 10^4 \text{ molec/cm}^3.
\end{aligned}$$

#### B.7 Degradation Rates

We recall the formula that the degradation rate of some species, $X$, is given by

$$d_X = \frac{\ln 2}{t_{1/2}^X} \tag{B.3}$$

where $t_{1/2}^X$ is the half-life of $X$.

##### B.7.1 Estimate for $d_{D_0}$

The time taken for immature DCs to degrade is estimated to be 28 days in mice [27]. We assume that this is similarly the case for humans, so that this corresponds to

$$d_{D_0} = \frac{1}{28 \text{ day}} = 3.57 \times 10^{-2} \text{ day}^{-1}.$$

##### B.7.2 Estimate for $d_D$

Mature DCs have a half-life of 1.5 – 2.9 days in mice [28]. We assume that this is similarly the case for humans, and take $t_{1/2}^D = 2.2$ day so that

$$d_D = \frac{\ln 2}{2.2 \text{ day}} = 3.15 \times 10^{-1} \text{ day}^{-1}.$$

##### B.7.3 Estimate for $d_{T_0^8}$

The half-life of naive CD8+ T cells in the lymph node was estimated to be 21.5 days in [29] so that

$$d_{T_0^8} = \frac{\ln 2}{21.5 \text{ day}} = 3.22 \times 10^{-2} \text{ day}^{-1}.$$

###### B.7.4 Estimate for $d_{T_8}$ and $d_{T_{\text{ex}}}$

It was measured in [30] that the mean death rate of circulating CD8+ T cells in HIV seronegative patients was 0.009 day$^{-1}$. We assume that this is the case for MSI-H/dMMR CRC, and so we set $d_{T_8} = d_{T_{\text{ex}}} = 0.009$ day$^{-1}$.

###### B.7.5 Estimate for $d_{T_0^r}$

The death rate of naive Tregs in the lymph node was estimated to be $2.2 \times 10^{-3}$ day$^{-1}$ in [31], and we assume that the death rate in MSI-H/dMMR CRC is similar, so that

$$d_{T_0^r} = 2.2 \times 10^{-3} \text{ day}^{-1}.$$

###### B.7.6 Estimate for $d_{T_r}$

The mean half-life of Tregs in healthy adults was measured to be approximately 11 days in [32]. We assume that this is similarly the case for MSI-H/dMMR CRC and that this corresponds to

$$d_{T_r} = \frac{\ln 2}{11 \text{ day}} = 6.30 \times 10^{-2} \text{ day}^{-1}.$$

###### B.7.7 Estimate for $d_{M_0}$

The lifespan for naive macrophages was found in humans to be approximately 1.37 days on average [33]. This corresponds to

$$d_{M_0} = \frac{1}{1.37 \text{ day}} = 0.73 \text{ day}^{-1}.$$

###### B.7.8 Estimate for $d_{M_2}$

The lifespan for M2 macrophages was found in humans to be approximately 7.41 days on average [33]. This corresponds to

$$d_{M_2} = \frac{1}{7.41 \text{ day}} = 1.35 \times 10^{-1} \text{ day}^{-1}.$$

###### B.7.9 Estimate for $d_{K_0}$ and $d_K$

The half-life of human NK cells varies between $1-2$ weeks [34–36]. We assume that the half-lives of resting and activated NK cells are both equal to 10 days, so that

$$d_{K_0} = d_K = \frac{\ln 2}{10 \text{ day}} = 6.93 \times 10^{-2} \text{ day}^{-1}.$$

###### B.7.10 Estimate for $d_{I_2}$

The half-life of IL-2 varies between $5-7$ minutes [37]. We take $t_{1/2}^{I_2} = 6.9$ min so that

$$d_{I_2} = \frac{\ln 2}{6.9 \text{ min}} = 1.45 \times 10^{2} \text{ day}^{-1}.$$

##### B.7.11 Estimate for $d_{I_\gamma}$

The half-life of IFN-γ varies between 25 – 35 minutes [38]. We take $t_{1/2}^{I_\gamma}$ to be 30 minutes so that

$$d_{I_\gamma} = \frac{\ln 2}{30 \text{ min}} = 3.33 \times 10^1 \text{ day}^{-1}.$$

##### B.7.12 Estimate for $d_{I_\alpha}$

The half-life of TNF varies between 15 – 30 minutes [39, 40]. We take $t_{1/2}^{I_\alpha}$ to be 18.2 minutes, so that

$$d_{I_\alpha} = \frac{\ln 2}{18.2 \text{ min}} = 5.48 \times 10^1 \text{ day}^{-1}.$$

##### B.7.13 Estimate for $d_{I_\beta}$

The half-life of active TGF-β is approximately 2 – 3 minutes [41]. We take $t_{1/2}^{I_\beta} = 2.5$ min, so that

$$d_{I_\beta} = \frac{\ln 2}{2.5 \text{ min}} = 3.99 \times 10^2 \text{ day}^{-1}.$$

##### B.7.14 Estimate for $d_{I_{10}}$

The half-life of IL-10 varies between 2.7 – 4.5 hours [42]. We take $t_{1/2}^{I_{10}} = 2.7$ hr so that

$$d_{I_{10}} = \frac{\ln 2}{2.7 \text{ hr}} = 6.16 \text{ day}^{-1}.$$

##### B.7.15 Estimate for $d_{I_{12}}$

The half-life of IL-12 varies between 5.3 – 10.3 hours [43]. We take $t_{1/2}^{I_{12}} = 7.8$ hr, so that

$$d_{I_{12}} = \frac{\ln 2}{7.8 \text{ hr}} = 2.13 \text{ day}^{-1}.$$

##### B.7.16 Estimate for $d_{P_D}$

The median lower bound on PD-1 half-life on human peripheral blood mononuclear cells was found to be 49.5 hours based on leucine enrichment in [44]. Hence, we take $t_{1/2}^{P_D} = 49.5$h so that

$$d_{P_D} = \frac{\ln 2}{49.5 \text{ hr}} = 3.36 \times 10^{-1} \text{ day}^{-1}.$$

##### B.7.17 Estimate for $d_{Q_A}$

The internalisation rate of the PD-1/pembrolizumab complex was estimated to be 0.43 day$^{-1}$ in [45], and so we estimate $d_{Q_A} = 0.43$ day$^{-1}$.

##### B.7.18 Estimate for $d_{A_1}$

The half-life of pembrolizumab varies between 22 – 27 days [46–48]. We take it to be 23.7 days in consistency with models from Li et al. and Ahamadi et al. [49–51] so that

$$d_{A_1} = \frac{\ln 2}{23.7 \text{ day}} = 2.92 \times 10^{-2} \text{ day}^{-1}.$$

##### B.7.19 Estimate for $d_{P_L}$

The half-life of fully glycosylated PD-L1 is approximately 12 hours [52], with PD-L1 on immune cells being heavily glycosylated [53]. Thus, we take $t_{1/2}^{P_L} = 12$ hr so that

$$d_{P_L} = \frac{\ln 2}{12 \text{ hr}} = 1.39 \text{ day}^{-1}.$$

#### B.8 Cytokine Production Parameters

To estimate many of the cytokine production constants, we consider (2.47) – (2.52) at steady state and use the data from [54]. For each immune cell, we assume that each cytokine's corresponding gene expression is proportional to its production rate by that cell.

##### B.8.1 Estimates for IL-2 ($I_2$)

Considering (2.21) at steady state, or equivalently considering (2.47), leads to

$$\lambda_{I_2 T_8}\overline{T_8} - d_{I_2}\overline{I_2} = 0 \implies \lambda_{I_2 T_8} = 1.63 \times 10^{-15} \ \text{(g/cell) day}^{-1}.$$

Consequently, considering (2.47), we have that

$$I_2(0) = \frac{\lambda_{I_2 T_8}}{d_{I_2}} T_8(0) = 2.73 \times 10^{-12} \ \text{g/cm}^3.$$

##### B.8.2 Estimates for IFN-$\gamma$ ($I_\gamma$)

Considering (2.22) at steady state, or equivalently considering (2.48), leads to

$$\lambda_{I_\gamma K}\overline{K} - d_{I_\gamma}\overline{I_\gamma} = 0 \implies \lambda_{I_\gamma K} = 1.17 \times 10^{-16} \ \text{(g/cell) day}^{-1}.$$

Consequently, considering (2.48), we have that

$$I_\gamma(0) = \frac{\lambda_{I_\gamma K}}{d_{I_\gamma}} K(0) = 1.83 \times 10^{-11} \ \text{g/cm}^3.$$

##### B.8.3 Estimates for TNF ($I_\alpha$)

Using values from [54] and considering (2.23) at steady state, or equivalently considering (2.23), leads to the equations

$$\frac{\lambda_{I_\alpha T_8}}{0.0654443776961264} = \frac{\lambda_{I_\alpha K}}{0.114108294134927},$$

and

$$\lambda_{I_\alpha T_8}\overline{T_8} + \lambda_{I_\alpha K}\overline{K} - d_{I_\alpha}\overline{I_\alpha} = 0.$$

Solving these simultaneously leads to

$$\lambda_{I_\alpha T_8} = 3.38 \times 10^{-16} \ \ (\text{g/cell})\,\text{day}^{-1},$$
$$\lambda_{I_\alpha K} = 5.90 \times 10^{-16} \ \ (\text{g/cell})\,\text{day}^{-1}.$$

Consequently, considering (2.49), we have that

$$I_\alpha(0) = \frac{1}{d_{I_\alpha}} \left(\lambda_{I_\alpha T_8} T_8(0) + \lambda_{I_\alpha K} K(0)\right) = 5.75 \times 10^{-11} \ \text{g/cm}^3.$$

##### B.8.4 Estimates for TGF-β ($I_\beta$)

Estimating the production constants for TGF-β is slightly more complicated than it is for other cytokines. We assume that the results for fibroblastic reticular cells in [55] translate directly to results for cancer-associated fibroblasts (CAFs), which are considered to be all fibroblasts found in the TME [55]. We assume that at steady state, CAFs produce twice as much TGF-β as cancer cells in the TME, and denote the production rate of TGF-β by CAFs as $\lambda_{I_\beta C_F}$. This, in conjunction with values from [54], and considering (2.24) at steady state, or equivalently considering (2.50), leads to the equations

$$\frac{\lambda_{I_\beta C_F}}{2} = \frac{\lambda_{I_\beta C}}{1},$$

and

$$\frac{\lambda_{I_\beta C_F}}{0.175283003265127} = \frac{\lambda_{I_\beta} M_2}{0.63070357154901},$$

and

$$\lambda_{I_\beta C}\overline{C} + \lambda_{I_\beta M_2}\overline{M_2} - d_{I_\beta}\overline{I_\beta} = 0.$$

Solving these simultaneously leads to

$$\lambda_{I_\beta C} = 1.35 \times 10^{-11} \ \ (\text{g/cell})\,\text{day}^{-1},$$
$$\lambda_{I_\beta M_2} = 9.72 \times 10^{-10} \ \ (\text{g/cell})\,\text{day}^{-1}.$$

Consequently, considering (2.50), we have that

$$I_\beta(0) = \frac{1}{d_{I_\beta}} \left(\lambda_{I_\beta C} C(0) + \lambda_{I_\beta M_2} M_2(0)\right) = 9.05 \times 10^{-7} \ \text{g/cm}^3.$$

##### B.8.5 Estimates for IL-10 ($I_{10}$)

Considering (2.25) at steady state, or equivalently considering (2.51), leads to

$$\lambda_{I_{10} C}\overline{C} - d_{I_{10}}\overline{C} = 0 \implies \lambda_{I_{10} C} = 2.14 \times 10^{-17} \ \ (\text{g/cell})\,\text{day}^{-1}.$$

Consequently, considering (2.51), we have that

$$I_{10}(0) = \frac{\lambda_{I_{10} C}}{d_{I_{10}}} C(0) = 6.22 \times 10^{-11} \ \text{g/cm}^3.$$

##### B.8.6 Estimates for IL-12 ($I_{12}$)

Cconsidering (2.26) at steady state, or equivalently considering (2.52), leads to

$$\lambda_{I_{12}D}\overline{D} - d_{I_{12}}\overline{I_{12}} = 0 \implies \lambda_{I_{12}D} = 2.67 \times 10^{-18} \ \text{(g/cell) day}^{-1}.$$

Consequently, considering (2.52), we have that

$$I_{12}(0) = \frac{\lambda_{I_{12}D}}{d_{I_{12}}} D(0) = 1.04 \times 10^{-12} \ \text{g/cm}^3.$$

#### B.9 Parameters for DCs, Macrophages, and NK Cells

##### B.9.1 Estimates for DCs ($D_0$ and $D$)

Adding (2.3) and (2.4) at steady state leads to

$$\mathcal{A}_{D_0} - \frac{\lambda_{D_0K}\overline{D_0K}}{2} - d_{D_0}\overline{D_0} - \lambda_{DD^{\text{LN}}}\overline{D} - d_D\overline{D} = 0.$$

In [56], it was also shown that the percentage of immature DCs that were lysed as a result of NK cells is roughly linear in the ratio of NK cells to immature DCs. When a 1:1 ratio of activated NK cells to immature DCs is present, after 24 hours, roughly 35.5% of immature DCs are lysed, whereas if a 5:1 ratio is present, 85.5% of immature DCs are lysed. At steady state, the ratio of NK cells to immature DCs is $\approx 2.39 : 1$, corresponding to an approximate 52.85% being lysed. However, if we consider only immature DC loss due to degradation, after 24 hours, only $1 - e^{-d_{D_0}} \approx 3.54\%$ is lost to it. Thus, we assume at steady state that

$$\frac{\lambda_{D_0K}\overline{D_0K}}{2 \times 0.5285} = \frac{d_{D_0}\overline{D_0}}{0.0354} \implies \lambda_{D_0K} = 2.21 \times 10^{-7} \ \left(\text{cell/cm}^3\right)^{-1} \text{day}^{-1}.$$

Considering (2.4) at steady state leads to

$$\frac{\lambda_{DN_c}\overline{D_0}}{2} - \lambda_{DD^{\text{LN}}}\overline{D} - d_D\overline{D} = 0.$$

Finally, it was found in [57] that only a limited number of DCs migrate up to the TDLN, with at most 4% of DCs reaching the TDLN in melanoma patients when DCs were injected intradermally. We assume at steady state that this holds, too, for MSI-H/dMMR CRC. Taking into account that only $e^{-d_D\tau_m}$ of mature DCs that leave the TS survive their migration to the TDLN, we have that

$$\frac{\lambda_{DD^{\text{LN}}}}{0.04e^{d_D\tau_m}} = \frac{d_D}{1 - 0.04e^{d_D\tau_m}}.$$

Solving these simultaneously leads to

$$\begin{aligned}
\mathcal{A}_{D_0} &= 9.89 \times 10^5 \ \left(\text{cell/cm}^3\right) \text{day}^{-1}, \\
\lambda_{DN_c} &= 2.17 \times 10^{-1} \ \text{day}^{-1}, \\
\lambda_{DD^{\text{LN}}} &= 1.68 \times 10^{-2} \ \text{day}^{-1}.
\end{aligned}$$

##### B.9.2 Estimates for Macrophages ($M_0$ and $M_2$)

To estimate the macrophage production constants, we consider (2.17) and (2.18) at steady state, and use the data from [54]. We assume that the magnitude of response to a specific cytokine is proportional to its corresponding macrophage polarisation rate, where the response is defined as the Euclidean distance between the centroid vectors of cytokine-treated macrophages and phosphate-buffered saline (PBS)-treated macrophages.

Considering (2.17) at steady state, or equivalently considering (2.46), leads to

$$\mathcal{A}_{M_0} - \frac{\lambda_{M_1 I_\gamma}\overline{M_0}}{2} - \frac{\lambda_{M_1 I_\alpha}\overline{M_0}}{2} - \frac{\lambda_{M_2 I_\beta}\overline{M_0}}{2} - \frac{\lambda_{M_2 I_{10}}\overline{M_0}}{2} - d_{M_0}\overline{M_0} = 0.$$

Additionally considering (2.18) at steady state leads to

$$\frac{\lambda_{M_2 I_\beta}\overline{M_0}}{2} + \frac{\lambda_{M_2 I_{10}}\overline{M_0}}{2} - d_{M_0}\overline{M_0} - d_{M_2}\overline{M_2} = 0.$$

Using values from [54] leads to

$$\frac{\lambda_{M_1 I_\gamma}}{2\times 12.54} = \frac{\lambda_{M_1 I_\alpha}}{2\times 10.77} = \frac{\lambda_{M_2 I_\beta}}{2\times 7.63} = \frac{\lambda_{M_2 I_{10}}}{2\times 6.81}.$$

Solving these simultaneously leads to

$$\begin{aligned}
\mathcal{A}_{M_0} &= 6.02\times 10^{6}\ \left(\text{cell/cm}^3\right)\text{day}^{-1},\\
\lambda_{M_1 I_\gamma} &= 7.27\times 10^{-1}\ \text{day}^{-1},\\
\lambda_{M_1 I_\alpha} &= 6.24\times 10^{-1}\ \text{day}^{-1},\\
\lambda_{M_2 I_\beta} &= 4.42\times 10^{-1}\ \text{day}^{-1},\\
\lambda_{M_2 I_{10}} &= 3.95\times 10^{-1}\ \text{day}^{-1}.
\end{aligned}$$

Consequently, considering (2.46), we have that

$$\begin{aligned}
M_0(0) &= \frac{\mathcal{A}_{M_0}}{\lambda_{M_1 I_\gamma}\frac{I_\gamma(0)}{K_{M_1 I_\gamma}+I_\gamma(0)} + \lambda_{M_1 I_\alpha}\frac{I_\alpha(0)}{K_{M_1 I_\alpha}+I_\alpha(0)} + \lambda_{M_2 I_\beta}\frac{I_\beta(0)}{K_{M_2 I_\beta}+I_\beta(0)} + \lambda_{M_2 I_{10}}\frac{I_{10(0)}}{K_{M_2 I_{10}}+I_{10(0)}} + d_{M_0}}\\
&= 5.58\times 10^{5}\ \text{cell/cm}^3.
\end{aligned}$$

##### B.9.3 Estimates for NK Cells ($K_0$ and $K$)

Adding (2.19) and (2.20) at steady state leads to

$$\mathcal{A}_{K_0} - d_{K_0}\overline{K_0} - d_K\overline{K} = 0 \implies \mathcal{A}_{K_0} = 3.67\times 10^{5}\ \left(\text{cell/cm}^3\right)\text{day}^{-1}.$$

Considering (2.20) at steady state leads to

$$\frac{1}{2}\left(\frac{\lambda_{K I_2}\overline{K_0}}{2} + \frac{\lambda_{K I_{12}}\overline{K_0}}{2}\right) - d_K\overline{K} = 0.$$

Furthermore, we use data from [54] and assume that the magnitude of response to a specific cytokine is proportional to its corresponding NK cell activation rate, where the response is defined as the Euclidean distance between the centroid vectors of cytokine-treated NK cells and PBS-treated NK cells. Thus,

$$\frac{\lambda_{KI_2}/2}{15.4} = \frac{\lambda_{KI_{12}}/2}{14.83}.$$

Solving these simultaneously leads to

$$\lambda_{KI_2} = 1.41\ \text{day}^{-1},$$
$$\lambda_{KI_{12}} = 1.36\ \text{day}^{-1}.$$

#### B.10 T Cell Parameters and Estimates

##### B.10.1 Estimates for CD8+ T Cells ($T_0^8$, $T_A^8$, $T_8$, and $T_{\text{ex}}$)

Considering (2.6) at steady state leads to

$$\mathcal{A}_{T_0^8} - \overline{R^8} - d_{T_0^8}\overline{T_0^8} = 0,$$

and in particular,

$$\overline{R^8} = \frac{\lambda_{T_0^8 T_A^8}\overline{D^{\text{LN}}}\,\overline{T_0^8}}{4}.$$

Considering (2.9) at steady state, or equivalently considering (2.44), leads to

$$\frac{2^{n_{\max}^8} e^{-d_{T_0^8}\tau_{T_A^8}}\overline{R^8}}{4} - \lambda_{T_A^8 T_8}\overline{T_A^8} - d_{T_8}\overline{T_A^8} = 0.$$

We first consider the case where no pembrolizumab is present. Considering (2.10) and (2.11) at steady state leads to

$$\frac{V_{\text{LN}}}{V_{\text{TS}}}\lambda_{T_A^8 T_8} e^{-d_{T_8}\tau_a}\overline{T_A^8} + \frac{\lambda_{T_8 I_2}\overline{T_8}}{4} - \frac{\lambda_{T_8 C}\overline{T_8}}{2} - \frac{d_{T_8}\overline{T_8}}{2} = 0,$$
$$\frac{\lambda_{T_8 C}\overline{T_8}}{2} - \frac{d_{T_{\text{ex}}}\overline{T_{\text{ex}}}}{2} = 0.$$

We assume that at steady state, 95% of positive $T_8$ growth is due to $T_A^8$ migration to the TS, and the other 5% is due to IL-2-induced proliferation. Thus, we have that

$$\frac{V_{\text{LN}}}{V_{\text{TS}}}\frac{\lambda_{T_A^8 T_8} e^{-d_{T_8}\tau_a}\overline{T_A^8}}{0.95} = \frac{\lambda_{T_8 I_2}\overline{T_8}/4}{0.05}.$$

To determine $\lambda_{T_{\text{ex}}A_1}$, we assume that when pembrolizumab is present, at steady state, 20% of exhausted CD8+ T cells are reinvigorated. That is, we assume that

$$\frac{\lambda_{T_{\text{ex}}A_1}\overline{T_{\text{ex}}}/2}{0.2} = \frac{d_{T_{\text{ex}}}\overline{T_{\text{ex}}}/2}{0.8}.$$

Solving these equations simultaneously leads to

$$\mathcal{A}_{T_0^8} = 3.88 \times 10^5\ \left(\text{cell/cm}^3\right)\text{day}^{-1},$$

$$\lambda_{T_0^8 T_A^8} = 1.05 \times 10^{-10} \ \left(\text{cell/cm}^3\right)^{-1} \text{day}^{-1},$$
$$\overline{R^8} = 1.90 \times 10^3 \ \left(\text{cell/cm}^3\right) \text{day}^{-1},$$
$$\lambda_{T_A^8 T_8} = 4.75 \times 10^{-1} \ \text{day}^{-1},$$
$$\lambda_{T_8 I_2} = 1.61 \times 10^{-3} \ \text{day}^{-1},$$
$$\lambda_{T_8 C} = 7.08 \times 10^{-2} \ \text{day}^{-1},$$
$$\lambda_{T_{\text{ex}} A_1} = 2.25 \times 10^{-3} \ \text{day}^{-1}.$$

Consequently, considering (2.44), we have that

$$T_A^8(0) = \frac{2^{n_{\max}^8} e^{-d_{T_0^8}\tau_{T_A^8}} \lambda_{T_0^8 T_A^8} D^{\text{LN}}(0) T_0^8(0)}{\left(\lambda_{T_A^8 T_8} + d_{T_8}\right)\left(1 + T_A^r(0)/K_{T_0^8 T_A^r}\right)\left(1 + Q^{8\text{LN}}(0)/K_{T_0^8 Q^{8\text{LN}}}\right)\left(1 + T_A^r(0)/K_{T_A^8 T_A^r}\right)\left(1 + Q^{8\text{LN}}(0)/K_{T_A^8 Q^{8\text{LN}}}\right)}$$
$$= 1.30 \times 10^6 \ \text{cell/cm}^3,$$

where $T_A^r(0)$ is estimated in Appendix B.10.2.

##### B.10.2 Estimates for Tregs ($T_0^r$, $T_A^r$, and $T_r$)

Considering (2.12) at steady state leads to

$$\mathcal{A}_{T_0^r} - \overline{R^r} - d_{T_0^r}\overline{T_0^r} = 0,$$

where

$$\overline{R^r} = \lambda_{T_0^r T_A^r} \overline{D^{\text{LN}}}\,\overline{T_0^r}.$$

Considering (2.15) at steady state leads to

$$2^{n_{\max}^r} e^{-d_{T_0^r}\tau_{T_A^r}} \overline{R^r} - \lambda_{T_A^r T_r}\overline{T_A^r} - d_{T_r}\overline{T_A^r} = 0.$$

Finally, considering (2.16) at steady state leads to

$$\frac{V_{\text{LN}}}{V_{\text{TS}}} \lambda_{T_A^r T_r}\overline{T_A^r} - d_{T_r}\overline{T_r} = 0.$$

Solving these equations simultaneously leads to

$$\mathcal{A}_{T_0^r} = 4.42 \times 10^4 \ \left(\text{cell/cm}^3\right) \text{day}^{-1},$$
$$\lambda_{T_0^r T_A^r} = 4.22 \times 10^{-8} \ \left(\text{cell/cm}^3\right)^{-1} \text{day}^{-1},$$
$$\overline{R^r} = 4.39 \times 10^4 \ \left(\text{cell/cm}^3\right) \text{day}^{-1},$$
$$\lambda_{T_A^r T_r} = 3.51 \ \text{day}^{-1}.$$

Consequently, considering (2.45), we have that

$$T_A^r(0) = \frac{2^{n_{\max}^r} e^{-d_{T_0^r}\tau_{T_A^r}} \lambda_{T_0^r T_A^r}}{\lambda_{T_A^r T_r} + d_{T_r}} D^{\text{LN}}(0) T_0^r(0) = 7.85 \times 10^5 \ \text{cell/cm}^3.$$

#### B.11 Cancer Cell Parameters

Considering (2.1) at steady state leads to

$$\lambda_C \left(1 - \frac{\overline{C}}{C_0}\right) - \frac{\lambda_{CT_8}}{4}\overline{T_8} - \frac{\lambda_{CK}}{4}\overline{K} - \frac{\lambda_{CI_\alpha}}{2} = 0.$$

Considering (2.2) at steady state leads to the equation

$$\frac{\lambda_{CI_\alpha}\overline{C}}{2} - d_{N_c}\overline{N_c} = 0.$$

We assume that CD8+ T cells and NK cells kill cancer cells with similar potency, so we approximate

$$\lambda_{CK}/4 = \lambda_{CT_8}/4 \implies \lambda_{CK} = \lambda_{CT_8}.$$

Using values from [1], we assume that

$$\begin{aligned}\lambda_C &= 1.77 \times 10^{-1} \text{ day}^{-1},\\ C_0 &= 8.15 \times 10^{7} \text{ cell/cm}^3,\\ d_{N_c} &= 6.55 \times 10^{-1} \text{ day}^{-1}.\end{aligned}$$

Substituting these in, and solving these simultaneously, leads to

$$\begin{aligned}\lambda_{CT_8} &= 2.58 \times 10^{-8} \left(\text{cell/cm}^3\right)^{-1}\text{day}^{-1},\\ \lambda_{CK} &= 2.58 \times 10^{-8} \left(\text{cell/cm}^3\right)^{-1}\text{day}^{-1},\\ \lambda_{CI_\alpha} &= 1.46 \times 10^{-1} \text{ day}^{-1}.\end{aligned}$$

#### B.12 Estimates for Immune Checkpoint-Associated Components in the TS

##### B.12.1 Estimate for $\lambda_Q$

The dissociation rate of the PD-1/PD-L1 complex was found to be 1.44 $\text{sec}^{-1}$ in [58]. Thus, we have that

$$\lambda_Q = 60 \times 60 \times 24 \times 1.44 \text{ sec}^{-1} = 1.24 \times 10^{5} \text{ day}^{-1}.$$

##### B.12.2 Estimate for $\lambda_{P_D P_L}$

The formation rate of the PD-1/PD-L1 complex was found to be $1.84 \times 10^5$ $\text{M}^{-1}\text{sec}^{-1}$ in [58]. To convert this to units of $(\text{molec/cm}^3)^{-1}\text{day}^{-1}$, we recall that 1 M $=$ 1 mol/L $= 10^{-3}$ $\text{mol/cm}^3$ $= 6.022 \times 10^{20}$ $\text{molec/cm}^3$. As such,

$$\lambda_{P_D P_L} = 60 \times 60 \times 24 \times 1.84 \times 10^5 \times (6.022 \times 10^{20})^{-1} = 2.64 \times 10^{-11} \text{ (molec/cm}^3)^{-1}\text{day}^{-1}.$$

##### B.12.3 Estimates for Synthesis Rates and Steady States

We first denote $\rho_{P_D^{T_8}}$ and $\rho_{P_D^K}$ as the number of PD-1 molecules expressed on the surface of CD8+ T cells and activated NK cells in the TS, respectively. To determine these parameters, we used the baseline data collected in [59] on 5 advanced cancer patients before their pembrolizumab infusions. The net number of PD-1 molecules on the surface of CD8+ T cells was 2761 molec/cell, and so we set

$\rho_{P_D^{T_8}} = 2.76 \times 10^3$ molec/cell. Despite the net number of PD-1 molecules on the surface of NK cells being below the lower limit of quantification in [59], NK cells substantially express PD-1 [60] in CRC, and so we set $\rho_{P_D^K} = \rho_{P_D^{T_8}}/5 = 5.52 \times 10^2$ molec/cell.

We next denote $\rho_{P_L C}$ and $\rho_{P_L M_2}$ as the number of PD-L1 molecules expressed on cancer cells and M2 macrophages, respectively. In their quantitative systems pharmacology model of colorectal cancer, Anbari et al. estimated the baseline numbers of PD-L1 molecules per cancer cell and per APC to be $180,000$ molec/cell and $266,666$ molec/cell, respectively [61]. This makes sense, noting that PD-L1 expression in macrophages is stronger and more continuous than that in cancer cells [62]. As such, we set $\rho_{P_L C} = 1.8 \times 10^5$ molec/cell and $\rho_{P_L M_2} = 2.67 \times 10^5$ molec/cell.

Considering (2.35) - (2.38) at steady state in the absence of pembrolizumab leads to

$$\begin{aligned}
\lambda_{P_D^{T_8}} \overline{T_8} - d_{P_D} \overline{P_D^{T_8}} &= 0, \\
\lambda_{P_D^K} \overline{K} - d_{P_D} \overline{P_D^K} &= 0, \\
\lambda_{P_L C} \overline{C} + \lambda_{P_L M_2} \overline{M_2} - d_{P_L} \overline{P_L} &= 0, \\
\overline{Q^{T_8}} - \frac{\lambda_{P_D P_L}}{\lambda_Q} \overline{P_D^{T_8}}\,\overline{P_L} &= 0, \\
\overline{Q^K} - \frac{\lambda_{P_D P_L}}{\lambda_Q} \overline{P_D^K}\,\overline{P_L} &= 0.
\end{aligned}$$

By considering the total number of PD-1 receptors expressed on each PD-1-expressing cell at steady state, we expect in the absence of pembrolizumab that

$$\begin{aligned}
\overline{P_D^{T_8}} + \overline{Q^{T_8}} &= \rho_{P_D^{T_8}} \overline{T_8}, \\
\overline{P_D^K} + \overline{Q^K} &= \rho_{P_D^K} \overline{K}.
\end{aligned}$$

We can also consider the total number of PD-L1 ligands at steady state so that

$$\overline{P_L} + \overline{Q^{T_8}} + \overline{Q^K} = \rho_{P_L C} \overline{C} + \rho_{P_L M_2} \overline{M_2}.$$

Finally, we expect the synthesis rates of PD-1 and PD-L1 to be proportional to the total number of PD-1 molecules expressed per PD-1- and PD-L1-expressing cell, so that

$$\begin{aligned}
\frac{\lambda_{P_D^{T_8}}}{\rho_{P_D^{T_8}}} &= \frac{\lambda_{P_D^K}}{\rho_{P_D^K}}, \\
\frac{\lambda_{P_L C}}{\rho_{P_L C}} &= \frac{\lambda_{P_L M_2}}{\rho_{P_L M_2}}.
\end{aligned}$$

Solving these simultaneously and ensuring all model parameters are positive leads to

$$\begin{aligned}
\lambda_{P_D^{T_8}} &= 9.26 \times 10^2 \ \text{(molec/cell) day}^{-1}, \\
\lambda_{P_D^K} &= 1.85 \times 10^2 \ \text{(molec/cell) day}^{-1}, \\
\lambda_{P_L C} &= 2.50 \times 10^5 \ \text{(molec/cell) day}^{-1},
\end{aligned}$$

$$\lambda_{P_L M_2} = 3.71 \times 10^5 \ \text{(molec/cell) day}^{-1}.$$

This leads to

$$\begin{aligned}
\overline{P_D^{T_8}} &= 4.91 \times 10^8 \text{ molec/cm}^3, \\
\overline{P_D^{K}} &= 2.66 \times 10^9 \text{ molec/cm}^3, \\
\overline{P_L} &= 6.39 \times 10^{12} \text{ molec/cm}^3, \\
\overline{Q^{T_8}} &= 6.67 \times 10^5 \text{ molec/cm}^3, \\
\overline{Q^{K}} &= 3.61 \times 10^6 \text{ molec/cm}^3.
\end{aligned}$$

##### B.12.4 Estimates for Initial Conditions

To determine the relevant initial conditions, we can simply consider the total number of PD-1 receptors on each PD-1-expressing cell and PD-L1 ligands in the absence of pembrolizumab, so that

$$\begin{aligned}
P_D^{T_8}(0) + Q^{T_8}(0) &= \rho_{P_D^{T_8}} T_8(0), \\
P_D^{K}(0) + Q^{K}(0) &= \rho_{P_D^{K}} K(0), \\
P_L(0) + Q^{T_8}(0) + Q^{K}(0) &= \rho_{P_L C} C(0) + \rho_{P_L M_2} M_2(0).
\end{aligned}$$

We can also consider (2.34) – (2.35) initially, so that

$$\begin{aligned}
Q^{T_8}(0) - \frac{\lambda_{P_D P_L}}{\lambda_Q} P_D^{T_8}(0) P_L(0) &= 0, \\
Q^{K}(0) - \frac{\lambda_{P_D P_L}}{\lambda_Q} P_D^{K}(0) P_L(0) &= 0.
\end{aligned}$$

Solving these simultaneously leads to

$$\begin{aligned}
P_D^{T_8}(0) &= 6.70 \times 10^8 \text{ molec/cm}^3, \\
P_D^{K}(0) &= 2.87 \times 10^9 \text{ molec/cm}^3, \\
P_L(0) &= 3.55 \times 10^{12} \text{ molec/cm}^3, \\
Q^{T_8}(0) &= 5.06 \times 10^5 \text{ molec/cm}^3, \\
Q^{K}(0) &= 2.17 \times 10^6 \text{ molec/cm}^3.
\end{aligned}$$

We note that excluding bound PD-1 receptors when considering the total number of PD-1 receptors on PD-1-expressing cells does not affect the parameter estimates, steady states, or initial conditions at this level of precision, since the number of unbound PD-1 receptors is several orders of magnitude larger than the number of bound PD-1 receptors on PD-1-expressing cells. Furthermore, this also applies when considering the total number of PD-L1 ligands.

#### B.13 Estimates for Immune Checkpoint-Associated Components in the TDLN

##### B.13.1 Estimates for Synthesis Rates and Steady States

For simplicity, we assume that the total number of PD-1 receptors on cells in the TDLN is equal to the number on the corresponding cells in the TS. Thus, denoting $\rho_{P_D^{8\mathrm{LN}}}$ as the number of PD-1 molecules expressed on the surface of CD8+ T cells in the TDLN, we have that $\rho_{P_D^{8\mathrm{LN}}} = \rho_{P_D^{T_8}}$.

We denote $\rho_{P_L^{\mathrm{LN}}D^{\mathrm{LN}}}$ as the number of PD-L1 molecules expressed on mature DCs in the TDLN. It was found in [58] that the PD-L1 expression on mature DCs was $80,372$ molec/cell. However, amongst advanced CRC patients, only 22% of colonic DCs were PD-L1+ in [63]. We thus assumed that $\rho_{P_L^{\mathrm{LN}}D^{\mathrm{LN}}} = 1.77 \times 10^4$ molec/cell.

The procedure for estimating parameters, steady states, and initial conditions for PD-1, PD-L1, and the PD-1/PD-L1 complex in the TDLN is the same as in the TS. Considering (2.39) and (2.42) – (2.43) at steady state in the absence of pembrolizumab, and making the same assumptions for estimation as in the TS, we obtain

$$\begin{aligned}
\lambda_{P_D^{8\mathrm{LN}}}\overline{P_D^{8\mathrm{LN}}} - d_{P_D}\overline{P_D^{8\mathrm{LN}}} &= 0,\\
\lambda_{P_L^{\mathrm{LN}}D^{\mathrm{LN}}}\overline{D^{\mathrm{LN}}} - d_{P_L}\overline{P_L^{\mathrm{LN}}} &= 0,\\
\overline{Q^{8\mathrm{LN}}} - \frac{\lambda_{P_D P_L}}{\lambda_Q}\overline{P_D^{8\mathrm{LN}}P_L^{\mathrm{LN}}} &= 0,\\
\overline{P_D^{8\mathrm{LN}}} + \overline{Q^{8\mathrm{LN}}} &= \rho_{P_D^{8\mathrm{LN}}}\overline{T_A^8},\\
\overline{P_L^{\mathrm{LN}}} + \overline{Q^{8\mathrm{LN}}} &= \rho_{P_L^{\mathrm{LN}}D^{\mathrm{LN}}}\overline{D^{\mathrm{LN}}},
\end{aligned}$$

Solving these simultaneously and ensuring all model parameters are positive leads to

$$\begin{aligned}
\lambda_{P_D^{8\mathrm{LN}}} &= 9.27 \times 10^2 \ \ (\mathrm{molec/cell})\,\mathrm{day}^{-1},\\
\lambda_{P_L^{\mathrm{LN}}D^{\mathrm{LN}}} &= 2.46 \times 10^4 \ \ (\mathrm{molec/cell})\,\mathrm{day}^{-1},\\
\lambda_{P_L^{\mathrm{LN}}T_A^1} &= 2.89 \times 10^3 \ \ (\mathrm{molec/cell})\,\mathrm{day}^{-1}.
\end{aligned}$$

This leads to

$$\begin{aligned}
\overline{P_D^{8\mathrm{LN}}} &= 2.37 \times 10^9 \ \mathrm{molec/cm}^3,\\
\overline{P_L^{\mathrm{LN}}} &= 1.07 \times 10^{11} \ \mathrm{molec/cm}^3,\\
\overline{Q^{8\mathrm{LN}}} &= 5.40 \times 10^4 \ \mathrm{molec/cm}^3.
\end{aligned}$$

##### B.13.2 Estimates for Initial Conditions

To determine the relevant immune checkpoint initial conditions, we can simply consider the total number of PD-1 receptors on each PD-1-expressing cell and PD-L1 ligands in the absence of pembrolizumab, so that

$$P_D^{8\mathrm{LN}}(0) + Q^{8\mathrm{LN}}(0) = \rho_{P_D^{8\mathrm{LN}}}T_A^8(0),$$

$$P_L^{\mathrm{LN}}(0) + Q^{8\mathrm{LN}}(0) = \rho_{P_L^{\mathrm{LN}} D^{\mathrm{LN}}} D^{\mathrm{LN}}(0).$$

We can also consider (2.43) initially, so that

$$Q^{8\mathrm{LN}}(0) - \frac{\lambda_{P_D P_L}}{\lambda_Q} P_D^{8\mathrm{LN}}(0) P_L^{\mathrm{LN}}(0) = 0.$$

Solving these simultaneously leads to

$$\begin{aligned} P_D^{8\mathrm{LN}}(0) &= 1.56 \times 10^9 \text{ molec/cm}^3, \\ P_L^{\mathrm{LN}}(0) &= 1.86 \times 10^{11} \text{ molec/cm}^3, \\ Q^{8\mathrm{LN}}(0) &= 6.18 \times 10^4 \text{ molec/cm}^3. \end{aligned}$$

We note again that excluding bound PD-1 receptors when considering the total number of PD-1 receptors on PD-1-expressing cells does not affect the parameter estimates, steady states, or initial conditions at this level of precision, since the number of unbound PD-1 receptors is several orders of magnitude larger than the number of bound PD-1 receptors on PD-1-expressing cells. Furthermore, this also applies when considering the total number of PD-L1 ligands.

#### B.14 Estimates for $A_1$ and $A_1^{\mathrm{LN}}$

##### B.14.1 Estimate for $f_{\mathrm{pembro}}$

To determine $f_{\mathrm{pembro}}$, we use the formula

$$f_{\mathrm{pembro}} = \frac{C_{\mathrm{max,ss}}(\xi_{\mathrm{pembro}}) - C_{\mathrm{min,ss}}(\xi_{\mathrm{pembro}})}{\xi_{\mathrm{pembro}}}, \tag{B.4}$$

where $C_{\mathrm{max,ss}}/C_{\mathrm{min,ss}}$ corresponds to the maximum and minimum serum concentration of pembrolizumab at steady state after a dose, $\xi_{\mathrm{pembro}}$, of pembrolizumab is administered, respectively.

For pembrolizumab, the mean $C_{\mathrm{min,ss}}/C_{\mathrm{max,ss}}$ was found to be approximately 32.6/85.8 $\mu$g/mL with triweekly 200 mg pembrolizumab treatment [25]. This results in $f_{\mathrm{pembro}} \approx 2.90 \times 10^{-7}$ $(\mathrm{g/cm}^3)$/mg of pembrolizumab administered for all doses. To convert this into units of $(\mathrm{molec/cm}^3)$/mg, we note that the molecular mass of pembrolizumab is approximately 149,000 g/mol [26], which corresponds to $f_{\mathrm{pembro}} \approx 1.17 \times 10^{12}$ $(\mathrm{molec/cm}^3)$/mg.

#### B.15 Estimates for PD-1/pembrolizumab Complex on Cells

##### B.15.1 Estimate for $\lambda_{Q_A}$

The dissociation rate of the PD-1/pembrolizumab complex was measured using biolayer interferometry to be 2.6 day$^{-1}$ in [45]. Thus, we take $\lambda_{Q_A} = 2.6$ day$^{-1}$.

##### B.15.2 Estimate for $\lambda_{P_D A_1}$

We use the value of $\lambda_{P_D A_1}$ from [1], so we assume that

$$\lambda_{P_D A_1} = 4.69 \times 10^{-13} \left(\mathrm{molec/cm}^3\right)^{-1} \mathrm{day}^{-1}.$$

#### B.16 Model Parameters

Table B.5: Parameter values for the model. TS denotes the tumour site. est. denotes estimated parameters.

| Parameter | Description | Value | Unit | Referencessupp |
|---|---|---|---|---|
| $f_{\text{pembro}}$ | $A_1/A_1^{\text{LN}}$ dose scaling factor | $1.17\times10^{12}$ | $(\text{molec/cm}^3)/\text{mg}$ | est. |
| $\mathcal{A}_{D_0}$ | Source of $D_0$ | $9.89\times10^{5}$ | $(\text{cell/cm}^3)\,\text{day}^{-1}$ | est. |
| $\mathcal{A}_{T_0^8}$ | Source of $T_0^8$ | $3.88\times10^{5}$ | $(\text{cell/cm}^3)\,\text{day}^{-1}$ | est. |
| $\mathcal{A}_{T_0^r}$ | Source of $T_0^r$ | $4.42\times10^{4}$ | $(\text{cell/cm}^3)\,\text{day}^{-1}$ | est. |
| $\mathcal{A}_{M_0}$ | Source of $M_0$ | $6.02\times10^{6}$ | $(\text{cell/cm}^3)\,\text{day}^{-1}$ | est. |
| $\mathcal{A}_{K_0}$ | Source of $K_0$ | $3.67\times10^{5}$ | $(\text{cell/cm}^3)\,\text{day}^{-1}$ | est. |
| $\lambda_C$ | Growth rate of $C$ | $1.77\times10^{-1}$ | $\text{day}^{-1}$ | [1] |
| $\lambda_{CT_8}$ | Elimination rate of $C$ by $T_8$ | $2.58\times10^{-8}$ | $(\text{cell/cm}^3)^{-1}\,\text{day}^{-1}$ | est. |
| $\lambda_{CK}$ | Elimination rate of $C$ by $K$ | $2.58\times10^{-8}$ | $(\text{cell/cm}^3)^{-1}\,\text{day}^{-1}$ | est. |
| $\lambda_{CI_\alpha}$ | Necrosis rate of $C$ by $I_\alpha$ | $1.46\times10^{-1}$ | $\text{day}^{-1}$ | est. |
| $\lambda_{DN_c}$ | Maturation rate of $D_0$ by $N_c$ | $2.17\times10^{-1}$ | $\text{day}^{-1}$ | est. |
| $\lambda_{D_0K}$ | Killing rate of $D_0$ by $K$ | $2.21\times10^{-7}$ | $(\text{cell/cm}^3)^{-1}\,\text{day}^{-1}$ | est. |
| $\lambda_{DD^{\text{LN}}}$ | Migration rate of $D$ to TDLN | $1.68\times10^{-2}$ | $\text{day}^{-1}$ | est. |
| $\lambda_{T_0^8T_A^8}$ | Activation rate of $T_0^8$ to $T_A^8$ | $1.05\times10^{-10}$ | $(\text{cell/cm}^3)^{-1}\,\text{day}^{-1}$ | est. |
| $\lambda_{T_A^8T_8}$ | Migration rate of $T_A^8$ to the TS | $4.75\times10^{-1}$ | $\text{day}^{-1}$ | est. |
| $\lambda_{T_8I_2}$ | Growth rate of $T_8$ by $I_2$ | $1.61\times10^{-3}$ | $\text{day}^{-1}$ | est. |
| $\lambda_{T_8C}$ | Exhaustion rate of $T_8$ by $C$ | $7.08\times10^{-3}$ | $\text{day}^{-1}$ | est. |
| $\lambda_{T_{\text{ex}}A_1}$ | Reinvigoration rate of $T_{\text{ex}}$ | $2.25\times10^{-3}$ | $\text{day}^{-1}$ | est. |
| $\lambda_{T_0^rT_A^r}$ | Activation rate of $T_0^r$ to $T_A^r$ | $4.22\times10^{-8}$ | $(\text{cell/cm}^3)^{-1}\,\text{day}^{-1}$ | est. |
| $\lambda_{T_A^rT_r}$ | Migration rate of $T_A^r$ to the TS | 3.51 | $\text{day}^{-1}$ | est. |
| $\lambda_{M_1I_\gamma}$ | Polarisation rate of $M_0$ to $M_1$ by $I_\gamma$ | 7.27 | $\text{day}^{-1}$ | est. |
| $\lambda_{M_1I_\alpha}$ | Polarisation rate of $M_0$ to $M_1$ by $I_\alpha$ | 6.24 | $\text{day}^{-1}$ | est. |
| $\lambda_{M_2I_\beta}$ | Polarisation rate of $M_0$ to $M_2$ by $I_\beta$ | 4.42 | $\text{day}^{-1}$ | est. |
| $\lambda_{M_2I_{10}}$ | Polarisation rate of $M_0$ to $M_2$ by $I_{10}$ | 3.95 | $\text{day}^{-1}$ | est. |
| $\lambda_{KI_2}$ | Maturation rate of $K_0$ by $I_2$ | 1.41 | $\text{day}^{-1}$ | est. |
| $\lambda_{KI_{12}}$ | Maturation rate of $K_0$ by $I_{12}$ | 1.36 | $\text{day}^{-1}$ | est. |
| $\lambda_{I_2T_8}$ | Production rate of $I_2$ by $T_8$ | $1.63\times10^{-15}$ | $(\text{g/cell})\,\text{day}^{-1}$ | est. |
| $\lambda_{I_\gamma K}$ | Production rate of $I_\gamma$ by $K$ | $1.17\times10^{-16}$ | $(\text{g/cell})\,\text{day}^{-1}$ | est. |
| $\lambda_{I_\alpha T_8}$ | Production rate of $I_\alpha$ by $T_8$ | $3.38\times10^{-16}$ | $(\text{g/cell})\,\text{day}^{-1}$ | est. |
| $\lambda_{I_\alpha K}$ | Production rate of $I_\alpha$ by $K$ | $5.90\times10^{-16}$ | $(\text{g/cell})\,\text{day}^{-1}$ | est. |
| $\lambda_{I_\beta C}$ | Production rate of $I_\beta$ by $C$ | $1.35\times10^{-11}$ | $(\text{g/cell})\,\text{day}^{-1}$ | est. |
| $\lambda_{I_\beta M_2}$ | Production rate of $I_\beta$ by $M_2$ | $9.72\times10^{-11}$ | $(\text{g/cell})\,\text{day}^{-1}$ | est. |
| $\lambda_{I_{10}C}$ | Production rate of $I_{10}$ by $C$ | $2.14\times10^{-17}$ | $(\text{g/cell})\,\text{day}^{-1}$ | est. |
| $\lambda_{I_{12}D}$ | Production rate of $I_{12}$ by $D$ | $2.67\times10^{-18}$ | $(\text{g/cell})\,\text{day}^{-1}$ | est. |
| $\lambda_{P_D^{T_8}}$ | Synthesis rate of $P_D^{T_8}$ | $9.26\times10^{2}$ | $(\text{molec/cell})\,\text{day}^{-1}$ | est. |
| $\lambda_{Q_A}$ | Dissociation rate of the PD-1/pembrolizumab complex | 2.6 | $\text{day}^{-1}$ | [45] |

| | | | | |
|---|---|---|---|---|
| $\lambda_Q$ | Dissociation rate of the PD-1/PD-L1 complex | $1.24 \times 10^5$ | $\text{day}^{-1}$ | [58] |
| $\lambda_{P_D A_1}$ | Formation rate of the PD-1/pembrolizumab complex | $4.69 \times 10^{-13}$ | $(\text{molec/cm}^3)^{-1}\,\text{day}^{-1}$ | [1] |
| $\lambda_{P_D P_L}$ | Formation rate of the PD-1/PD-L1 complex | $2.64 \times 10^{-11}$ | $(\text{molec/cm}^3)^{-1}\,\text{day}^{-1}$ | [58] |
| $\lambda_{P_D^K}$ | Synthesis rate of $P_D^K$ | $1.85 \times 10^2$ | $(\text{molec/cell})\,\text{day}^{-1}$ | est. |
| $\lambda_{P_L C}$ | Synthesis rate of $P_L$ by $C$ | $2.50 \times 10^5$ | $(\text{molec/cell})\,\text{day}^{-1}$ | est. |
| $\lambda_{P_L M_2}$ | Synthesis rate of $P_L$ by $M_2$ | $3.71 \times 10^5$ | $(\text{molec/cell})\,\text{day}^{-1}$ | est. |
| $\lambda_{P_D^{8\text{LN}}}$ | Synthesis rate of $P_D^{8\text{LN}}$ | $9.27 \times 10^2$ | $(\text{molec/cell})\,\text{day}^{-1}$ | est. |
| $\lambda_{P_L^{\text{LN}} D^{\text{LN}}}$ | Synthesis rate of $P_L^{\text{LN}}$ by $D^{\text{LN}}$ | $2.46 \times 10^4$ | $(\text{molec/cell})\,\text{day}^{-1}$ | est. |
| $K_{CI_\alpha}$ | Half-saturation constant of $I_\alpha$ for $C$ | $5.30 \times 10^{-11}$ | $\text{g/cm}^3$ | est. |
| $K_{DN_c}$ | Half-saturation constant of $N_c$ for $D$ | $3.68 \times 10^6$ | $\text{cell/cm}^3$ | est. |
| $K_{T_8 I_2}$ | Half-saturation constant of $I_2$ for $T_8$ | $2.00 \times 10^{-12}$ | $\text{g/cm}^3$ | est. |
| $K_{T_8 C}$ | Half-saturation constant $T_8$ exhaustion due to $C$ exposure | $3.31 \times 10^7$ | $(\text{cell/cm}^3)\ \text{day}$ | est. |
| $K_{T_{\text{ex}} A_1}$ | Half-saturation constant of $T_{\text{ex}}$ reinvigoration by $A_1$ | $2.05 \times 10^{14}$ | $\text{molec/cm}^3$ | est. |
| $K_{M_1 I_\gamma}$ | Half-saturation constant of $I_\gamma$ for $M_1$ | $1.69 \times 10^{-11}$ | $\text{g/cm}^3$ | est. |
| $K_{M_1 I_\alpha}$ | Half-saturation constant of $I_\alpha$ for $M_1$ | $5.30 \times 10^{-11}$ | $\text{g/cm}^3$ | est. |
| $K_{M_2 I_\beta}$ | Half-saturation constant of $I_\beta$ for $M_2$ | $1.51 \times 10^{-6}$ | $\text{g/cm}^3$ | est. |
| $K_{M_2 I_{10}}$ | Half-saturation constant of $I_{10}$ for $M_2$ | $1.15 \times 10^{-10}$ | $\text{g/cm}^3$ | est. |
| $K_{KI_2}$ | Half-saturation constant of $I_2$ for $K$ | $2.00 \times 10^{-12}$ | $\text{g/cm}^3$ | est. |
| $K_{KI_{12}}$ | Half-saturation constant of $I_{12}$ for $K$ | $6.00 \times 10^{-13}$ | $\text{g/cm}^3$ | est. |
| $C_0$ | Carrying capacity of $C$ | $8.15 \times 10^7$ | $\text{cell/cm}^3$ | [1] |
| $K_{CI_\beta}$ | Inhibition constant of $T_8$ and $K$ elimination of $C$ by $I_\beta$ | $1.51 \times 10^{-6}$ | $\text{g/cm}^3$ | est. |
| $K_{CQ^{T_8}}$ | Inhibition constant of $T_8$ elimination of $C$ by $Q^{T_8}$ | $6.67 \times 10^5$ | $\text{molec/cm}^3$ | est. |
| $K_{CQ^K}$ | Inhibition constant of $K$ elimination of $C$ by $Q^K$ | $3.61 \times 10^6$ | $\text{molec/cm}^3$ | est. |
| $K_{D_0 I_\beta}$ | Inhibition constant of $K$ elimination of $D_0$ by $I_\beta$ | $1.51 \times 10^{-6}$ | $\text{g/cm}^3$ | est. |
| $V_{\text{TS}}$ | Volume of the TS | $2.76 \times 10^1$ | $\text{cm}^3$ | [15] est. |
| $V_{\text{LN}}$ | Volume of the TDLN | $9.20 \times 10^{-2}$ | $\text{cm}^3$ | [16] est. |
| $K_{T_0^8 T_A^r}$ | Inhibition constant of $T_0^8$ activation by $T_A^r$ | $7.81 \times 10^5$ | $\text{cell/cm}^3$ | est. |

| | | | | |
|---|---|---|---|---|
| $K_{T_0^8 Q^{8\text{LN}}}$ | Inhibition constant of $T_0^8$ activation by $Q^{8\text{LN}}$ | $5.40 \times 10^4$ | molec/cm$^3$ | est. |
| $K_{T_A^8 T_A^r}$ | Inhibition constant of $T_A^8$ activation by $T_A^r$ | $7.81 \times 10^5$ | cell/cm$^3$ | est. |
| $K_{T_A^8 Q^{8\text{LN}}}$ | Inhibition constant of $T_A^8$ proliferation by $Q^{8\text{LN}}$ | $5.40 \times 10^4$ | molec/cm$^3$ | est. |
| $K_{T_8 T_r}$ | Inhibition constant of $I_2$-mediated growth of $T_8$ by $T_r$ | $1.45 \times 10^5$ | cell/cm$^3$ | est. |
| $K_{T_8 I_{10}}$ | Inhibition constant of $T_8$ death by $I_{10}$ | $1.15 \times 10^{-10}$ | g/cm$^3$ | est. |
| $K_{T_{\text{ex}} I_{10}}$ | Inhibition constant of $T_{\text{ex}}$ death by $I_{10}$ | $1.15 \times 10^{-10}$ | g/cm$^3$ | est. |
| $K_{K I_\beta}$ | Inhibition constant of NK cell activation by $I_\beta$ | $1.51 \times 10^{-6}$ | g/cm$^3$ | est. |
| $d_{N_c}$ | Removal rate of $N_c$ | $6.55 \times 10^{-1}$ | day$^{-1}$ | est. |
| $d_{D_0}$ | Death rate of $D_0$ | $3.57 \times 10^{-2}$ | day$^{-1}$ | [27] est. |
| $d_D$ | Death rate of $D$ | $3.15 \times 10^{-1}$ | day$^{-1}$ | [28] est. |
| $d_{T_0^8}$ | Death rate of $T_0^8$ | $3.22 \times 10^{-2}$ | day$^{-1}$ | [29] est. |
| $d_{T_8}$ | Death rate of $T_8$ | $9 \times 10^{-3}$ | day$^{-1}$ | [30] |
| $d_{T_{\text{ex}}}$ | Death rate of $T_{\text{ex}}$ | $9 \times 10^{-3}$ | day$^{-1}$ | [30] |
| $d_{T_0^r}$ | Death rate of $T_0^r$ | $2.2 \times 10^{-3}$ | day$^{-1}$ | [31] |
| $d_{T_r}$ | Death rate of $T_r$ | $6.30 \times 10^{-2}$ | day$^{-1}$ | [32] est. |
| $d_{M_0}$ | Death rate of $M_0$ | 0.73 | day$^{-1}$ | [33] |
| $d_{M_2}$ | Death rate of $M_2$ | $1.35 \times 10^{-1}$ | day$^{-1}$ | [33] |
| $d_{K_0}$ | Death rate of $K_0$ | $6.93 \times 10^{-2}$ | day$^{-1}$ | [34–36] est. |
| $d_K$ | Death rate of $K$ | $6.93 \times 10^{-2}$ | day$^{-1}$ | [34–36] est. |
| $d_{I_2}$ | Degradation rate of $I_2$ | $1.45 \times 10^2$ | day$^{-1}$ | [37] est. |
| $d_{I_\gamma}$ | Degradation rate of $I_\gamma$ | $3.33 \times 10^1$ | day$^{-1}$ | [38] est. |
| $d_{I_\alpha}$ | Degradation rate of $I_\alpha$ | $5.48 \times 10^1$ | day$^{-1}$ | [39, 40] est. |
| $d_{I_\beta}$ | Degradation rate of $I_\beta$ | $3.99 \times 10^2$ | day$^{-1}$ | [41] est. |
| $d_{I_{10}}$ | Degradation rate of $I_{10}$ | 6.16 | day$^{-1}$ | [42] est. |
| $d_{I_{12}}$ | Degradation rate of $I_{12}$ | 2.13 | day$^{-1}$ | [43] est. |
| $d_{P_D}$ | Degradation rate of unbound PD-1 receptors | $3.36 \times 10^{-1}$ | day$^{-1}$ | [44] |
| $d_{Q_A}$ | Internalisation rate of the PD-1/pembrolizumab complex | 0.43 | day$^{-1}$ | [45] |
| $d_{A_1}$ | Elimination rate of $A_1/A_1^{\text{LN}}$ | $2.92 \times 10^{-2}$ | day$^{-1}$ | [49–51] est. |
| $d_{P_L}$ | Degradation rate of unbound PD-L1 | 1.39 | day$^{-1}$ | [52] |
| $\tau_m$ | DC migration time from TDLN to the TS | 0.75 | day | [17] est. |
| $\Delta_8^0$ | Time taken for first CTL division | 1.63 | day | [20] |
| $n_{\max}^8$ | Maximal number of CTL divisions in the TDLN | 10 | dimensionless | [21, 22] est. |
| $\Delta_8$ | Time taken for successive CTL divisions | 0.36 | day | [21] |

| | | | | |
|---|---|---|---|---|
| $\tau_{T_A^8}$ | Time taken for CTL division program | 4.87 | day | est. |
| $\tau_a$ | T cell migration time between the TDLN to the TS | 0.27 | day | est. |
| $\Delta_r^0$ | Time taken for first Treg division | 0.77 | day | [23] est. |
| $n_{\max}^r$ | Maximal number of Treg divisions in the TDLN | 6 | dimensionless | [24] est. |
| $\Delta_r$ | Time taken for successive Treg divisions | 0.42 | day | [23] est. |
| $\tau_{T_A^r}$ | Time taken for Treg division program | 2.87 | day | est. |
